# The secrets to domestic bliss – Partner fidelity and environmental filtering preserve stage-specific turtle ant gut symbioses for over 40 million years

**DOI:** 10.1101/2021.09.13.460005

**Authors:** Yi Hu, Catherine L. D’Amelio, Benoît Béchade, Christian S. Cabuslay, Jon G. Sanders, Shauna Price, Emily Fanwick, Scott Powell, Corrie S. Moreau, Jacob A. Russell

## Abstract

**Background:** Gut microbiomes can vary across development, a pattern often found for insects with complete metamorphosis. With varying nutritional need and distinct opportunities for microbial acquisition, questions arise as to how such ‘holometabolous’ insects retain helpful microbes at larval and adult stages. Ants are an intriguing system for such study. In a number of lineages adults digest only liquid food sources, while larvae digest solid foods. Like some other social insects, workers and soldiers of some ant species engage in oral-anal trophallaxes, enabling microbial transfer among siblings. But do queens, the typical colony founding caste, obtain symbionts through such transfer? Does this enable transgenerational symbiont passage? And does the resulting partner fidelity promote the evolution of beneficial symbionts? Furthermore, how might such adult-centric biology shape larval microbiomes? To address these questions, we characterized symbiotic gut bacteria across 13 species of *Cephalotes* turtle ants, with up to 40-million years of divergence. Adding to the prior focus on workers we, here, study underexplored castes and stages including queens, soldiers, and larvae, by performing 16S rRNA qPCR, amplicon sequencing, and phylogenetic classification.

**Results:** We show that adult microbiomes are conserved across species and largely across castes. Nearly 95% of the bacteria in adults have, thus far, been found only in *Cephalotes* ants. Furthermore, the microbiomes from most adults exhibit phylosymbiosis, a trend in which microbiome community similarity recapitulates patterns of host relatedness. Additionally, an abundant, adult-enriched symbiont cospeciates with some *Cephalotes*. Evidence here suggests that these partner fidelity patterns extend from transgenerational symbiont transfer through alate gyne dispersal and subsequent colony-founding by queens. Like adults, larvae of *Cephalotes* species exhibit strong microbiome conservation. Phylosymbiosis patterns are weaker, however, with further evidence elevating environmental filtering as a primary mechanism behind such conservation. Specifically, while adult-enriched symbionts are found in most larvae, symbionts of older larvae are highly related to free-living bacteria from the Enterobacteriaceae, Lactobacillales, and Actinobacteria.

**Conclusions:** Our findings suggest that both partner fidelity and conserved environmental filtering drive stable, stage-specific, social insect symbioses. We discuss the implications for our broader understanding of insect microbiomes, and the means of sustaining a beneficial microbiome.

## Background

Symbioses are a major feature of the ecology and evolution of many animals, shaping digestion, nutrient acquisition, defense, and dietary detoxification [1–9]. A persistent challenge for animals and other eukaryotes hosting symbiotic microbes is the need to avoid acquisition of harmful microbes, or the spread of cheaters among populations of mutualistic symbionts. Brakes to these threats can be provided by a series of mechanisms encompassed by partner fidelity, partner choice, and host-imposed sanctions [10–12].

Insects have provided useful insights into these mechanisms and their roles in sustaining beneficial symbioses [13]. Partner fidelity is the most widely documented, being evidenced by the many symbionts with transovarial transmission (e.g. Koga et al. [14]; Luan et al. [15]) and those passaged through fidelity-promoting “out-of-body” mechanisms, like egg smearing or egg capsule deposition [16]. But a range of insects acquire symbionts from the environment, retaining a subset of the inoculated microbes [17–20] due to their physiology and behavior [21, 22], or to competitive actions of their regular symbiont partners ( e.g. Worsley et al. [23]; Itoh et al. [24]). Among the diverse range of hexapods with environmental symbiont acquisition, some have evolved highly selective partner choice mechanisms, retaining a specific, beneficial subset of their starting inoculum due to anatomical and physiological innovations [25, 26].

While insects harbor symbiotic microbes in and on a variety of tissues and surfaces, their gut chambers – i.e. the crop, midgut, ileum, and rectum – comprise a common set of habitats for colonization [21]. Eukaryotic hosts are often touted as stable environments for microbial symbionts, but gut-inhabiting symbionts may encounter unstable conditions due to varying diet, the introduction of competitors, and the impacts of insect development [21, 27, 28]. Promoting shifts in microbiomes across life stages for other organisms [29, 30], the behavioral, morphological, and physiological changes unfolding across development have especially pronounced effects on gut symbiosis in holometabolous insects. This stems, partially, from gut tissue re-organization at metamorphosis [31] and to ejection of symbionts with the elimination of the meconium [32], necessitating separate symbiont acquisitions in larvae and adults. The resulting potential for stage-specific microbiomes is enhanced by use of differing diets across larval and adult stages, for many holometabolous insects. Several studies have, accordingly, reported changes in species composition or in the abundances of shared species on the opposite sides of metamorphosis [33–36], with some intriguing exceptions (e.g. Estes et al. [37]).

Despite these findings and widening research interest in holometabolous insect symbiosis, questions persist as to how often extracellular gut symbionts persist across metamorphosis (e.g. Coon et al. [38]), whether symbionts lost at pupation are reacquired in adulthood, and how hosts might promote acquisition of helpful microbes during different inoculation stages. Lineages of eusocial, holometabolous insects including bees and ants have evolved solutions to this latter challenge through trophallactic symbiont transmission [39–41] and symbiont persistence within the founders of new colonies [42–45]. Such fidelity-promoting mechanisms have favored a range of beneficial digestive, nutritional, and defensive services [4, 46, 47], yielding tightly intertwined partner histories. The partially congruent host and symbiont phylogenies within holometabolous corbiculate bees reveal a long term consequence of such fidelity; such cospeciation may also be prevalent among eusocial, hemimetabolous termites [40, 48–50].

Within the former holometabolous taxon, extensive research on honeybee microbiomes has revealed that specialized symbionts are enriched within adults [40], and that symbiotic communities may differ among castes within a single colony [51–53]. Similar work among the ants – distant, eusocial cousins of corbiculate bees – remains in its early-stages (e.g. Rubin et al. [54]; Segers et al. [55]). With the ants numbering in excess of 12,000 species, and weighing in to comprise the world’s most abundant insect group, our lack of knowledge on the impacts of development, physiology, and social behaviors on the ants’ symbiotic gut microbes represents a substantial shortcoming. In addition, given the often unique contributions of particular ant castes and life stages to the function [56, 57] and founding of ant colonies [58, 59], studies of microbiome composition among adults and larvae within a single colony hold promise in the broader effort to understand whether these social insects employ safeguards promoting mutualistic symbiont services, and the means by which symbionts shape the complex nature of colony-level fitness [55, 60].

To address these uncertainties, we focus here on the genus *Cephalotes*. Adult workers of this arboreal, New World ant clade harbor a conserved microbiome of extracellular gut bacteria that have lived with these ants and those from the sister genus, *Procryptocerus* (two genera forming the “cephalotines”, formerly tribe Cephalotini and now a junior synonym of tribe Attini [61]), since their descent from common ancestor over 50 million years ago [7, 62–64]. There is little evidence that most microbes found in adult workers live anywhere else (e.g. Russell et al. [65]; Anderson et al. [66]), with this high degree of specialization being thought to partially extend from social transmission. Indeed, mature adults from these two genera appear to passage their symbionts to newly eclosed adults, fresh-removed from pupation, using oral-anal trophallaxis [67–69]. At this stage, symbionts are sealed into the midgut, ileum, and rectal chambers through development of a fine filter over the proventriculus, an organ separating the crop from these downstream gut chambers [41]. Such specialized behavior and morphology are thought to favor the persistence of bacterial gut symbionts within the colony, while protecting adult microbiomes from invasion by outside microbes. This may be key to the long-term maintenance of beneficial symbioses, evidenced by symbiont contributions to nitrogen-recycling, amino acid provisioning, and the building of the *Cephalotes* cuticle, and by the widespread conservation of genes enabling these capacities across the genus [7, 70].

Symbiont movement across colonies – and generations – has been less studied, however, among the cephalotines, And, indeed, such transgenerational transfer may be key to the evolution of beneficial symbioses [10]. In *Cephalotes*, colonies are established in a suitable arboreal nesting cavity by one or more winged queens, after mating flights. During the early stages of colony life, queens will lose their wings, laying fertilized eggs, and act as the sole caretakers of first-born offspring. Given the expectation of symbiont persistence in these colony foundresses, young queens may also provide trophallactic inoculation, seeding their initial offspring with gut microbiomes at the larval and/or adult stage. Studies of gut microbes in such reproductive females during their pre- and post-colony founding states could, hence, help test the hypothesis that queens’ acquisition of symbionts from their parent colony, and subsequent symbiont persistence, promote the observed patterns of long-term partner fidelity [63].

As for many ants, we are unaware of whether cephalotine larvae harbor dense symbiotic gut communities, and we are similarly ignorant about the origins of any such symbionts. Might specialized bacteria from adults dominate larval gut microbiomes? Given the likely absence of gut microbes in pupae and newly eclosed adults [71], the presence of specialized, adult-enriched symbionts in larvae would suggest early acquisition, loss, and re-acquisition in adulthood. But due to the differing diets of larvae (e.g. solid vs. liquid food [57, 72]), and to their lack of a microbe-excluding proventricular filter, larval microbiomes may be distinct from those in adults, as may the mechanisms by which they are gained and constructed.

To test and refine hypotheses on the forces shaping *Cephalotes* gut microbiomes across castes and development, we apply quantitative PCR and amplicon 16S rRNA sequencing across 18 colonies from 13 *Cephalotes* species, in addition to a series of phylogenetic analyses assessing symbiont specialization and host-microbe cospeciation (Fig. 1). Our results give a colony-level view of gut microbiome evolution. And results from a recent metagenomic study [73] give insights into the functional differences of symbionts found across members of a *Cephalotes* colony. Given the known and candidate functions of these gut symbionts, our findings help to, further, uncover the mechanisms behind acquisition and retention of beneficial symbionts across divergent stages of a eusocial, holometabolous insect.

**Figure 1:**
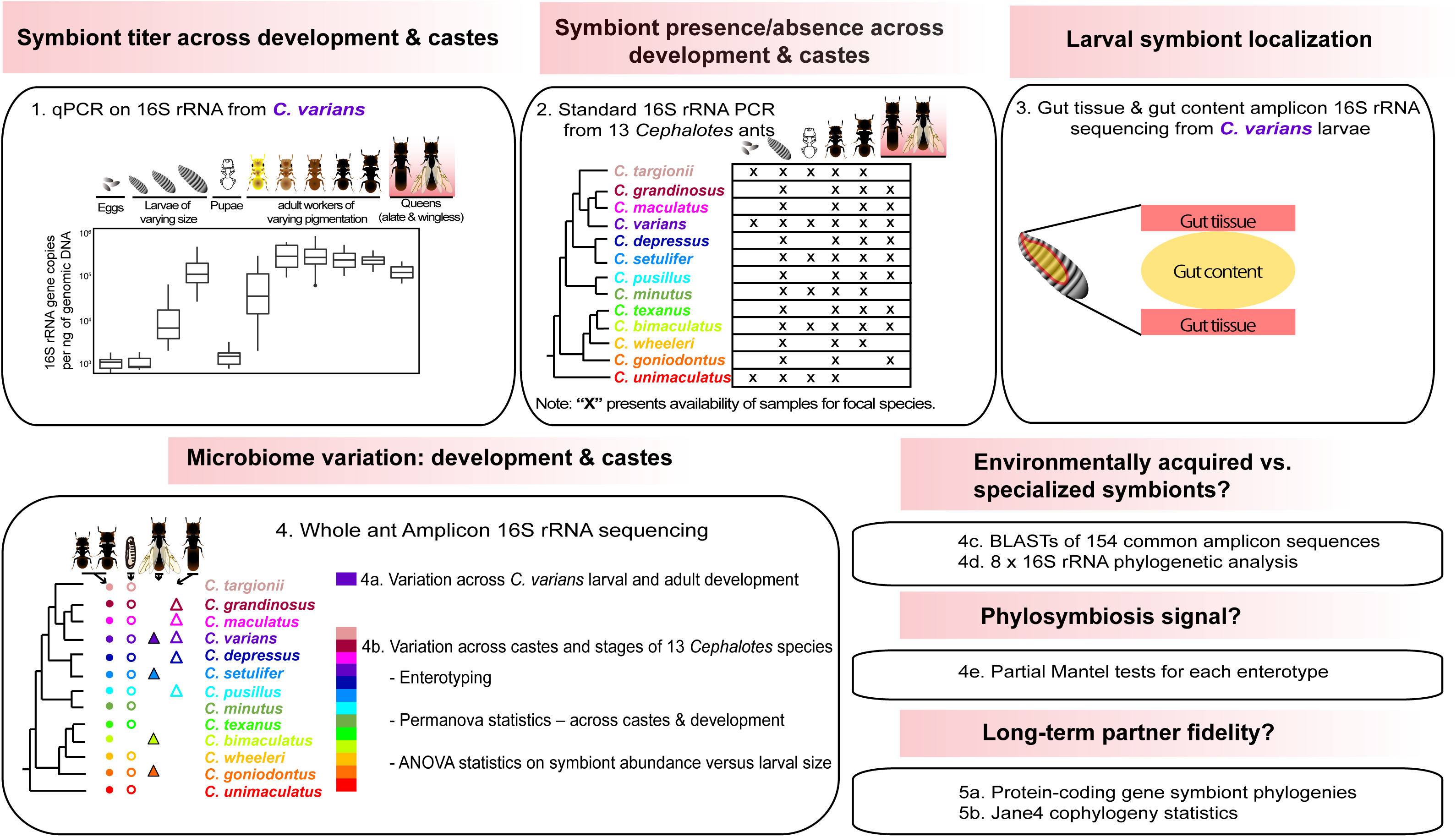
Methodological approach, and research goals of our study. Through extensive studies of *Cephalotes varians* ants, and 12 congeneric species, we aimed to pinpoint the timing of symbiont acquisition and proliferation, in addition to the composition of turtle ant gut microbiomes across castes and development. Further goals included localizing symbionts to the larval gut, inferring the likely origins of adult and larval symbionts (i.e. environmentally acquired vs. specialized, with plausible vertical transmission), and measuring the capacities for short-term partner fidelity to yield longer-term evolutionary patterns. New here, relative to prior work on this social, holometabolous insect, includes qPCR and 16S rRNA amplicon sequencing across-castes and developmental stages; phylosymbiosis analyses; and cophylogeny statistics.

## Methods

### Sample collections, DNA extraction, and PCR amplification

Ant samples used in this study were collected between 2012 and 2016 in Brazil, Costa Rica, Dominican Republic, Mexico, and United States (**Supplementary Table 1**). Collected samples were identified to the species, or near-species, level and preserved in 100% ethanol at – 80°C until further processing. The thirteen targeted species came from 18 colonies, and included *Cephalotes* aff. *bimaculatus*, *Cephalotes depressus*, *Cephalotes grandinosus*, *Cephalotes goniodontus*, *Cephalotes maculatus*, *Cephalotes minutus*, *Cephalotes targionii*, *Cephalotes pusillus*, *Cephalotes setulifer*, *Cephalotes texanus*, *Cephalotes unimaculatus*, *Cephalotes varians* and *Cephalotes* aff. *wheeleri*,

Prior studies have suggested that the vast majorities of microbes found in or on *Cephalotes* adults reside within the gut – primarily within the midgut and ileum [74, 75]. When coupled with the distinct morphology of adult vs. larval guts [76], we adopted an apples-to-apples approach involving DNA extraction of whole individuals. For each colony, we thus aimed to include whole-body, representative specimens from all available developmental stages, sexes, and castes, including workers, soldiers, queens, males, larvae, pupae, pre-pupae and eggs. Ant specimens were rinsed in 70% ethanol and then washed three times with molecular-grade water before DNA extraction. Whole bodies of single ants were then ground with a pestle in sterile 1.5-ml tubes after flash-freezing in liquid nitrogen. These samples were then used for individual-level DNA extraction, following the manufacturer’s protocol for Gram-positive bacteria, using DNeasy Blood and Tissue kits (Qiagen Ltd., Hilden, Germany). To detect possible laboratory- or reagent-derived contamination with bacterial DNA, we included “blank” samples in our DNA extraction batches, following the aforementioned protocol except for the addition of ant tissues. To assess the presence or potential absence of bacteria in our studied ants, we amplified the bacterial 16S rRNA gene using universal eubacterial primers 9Fa (5’ GAGTTTGATCITIGCTCAG 3’) and 1513R (5’ TACIGITACCTTGTTACGACTT 3’). We also checked for template DNA quality by amplifying a part of the ant mitochondrial cytochrome oxidase (*COI*) gene using universal insect primers LCO-1490 (5’ GGTCAACAAATCATAAAGATATTGG 3’) and BEN (5’ GCTACTACATAATAKGTATCATG3’). Further information on PCR primers, cocktail recipes, and cycling conditions can be found in **Supplementary Table 2**, and data on sample quality and bacterial presence are available in **Supplementary Table 3**.

In addition to whole-body extractions, we also dissected the guts from three older *C. varians* larvae, using sterilized fine-tipped foreceps. For each gut, we separated the gut wall/lining from the solid mass of gut content. DNA was extracted for these separate portions, as above, and the n=6 resulting samples were subjected to amplicon 16S rRNA sequencing as described below.

### Relative age estimates for all larvae and *C. varians* adults

Prior to DNA extraction, all specimens were photographed from the dorsal side by a digital camera connected to a Leica 80x dissecting microscope under constant-lighting conditions (**Supplementary Figure 1**). For all larval specimens, a calibration slide was used to measure length, which we used as an approximation of larval age [77, 78]. In *Cephalotes varians* we arbitrarily defined three age groups for larvae for use in statistics confined to this species. These included a young larval group with <=2mm body length, a middle-aged group with 2-3 mm body length, and an older group with >=3mm.

Beyond *C. varians*, we measured larval length from 13 colonies of 11 additional *Cephalotes* species. Using all of these data, we normalized larval length measures – dividing these values by the average worker mesosomal length from the same colony (**Supplementary Table 4)**. We used these normalized measures as an approximate means of relative larval age assignment.

To ascertain relative ages of adult *C. varians*, we measured pigmentation of the ant cuticle, adopting a method developed for leafcutter ants [79]. All photos of workers, soldiers and queens from this species were converted to black and white. Dorsal abdominal segments were then used for pigment quantification with ImageJ software. To specifically enable this, we recorded the mean gray value for each specimen, which we calculated by dividing the total gray value of imaged pixels by the total number of pixels in the selected region. For subsequent analyses, designed to compare microbiomes and bacterial titers across age, adult workers of *C. varians* were divided into four groups based on these gray-scale values, with arbitrarily chosen pixel value bins of 0-20 (oldest), 20-40, 40-60, and 60-80 (youngest – i.e. callow workers) (**Supplementary Table 4**).

### Bacterial quantitative PCR

To measure symbiont densities in different developmental stages and castes, we selected 151 DNA samples from individual ants across four *C. varians* colonies for quantitative PCR (qPCR) (**Supplementary Table 5**). qPCR quantification of bacterial 16S rRNA gene copies was performed with PerfeCTa SYBR Green FastMix (Quanta BioSciences Inc., Gaithersburg, MD, USA) using the primers 515F (5’-GTGCCAGCMGCCGCGGTAA-3’) and 806R (5’-GGACTACHVGGGTWTCTAAT-3’) [80]. One microliter of DNA template was added into a cocktail of 10 µl of PerfeCTa SYBR Green FastMix (Quanta BioSciences Inc., Gaithersburg, MD, USA), 0.125 μl of each primer at 20 μM (Invitrogen, Carlsbad, CA), and 8.75 µl of PCR water (Sigma). Reactions were performed on an Mx3005P qPCR System (Agilent Technologies, Santa Clara, CA, USA), using 1 cycle of 94°C for 10min and 35 cycles of 45s at 94°C, 60s at 50°C and 60s at 72°C; with a plate read at the end of each extension step. Melting curve analysis was performed immediately after the last amplification step, and included 90 cycles of 0.5°C increments from 50°C to 95°C, with 30s at each temperature. Averages were taken from two independent replicates performed for each ant sample across separate plates. For absolute quantification, each plate included a triplicated 1:10 serial dilution standard curve generated from linearized plasmid with the 16S rRNA gene of *Escherichia coli*. We consistently achieved r^2^ values of ≥0.999 and reaction efficiencies of ≥ 94%. Further details on generation and construction of the standard curve for qPCR protocols can be found in a previous study [81].

To normalize bacterial density estimates, we divided each 16S rRNA gene copy number average by the total ng of DNA for the given sample, obtained using the Quant-iT™ dsDNA High Sensitivity Assay kit (Life Technologies, Grand Island, NY, USA) on the GloMax Multi Detection system (Promega, Madison, WI, USA), following the manufacturer’s protocol. We used a one-way ANOVA to test whether normalized bacterial densities differed across developmental stages and castes, applyingTukey HSD post hoc tests to detect pairwise differences among development/caste groups.

### Bacterial 16S rRNA amplicon sequencing

DNA extractions from a total of 371 *Cephalotes* ants, 3 gut content and 3 gut wall samples from three *C. varians* colonies, and 33 blank extraction samples were submitted for amplicon sequencing of the V4 hypervariable region of the 16S rRNA gene following the protocols of the Earth Microbiome Project [80]. Library preparation and sequencing were performed at Argonne National Laboratory (Argonne, IL, USA), with paired-end 151 bp sequencing reactions on multiplexed samples across five Illumina MiSeq lanes.

Raw sequence reads were analyzed using mothur v.1.35.1 [82]. After read pair assembly, contigs from the five sequencing batches were grouped and further quality-filtered by removing contigs falling outside the 251-255 bp size range, those with ≥10 ambiguous nucleotides. and those with homopolymer tracts of ≥8 bp. Contigs from across the 404 sequence libraries were extracted using the get.groups command and concatenated into one FastA and group file.

The command unique.seqs was used to identify unique sequences and to reduce the size of the dataset, which was then subjected to a previously published decontamination procedure [83]. Briefly, the maximum relative abundance of each unique sequence was calculated across all blank libraries (value a) and, separately, all ant libraries (value b). Unique sequences with a ratio of value a:value b greater than or equal to 0.2 were classified as contaminants and removed from all libraries in the dataset. We then removed rare unique sequences that did not comprise at least 0.1% of one or more ant sequence libraries. This procedure resulted in the loss of >75% of the starting read number for 18 libraries. Due to this heavy contamination, we removed these libraries from subsequent analyses.

The remaining quality-filtered unique sequences (See details in **Supplementary Table 6**) were aligned to the SILVA database using the align.seqs command. The commands screen.seqs and filter.seqs were used to make sure that all aligned sequences mapped to the same region. We then performed chimera checking using UCHIME [84], removing all chimeric sequences. Finally, the remaining sequences were classified using the Ribosomal Database Project (RDP) 16S rRNA gene training set (version 16). Those classified as mitochondria, chloroplast, Archaea, or Eukarya were removed using the command remove.lineage.

Unique sequences were then assigned to operational taxonomic units (OTUs) at 97%, 98% and 99% sequence identity levels using the default settings of the opticlust algorithm in mothur. Using this output information, we compiled OTU tables containing the numbers of filtered reads assigned to each OTU per library, taxonomic assignments, and a representative sequence corresponding to the most abundant sequence assigned each OTU) (see details in **Supplementary Tables S7-S9**).

### Minimum Entropy Decomposition (oligotyping)

To more precisely characterize diversity, Minimum Entropy Decomposition (MED) was also performed on our filtered 16S rRNA amplicon Illumina sequencing libraries. The MED method identifies closely related but distinct bacterial sequences. In place of the aforementioned, sequence similarity-based OTU clustering, this method uses Shannon entropy to differentiate biologically meaningful signals from sequencing errors, providing an advantage over methods like 99% OTU clustering or unique sequence grouping. which use all variant positions, whether spurious or truly variable [85, 86]. To enable this, we formatted the unique sequence count table, fastA file, and taxonomy file obtained just before the aforementioned OTU picking steps (during mothur analyses), using the “mothur2oligo” tool (available from https://github.com/DenefLab/MicrobeMiseq/tree/master/mothur2oligo). Minimum entropy decomposition analysis was then performed using default parameters. A total 3744 of unique sequences were analyzed, and 1214 outliers were removed due to exceeding of the minimum substantive abundance and maximum variation allowed at each node. This tool classified the remaining 2530 unique sequences into 271 MED nodes, or “oligotypes”. Following the approach described above for 97%, 98%, and 99% OTUs, we then manually processed output files to create an oligotype OTU table containing, displaying the number of times each oligotype appeared in each sequence library, taxonomic information, and the most abundant (representative) sequence for each oligotype (**Supplementary Table 10**).

### Assessing beta diversity across colonies, developmental stages, and castes

To determine the similarity of microbial communities from different colonies, developmental stages, and castes of our best sampled species - *Cephalotes varians* – we computed Bray-Curtis distances between all *C. varians* libraries based on the 97% OTU table using the vegdist function within the vegan package for R v3.6.2 [87]. The Bray-Curtis distance matrix was used for Principal Coordinates Analysis (PCoA) through execution of the function pco within the package labdsv package for R [88]; PCoA plots were then visualized using R v3.6.2. We further used Adonis [89] from the vegan package to perform permutational ANOVAs comparing the composition of bacterial communities across colonies, developmental stages, and castes, based on Bray-Curtis or Jaccard distance matrices.

Due to the sensitivity of presence/absence analyses to contaminating sequences, we identified a minimum abundance threshold for filtering datasets prior to the aforementioned and below-described Jaccard index calculations. To achieve this, we leveraged the inclusion of army ant samples in one of our Illumina MiSeq lanes. Army ants have specialized symbionts that are not found in *Cephalotes*, including members of a group referred to informally as Unclassified Firmicutes [83]. Sequences from this symbiont group were detected in *Cephalotes* ant sequence libraries, comprising a maximum proportion of 0.002994 within the quality-filtered sequence libraries. As such, any sequence with a relative abundance of ≤0.002994 was considered as a “cross-contaminant” sequence in that library and was removed. Using this cross-contaminant filtered dataset, we were then able to compute Jaccard distances for 97%, 98%, and 99% OTUs, and for our oligotype clustering results.

To test whether *C. varians* workers harbor more variable microbial communities in early adulthood, closer to the expected time of trophallactic symbiont inoculation [41], we compared differences in variation in species composition across age for four age groups of *C. varians* workers. This was specifically enabled by implementation of a multivariate homogeneity of variance test using the Bray-Curtis distance matrix and the betadisper function within vegan package in R. As described above, age was approximated by cuticular pixel values.

Explorations of microbiome differences across colonies and developmental stages were also performed for the full dataset of 13 *Cephalotes* species. Visualization of symbiont composition was facilitated through the construction of heatmaps using heatmap.2 in R. In these graphics, generated separately for larvae and adults, we included the 39 most abundant OTUs, used for enterotype analyses (see below), plus one *Wolbachia* OTU. We also calculated Bray-Curtis and Jaccard distances among sequence libraries for these ant species, based on 97% OTUs, using the vegdist function (vegan package in R). We used Adonis to compare the composition of the microbial communities among different developmental stages, among colonies of the same ant species, and to search for interactions between developmental stage and colony.

### Enterotyping and indicator OTU analysis vs. caste and stage

To further identify ants with similar bacterial communities, samples from all *Cephalotes* species were clustered into ‘enterotypes’ using a previously described method [90]. Briefly, the 97% OTU table was converted to a biom-format OTU table using QIMME version 1.9.1 [91] for computation of the weighted UniFrac distance metric according to the relative abundance of the top 39 identified OTUs. All ant libraries were clustered by using the partitioning around medoids (PAM) clustering algorithm. The optimal number of clusters in our dataset was identified by the Calinski-Harabasz (CH) index (**Supplementary Table 11**). We then performed the between-class analysis using the package “ade4” in R, plotting the results using the s.class() function in package “adegraphics” in R. We identified indicator 97% OTUs, for which indicator values were >0.25, with p <0.05, using the indval function in the package “labdsv” in R (**Supplementary Table 12**)[88]. To gain further insight into the enterotype distributions across different developmental stages or castes, we plotted the frequency distribution of the three enterotypes across larvae with different normalized body lengths, as a proxy for larval age, and across workers, soldiers, alate queens, and wingless queens, with queens known more typically in the social insect literature as gynes.

### Assessing correlations between host phylogenetic relatedness, geography, and microbiome similarity

Including the data for all *Cephalotes* species, we used the labdsv package in R [88] to perform principal coordinates analyses (PCoAs) separately for each enterotype. Bray-Curtis distance measures, computed from oligotype distributions, were used as inputs for these analyses, and for each of our three separate analyses we included only those sequence libraries assigned to the focal enterotype. Symbols were used on PCoA plots to differentiate castes and developmental stages, while varying colors indicated phylogenetic origin.

Following up on these visual explorations, we conducted partial Mantel tests in the vegan package in R, to quantify the impacts of host phylogeny on microbiome composition. To account for possibly confounding effects of geography, between-sample geographic distances were computed based on our GPS coordinates for each sample (**Supplementary Table 1**) using the AMNH geographic distance calculator tool [92]. An average of geographic distances was calculated if there were more than one colony included for a specific ant species. Host phylogenetic distances were calculated from a time-calibrated *Cephalotes* phylogeny [93], using the cophenetic function within the ape package, implemented in R. Mantel tests were then implemented using the vegan package in R, testing for significant associations between bacterial community dissimilarity, host phylogenetic distance, and geographic distance among the collected ant colonies. As for the above-mentioned PCoA analyses, Mantel tests were run separately for each of the three enterotypes. For our measures of community similarity we utilized both Bray-Curtis and Jaccard distances, calculated from oligotype data, along with 99%, 98%, and 97% OTUs, and processed in separate analyses. We took the average of these dissimilarities for each species within a given enterotype analysis to reduce values to a single datapoint.

### Evolutionary origins of adult & larval-enriched symbionts – BLASTn searches

With the expectation that host phylogenetic vs. microbiome correlations could be driven by a history of specialized symbiose with faithful symbiont transmission across generations, we next set out to understand whether microbiomes from varying developmental stages or castes differed in their enrichment for host-specialized microbes. Prior studies on adult microbiomes have shown that approximately 95% of 16S rRNA sequences from *Cephalotes* workers are derived from specialized lineages, with long branch separation from free-living relatives and membership exclusive to bacteria from *Cephalotes* and its sister taxon *Procryptocerus* [7, 62]. To assess the extent to which bacteria of larvae hail from such specialized lineages, without the need for a systematic phylogenetic reassessment given prior study on the matter [7, 62, 65, 66], we performed BLASTn searches against the NCBI nr database using the most abundant, “representative”, 16S rRNA sequence from each of 154 97% OTUs. These OTUs comprised the entirety of those either exceeding the contamination-robust relative abundance threshold of 0.002994 in at least two sequence libraries across any caste or stage, or at least 0.05 in one library. When top hits: 1) came from another cephalotine ant, 2) exhibited >90% sequence identity vs. our query sequence, and 3) had been defined previously as a member of a cephalotine-specific clade, we considered the query OTU to be a specialized symbiont. Those not meeting these criteria were considered, at this stage, to be non-specialized.

### Evolutionary origins of adult & larval-enriched symbionts – phylogenetics

We, next, gave additional scrutiny to many of these putatively non-specialized OTUs, seeing if they might, in fact, fall into previously unforeseen clades showing cephalotine exclusivity. In our first approach, we prioritized the most abundant 97% OTUs from the larva-dominant orders Rhizobiales (Analysis #1), Enterobacterales (Analysis #2), and Lactobacillales (Analysis #3), identifying close relatives through BLASTn searches against the NCBI nr database. For query sequences we chose all unique 16S rRNA sequences from the focal OTUs with relative abundance ≥ 0.01 in at least one larval sequence library. We retained six to seven of the top BLASTn hits for each unique sequence for our phylogenetic analyses, In addition, we included 16S rRNA from related bacterial isolates cultured from *Cephalotes* larvae or adults [7]. We also included near-full-length 16S rRNA symbiont sequences obtained from *Cephalotes varians* by culture-independent Sanger sequencing of PCR products obtained with universal eubacterial primers (9Fa and 1513R). Larvae used in this direct sequencing were chosen upon inspection of amplicon sequencing data, and for each selected sample, the targeted OTU of interest made up ≥0.90 relative abundance, maximizing our chances of obtaining a ‘clean’ sequence.

The above-mentioned sequences were aligned using the Ribosomal Database Project Sequence aligner [94]. Alignments were uploaded to the CIPRES web portal [95], where maximum-likelihood phylogenies were inferred using RAxML-HPC BlackBox with default parameters [96]. Phylogenies were then visualized and annotated using the Interactive Tree of Life Website [97]. Using these figures, we searched for patterns suggestive of newly identified cephalotine-specific lineages – i.e. containing representatives from multiple *Cephalotes* and/or *Procryptocerus* species in monophyletic groupings, and showing low relatedness to bacteria from non-cephalotine habitats.

While the three aforementioned taxa comprised substantial majorities of our larval sequence libraries, we detected a range of additional sequences in association with turtle ant immatures, with some occasionally reaching high abundance. We dissected the evolutionary origins of such common, though less ubiquitous larvae-associated bacteria through five additional analyses. The first (Analysis #4), focused on unique sequences (n=73) from the 25 OTUs unambiguously classifying to the phylum Actinobacteria – a group of interest due to their common production of antimicrobials. For the next analysis (Analysis #5), we broadened our focus on the Lactobacillales to encompass less abundant sequences not part of our initial phylogenetic inference for this order in Analysis #3. We, next, studied the ambiguously classified, but occasionally abundant proteobacterial sequences of OTU046 (Analysis #6). Additionally analyzed were sequences from OTU116 (Analysis #7), an ambiguously classifying group with a top BLASTn hit showing ∼85% sequence identity to a specialized Sphingobacteriales symbiont from *Cephalotes*. For our final analysis (#8), we focused on a subset of the aforementioned 154 OTUs, ranking within the top 100 of all named OTUs that: 1) did not belong to an OTU assigning as a specialized cephalotine bacterium via BLAST; 2) had not been studied extensively in previously published phylogenetic analyses on ant-associated bacteria (e.g. *Wolbachia*, Entomoplasmatales – Russell et al. [98]; Funaro et al. [99]); and 3) were not included in phylogenetic Analyses #s 1-7. Top BLASTn hits were retrieved from NCBI for these analyses, with subsequent ClustalO alignment, and PhyML-implemented phylogenetic inference in SeaView. Additional details on methodology can be found in the supplementary figure legends for each inferred phylogeny.

### Proportions of the microbiome comprised by specialized symbionts

Our above BLASTn and phylogenetic assessments allowed us to infer whether bacteria in *Cephalotes* were likely obtained from environmental sources (i.e. those with close relatedness to bacteria from non-cephalotine habitats) or through social inheritance promoting partner fidelity (i.e. those from cephalotine-specialized clades). To ascertain whether the adult castes or larvae varied in their proportions of microbiomes comprised of cephalotine-specialist bacteria, and hence the means by which they likely acquire their microbiomes, we computed average relative abundance values for each of the 154 aforementioned, threshold-exceeding OTUs in each stage and caste, per colony. We then computed sums for those from specialized vs. non-specialized clades, computing pooled averages and medians across *Cephalotes* species for comparison.

### Distributions and abundances of bacterial taxa across stages and castes – impacts of larval size

With a greater understanding of bacterial relatedness, we next built off of the earlier-described enterotyping/indicator OTU analyses, wishing to ascertain the identities of bacteria with potentially varying abundance across stages of larval development. We focused on 47 OTUs equaling or exceeding relative abundance of >0.0095 (i.e. nearly 1% or more) when averaged across the larvae of at least one colony, examining the relative abundance of each of these selected 97% OTUs across all individual larvae of varying size, and applying the >0.002994 threshold-filter for declaring symbionts as present/absent. To simplify graphics illustrating symbiont abundance across larval stages, and to avoid a large number of statistical tests and the necessary statistical corrections, we pooled relative abundances from a number of related 97% OTUs as described in a subsequent figure legend.

Using these data we computed stem and whisker plots graphing quartiles, medians, and outliers of symbiont relative abundance from these larvae-enriched OTUs across 11 larval size bins. All but the largest of these size bins, with only n=3 larvae, contained n=10 larvae of similar size. This binning approach enabled the use of an ANOVA framework, which was preferred given the uncertainty as to whether symbiont abundance should change monotonically with larval age. With these ranked size bins as treatment levels for our categorical x-variable (i.e. larval size, as a proxy for age), we performed ANOVAs on arcsin transformed relative abundances of larvae-enriched OTUs, assessing whether they change in predominance over development.

### Testing for cospeciation between *Cephalotes* species and their dominant *Cephaloticoccus* (Opitutales) symbionts

Recent metagenome sequencing reinforced the prevalence and high relative abundance of *Cephaloticoccus* (Opitutales) symbionts [100] in *Cephalotes* turtle ants [7]. To understand whether these seemingly ubiquitous and dominant symbionts might cospeciate with their ant hosts we mined these metagenomes for DNA sequences enabling finer-scale phylogenetic inference than the presently sampled V4 region of 16S rRNA. Toward this end, we chose five genes – *uvrB*, *23S rRNA, dnaA*, *recA*, and *rpoB* – conserved in many bacterial clades, including Verrucomicrobia [101–103]. Using these sequences, we performed a series of BLASTn analyses against the Integrated Microbial Genes and Genomes (IMG) database. We began by extracting these genes from genomes of two cultured isolates, *Cephaloticoccus primus* (host: *Cephalotes rohweri;* IMG Project ID: Gp0154034) and *Cephaloticoccus capnophilus* (host: *Cephalotes varians;* IMG Project ID: Gp0110136). We, next, used these sequences as queries in BLASTn searches performed on IMG, downloading all hits exceeding 30% query coverage and a range of chosen sequence identity thresholds, which corresponded to natural breakpoints in the data – below which the hits appeared much less similar and appeared to derive from scaffolds classifying to other taxa (i.e. not Verrucomicrobial or Opitutales). Thresholds chosen were: 83% for *uvrB* and *recA*; 87% for *dnaA*; 88% for *rpoB*; and 98% for 23S rRNA. The top two BLAST hits from NCBI, excluding those from *Cephalotes* hosts, were downloaded as outgroups.

We aligned sequences in SeaView using the muscle algorithm, and then performed maximum likelihood phylogenetic analyses using RAxML on the Cipres Web portal [104]. After preliminary analyses we identified duplicate sequences – i.e. those from the same host species that grouped into a host-species-specific clade at the tips of the phylogeny. These include partial gene sequences (i.e. from the 5’ and 3’ ends) and replicate samples from the same host species obtained from different colonies. To prevent issues from pseudoreplication, we eliminated duplicates – keeping only the longest sequence from each such cluster. We then re-ran the RAxML analysis, performing 1000 bootstrap replicates. Trees were saved and visualized using the interactive Tree of Life (iTOL) website. We kept the two trees with the highest average bootstrap support at each node – i.e. the *rpoB* and *uvrB* genes. Applying default parameters and event costs, we then compared the topologies of these two gene trees to that of a *Cephalotes* host tree modified from Price et al. [93], using the cophylogeny software Jane4. A sample size of 1000 randomized tip mapping permutations was used to evaluate statistical significance.

## Results

### Symbiont densities vary across *Cephalotes varians* development – qPCR assays

In our first assessment of microbiome variation across *Cephalotes* ants, we utilized 16S rRNA-based qPCR assays to measure bacterial titer in four colonies of *C. varians*. Our two-way ANOVA detected no significant effect of colony on normalized bacterial density (F-value = 0.742; p=0.529; **Supplementary Table 13**). In contrast, this same ANOVA model revealed a significant effect of developmental stage/caste designations (F-value = 95.08; p < 2 × 10^-16^), suggesting variation in symbiont density between some combination of eggs; larvae at different stages of development; prepupae; pupae; soldiers; queens; or adult workers of varying age, as approximated by pigmentation measures.

Tukey’s post-hoc HSD tests allowed us to ascertain which of these stages or castes harbored varying bacterial loads. Eggs and young larvae (<2) had the lowest numbers of bacteria across the four studied *C. varians* colonies (Fig. 2a), with average 16S rRNA gene copy numbers (per ng total DNA) respectively estimated at 4.34 × 10^3^ and 5.08 × 10^3^. Unnormalized qPCR measures were comparable between eggs and our blank, negative control samples (8.63 × 10^2^ vs. 5.00 × 10^2^, respectively) suggesting that *C. varians* ants are nearly sterile upon hatching. The first detectable change in bacterial titer unfolded as larvae transitioned from small (<2 mm) to intermediate size (2-3 mm), with 16S rRNA copy number/ng DNA increasing to 2.82 × 10^4^ (p-value for Tukey’s HSD_<2 vs. 2-3_ = 0.040; **Supplementary Table 13**). Titers increased again as larvae aged further, reaching 2.50 × 10^5^ for individuals >3mm in length (p-value for Tukey’s HSD_2-3 vs. >3_ = 0), before falling drastically at the pre-pupal stage to 3.17 × 10^3^ copies/ng DNA (p-value for Tukey’s HSD_>3 vs. pre-pupae_ = 0). There was no detectable change in titer into the pupal stage, with normalized 16S rRNA copy number estimates hovering at 2.49 × 10^3^ (p-value for Tukey’s HSD_pre-pupae vs. pupae_ = 0.999). But callow workers (pixel value ≥ 60) showed a spike in symbiont density, with normalized copy number estimates rising to 1.31 × 10^5^ (p-value for Tukey’s HSD_pupae vs. adult-pixel>60_ = 0). In addition, workers at this young adult stage showed some of the highest variability detected for symbiont titer (Fig. 2a). Adult workers at the next age class (pixel value = 40-60) showed an additional, but smaller increase in bacterial titer, with 5.45 × 10^5^ 16S rRNA copies/ng DNA (p-value for Tukey’s HSD_adult-pixel≥60 vs. adult-pixel-40-60_ = 2 × 10^-7^). Further aging was not detectably associated with titer changes, and workers at these older ages appeared to show greater consistency in 16 rRNA copy number when compared to callows (Fig. 2a**; Supplementary Table 13**).

**Figure 2:**
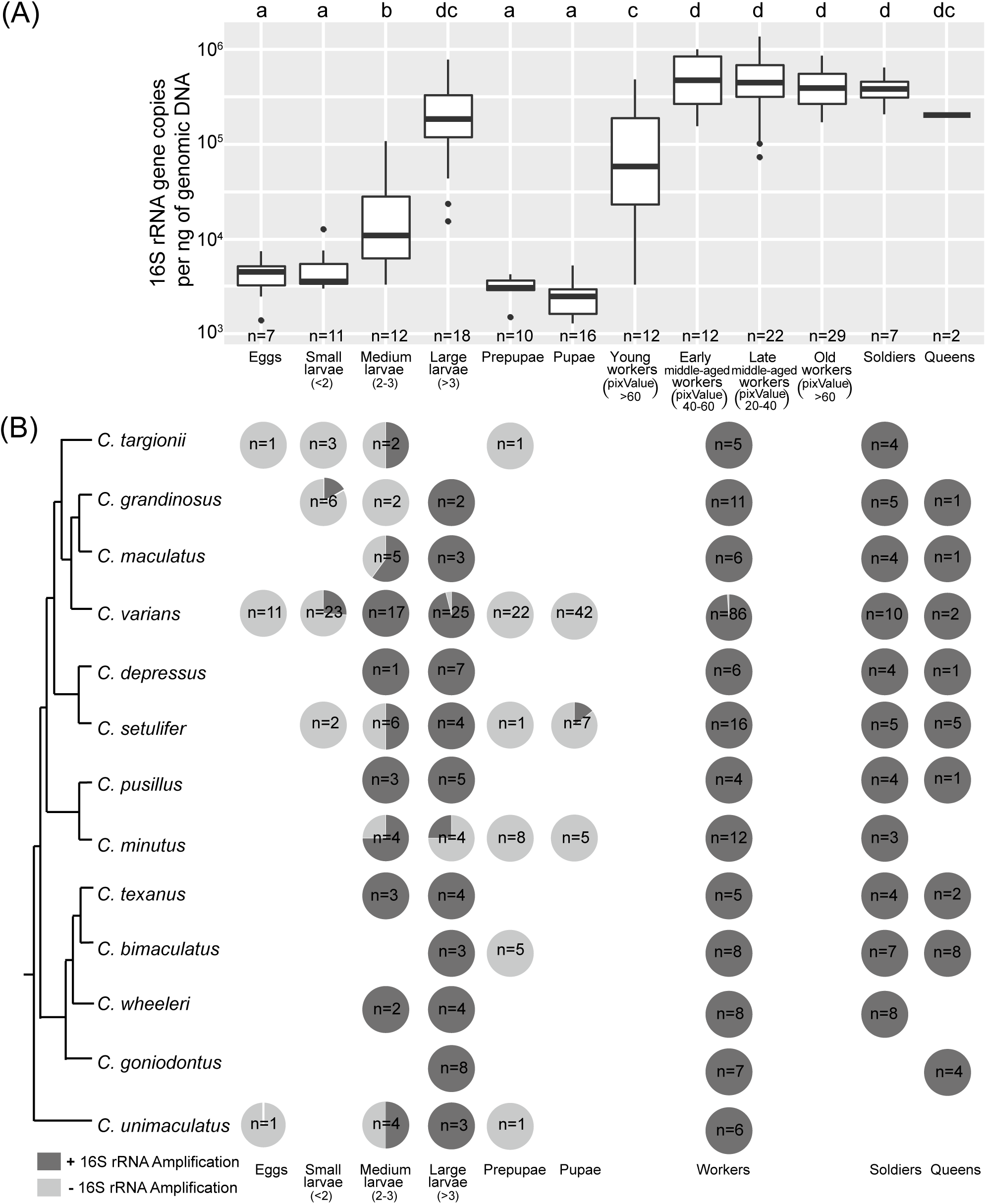
Bacterial presence and abundance across developmental stages and castes of *Cephalotes* ants. A) Results of qPCR assays. Stem and whisker plots illustrate medians, quartiles, and outliers of 16S rRNA copy number/ng genomic DNA for each stage/caste. Letters above the graph reveal results of Tukey’s HSD comparisons, with different letters revealing castes/stages with differing quantities of bacteria. Sample sizes indicated at bottom of graph. B) Relatedness among the thirteen studied *Cephalotes* species (Price et al. [93]), and our overall results of standard PCR with universal bacterial 16S rRNA primers indicating whether individuals from particular castes/stages had detectable titers of bacteria (dark gray) or no detectable quantities (light gray). Results shown in the form of pie charts to indicate the proportions of each caste/stage amplifying, with sample sizes inside each pie chart. Alate and wingless queen data are pooled here. PCoA analysis on Bray-Curtis distances among ant sequence libraries, based on compositions defined at the 97% OTU level.

The most mature workers (pixel value = 0-20) showed pigmentation overlap with one alate queen, one wingless queen, and ten soldiers (**Supplementary Table 4**), suggesting similar age. Comparisons of bacterial titers among these adult categories suggested little difference in symbiont quantities, with 16S rRNA copy numbers/ng DNA equaling 3.57 × 10^5^ (alate) and 1.14 × 10^5^ (wingless) for the single queens, and averages of 4.03 × 10^5^ and 4.18 × 10^5^ for soldiers and workers. Accordingly, p-values from pairwise Tukey’s HSD comparisons were non-significant, ranging from 0.997 – 1.0 (**Supplementary Table 13**).

### Symbiont presence/absence varies across *Cephalotes* development – standard PCR assays

After discovering the above-described developmental time course for symbiont proliferation, and ascertaining that all female castes harbor symbionts during adulthood, we used standard PCR amplification with universal 16S rRNA gene primers as a crude approximation of symbiont presence/absence across 522 ants spanning various castes and stages of *C. varians* and twelve additional *Cephalotes* species (Fig. 2b**; Supplementary Table 3**). Eggs from three species (n=13 samples) failed to amplify with universal bacterial PCR primers. But amplification did occur for 20.6% (n=34) of young larvae (i.e. normalized size <2) from across four *Cephalotes* species. In contrast, 74.6% (n=47) of middle-aged larvae from 11 species (i.e. standardized size class: 2-3) yielded 16S rRNA PCR amplification, as did 94.5% (n=73) of older larvae (i.e. standardized size class: >3) from 12 cephalotines. All 37 pre-pupae, sampled across seven *Cephalotes*, failed to amplify with universal bacterial primers, while amplification was seen for only 1/57 pupae obtained from three host species. At the adult stage, all wingless queens (n=18, from 6 species), alate queens/gynes (n=7, sampled from 4 species), and soldiers (n=57, from 11 species) yielded PCR amplification in these 16S rRNA assays, as did 177/178 workers (from 13 species).

### Whole ant community composition across castes and stages of *Cephalotes varians*

With a sequence of bacterial acquisition, proliferation, loss, and reacquisition being suggested by the aforementioned data, we next set out to understand the composition of symbiotic bacterial communities across castes and development. We began with a study of *C. varians*, subjecting 145 samples with PCR-detectable bacterial communities to amplicon sequencing of the bacterial 16S rRNA gene using Illumina MiSeq technology. We removed one library where the number of reads was less than 1,000, and focused on the remaining 144. After removing contaminants, chimeras, and non-bacterial sequences, 1,597,177 quality-controlled sequences remained for further analysis, an average of 11,092 sequences per library (see unique sequence, OTU, and oligotype tables in **Supplementary Tables S6-S14**).

Permutational multivariate ANOVA statistics indicated that some combination of species composition and/or relative abundances of bacterial community members differed significantly across developmental stages of *C. varians* (Fig. 3a; **Supplementary Table 14**; Bray-Curtis: *F* = 129.617, *R^2^* = 0.415, *P* = 0.001; Jaccard: *F* = 89.087, *R^2^* = 0.342, *P* = 0.001), and also among colonies (Bray-Curtis: *F* = 7.813, *R^2^* = 0.075, *P* = 0.001; Jaccard: *F* = 6.329, *R^2^* = 0.073, *P* = 0.001). Across all larval libraries from four *C. varians* colonies, six dominant 97% OTUs, including Rhizobiales (OTU001 and OTU004), Lactobacillales (OTU007, OTU022 and OTU025), and Enterobacteriales (OTU006), accounted for an average of 87.1% of reads. In very young larvae (i.e. normalized body length < 2), Rhizobiales OTU004 was the only dominant bacterium. Another type of Rhizobiales, from OTU001, became common subsequently, beginning in middle-aged larvae (i.e. normalized body length 2-3) and continuing into the oldest larval stages (i.e. normalized body length >3). Lactobacillales OTUs 007, 022, and 025, as well as Enterobacteriales OTU006, were the most abundant taxa in larvae assigned to the oldest age class, but were common also in some middle-aged *C. varians* larvae.

**Figure 3:**
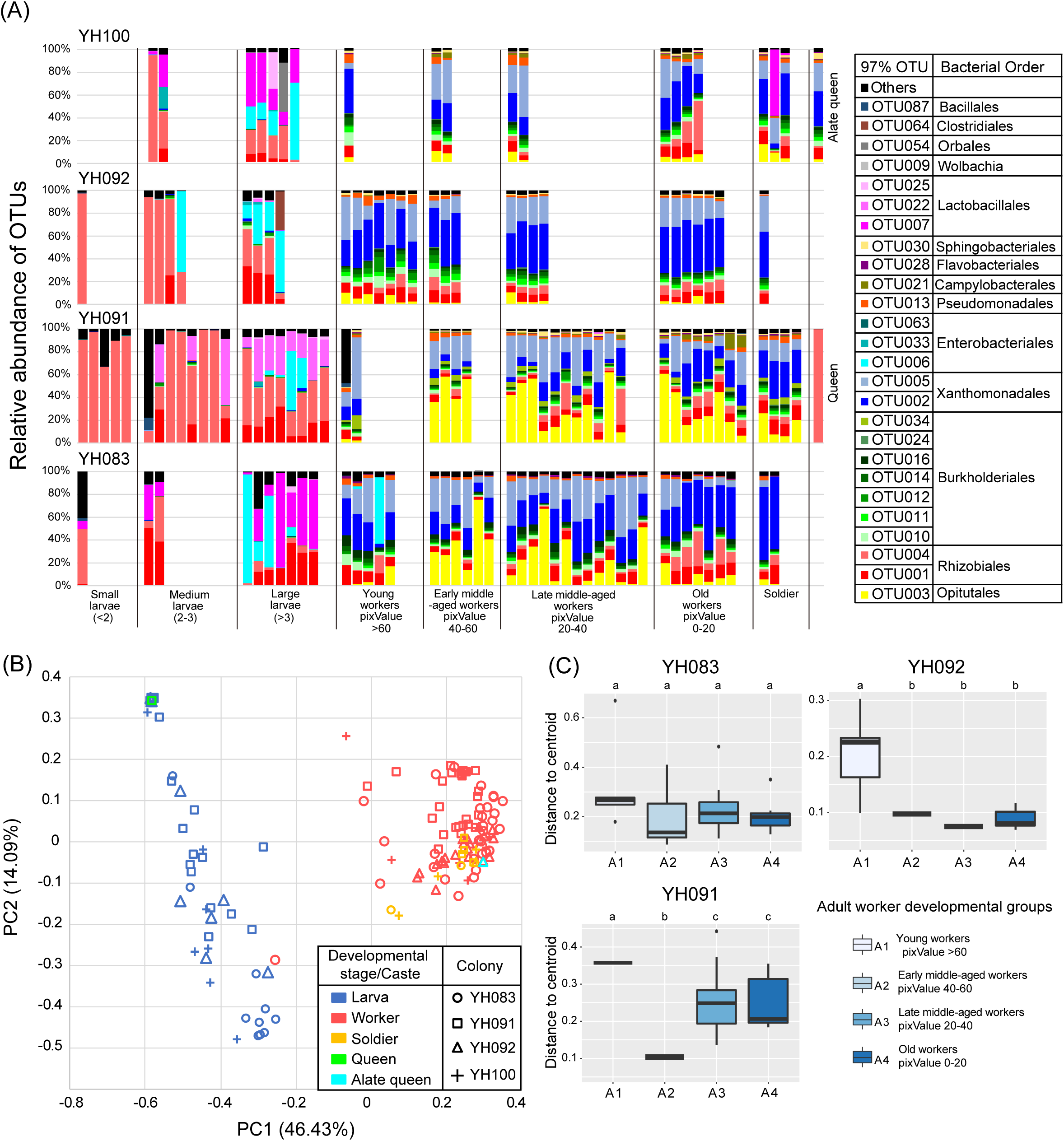
Bacterial composition, beta-diversity, and community dispersion among developmental stages and castes of *Cephalotes varians*. All data obtained via Illumina amplicon sequencing of the 16S rRNA gene from individual ants. A) Taxonomic composition of each individual sequence library from *C. varians*, across four colonies (YH100, YH091, YH092, YH083). Stacked bar graphs illustrate bacterial composition at the 97% OTU level, with similarly colored OTUs hailing from the same orders. B) PCoA plot based on Bray-Curtis dissimilarity values, measured among pairs of *C. varians* sequence libraries defined at the 97% OTU level. Castes and developmental stages are distinguished by unique symbols and colors. C) Betadisper analyses, performed within *C. varians* colonies, to ascertain whether younger workers harbor more variable microbiomes. The four age classes analyzed here include the most lightly pigmented, youngest workers from the G1 category (cuticular pixel value > 60), and workers of successively older ages with darkening pigmentation, including G2 (pixel value 40-60), G3 (pixel value 20-40) and G4 (pixel value 0-20). Stem and whisker plots show medians, quartiles, and outliers for the community variability measure of distance to the centroid. Letters at the tops of each graph indicate significant differences in pairwise comparisons from the betadisper permutation test for homogeneity of multivariate dispersions, based on the Bray-Curtis distance matrix computed from 97% OTUs. Age classes with different letters exhibited significant differences in microbiome variability within the given colony. Small sample sizes in some age classes, for certain colonies, limited statistical power, as did inclusion of only 3 out of 4 colonies due to more extreme sample size issues in colony YH100.

With the exception of one mature queen, discussed below, microbiomes of *C. varians* adults were quite different from those of larvae (e.g. Fig. 3a). To begin, Lactobacillales and Enterobacteriales were extremely rare in workers, soldiers, and one alate queen/gyne, while Rhizobiales were ubiquitous, but only modestly abundant. Instead, adults harbored OTUs found rarely in larvae, including highly abundant OTUs from the orders Opitutales (OTU003) and Xanthomonadale (OTU002 and OTU005), seven frequently present OTUs from the Burkholderiales, and individual OTUs from the orders Pseudomonadales, Flavobacteriales, Campylobacterales, and Sphingobacteriales. The dominant OTUs from these orders accounted for an average of 93.95% of the sequence reads in each library.

We re-ran our permutational multivariate ANOVA analysis using a 97% OTU table containing samples from the most darkly pigmented and, hence, most mature workers and soldiers (pixel value <20). Our assessment of caste, however, revealed no differences among these two groups, whether beta diversity was measured through the Bray-Curtis or Jaccard index (**Supplementary Table 14**; Bray-Curtis: *F* = 2.376, *R^2^* = 0.043, *P* = 0.074; Jaccard: *F* = 2.023, *R^2^* = 0.029, *P* = 0.107). Our remaining caste sampling lacked sufficient replication for statistics. But anecdotally, a virgin winged queen/gyne (colony YH100) harbored a similar gut bacterial community in comparison to its sibling workers and soldiers, while the previously mentioned mature, wingless queen (colony YH091) was dominated by Rhizobiales OTU004, similar to some young larval samples from the same colony (Fig. 3a).

Principal Coordinates Analysis (PCoA) on the full 97% OTU table reinforced the above results (Fig. 3b). For example, gut communities of larvae were separated from those of workers and soldiers, along the first PCoA axis (46.43%). In fact, only two adults harbored microbiomes clustering with microbiomes of larvae, including a worker dominated by Lactobacillales OTU007, and the previously described wingless queen. Matching the compositional patterns from Fig. 3a, the symbiotic community of the alate queen/gyne clustered along the first axis with all worker samples and all but one of the above-described solider samples.

Bar graphs suggested high microbiome variability and occasionally unusual composition among the youngest, most lightly pigmented workers (Fig. 3a). To pursue this pattern, we ran multivariate homogeneity of variance tests separately in the three *C. varians* colonies with adequate sampling across stages (Fig. 3c). We found that the average distance to the centroid for the youngest worker group (pixel value = 40-60), and occasionally members of the next youngest cohort (pixel value = 40-60), was significantly higher than for the older worker age groups, in two of the three colonies with sufficient sample sizes (**Supplementary Table 15**). While still somewhat preliminary, these patterns suggest higher variability in microbiome composition during early adulthood.

### Gut localization of larvae-associated symbionts in *Cephalotes varians*

Microbes dominating amplicon sequencing libraries from gut tissue and solid gut contents of three older *C. varians* larvae were highly similar to the communities seen in the above-described amplicon sequencing of whole-larva DNA extractions (**Supplementary Figure 2** vs, Fig. 3a). Highly abundant were Rhizobiales OTUs 001 & 004, Lactobacillales OTU007, and – in one case – Enterobacteriales OTU006. Present at lower titers were OTUs from adult-enriched symbionts in the Xanthomonadales and Burkholderiales. Similarly present at lower relative abundance was OTU025 from the Lactobacillales. Sequences from additional OTUs made up <10% from each library. OTU004 (Rhizobiales) was consistently more abundant in all three gut tissue vs. gut content comparisons. The remaining, aforementioned OTUs showed similar relative abundances across the two sample types. Given the small number of dissected samples, these trends require further study.

### Whole ant microbiome composition across castes and stages of 13 *Cephalotes* species – 97% OTU distributions

We further examined whether patterns of community structure found in *C. varians* were consistent across the *Cephalotes* genus, expanding our study to include 12 additional species. A total of 209 further 16S rRNA sequence libraries were generated toward this end, with an average of 8714 sequences per library after quality and contaminant filtering. Our remaining analyses included 192 of these QC’d libraries (i.e. all those except for libraries from 16 callow adults and 1 pupa). Focusing on 97% OTU clusters, a permutational multivariate Analysis of Variance revealed significant differences in the bacterial communities between larvae and adults (**Supplementary Table 16**; Adonis permANOVA tests: p < 0.05) for all *Cephalotes* species, except *C. targionii.* This was true for both Bray-Curtis and Jaccard measures of beta-diversity.

To characterize the taxa driving these differences across *Cephalotes* development, we constructed a heatmap showing 40 abundant 97% OTUs in all 336 whole ant sequence libraries (Fig. 4), in addition to a heatmap illustrating average 97% OTU relative abundance for each stage and caste per colony (**Supplementary Figure 3**). Similar to patterns for *C. varians*, larval samples of different *Cephalotes* species harbored overlapping, yet distinct symbiotic communities compared to adults, with most adult-enriched symbionts exhibiting rarity, or low-abundance, in larvae (**Supplementary Table 17**). Using a relative abundance threshold of 0.00294 to ascertain presence/absence (explained below), and focusing on OTUs found in least 30% of the 154 sampled adult workers (**Supplementary Figure 4)**, we found that the most depleted OTUs came from the Opitutales (OTU003 – 98.1% of workers vs. 8.7% of larvae), Comamonadaceae (OTU020 – 74.0% of workers vs. 8.7% of larvae), Xanthomonadales (OTU005 – 70.8% of workers vs. 1.9% of larvae), Pseudomonadales (OTU013 – 69.5% of workers vs. 4.9% of larvae), Flavobacteriales (OTU028 – 46.8% of workers vs. 1.0% of larvae), Campylobacterales (OTU021 – 40.3% of workers vs. 3.9% of larvae), and Sphingobacteriales (OTU030 – found in 33.1% of workers vs. 0% of 103 larvae). Showing more intermediate patterns were Alcaligenaceae OTUs 010 (93.5% of workers vs. 35.0% of larvae), 012 (97.4% of workers vs. 40.8% of larvae), and 016 (83.8% of workers vs. 19.4% of larvae), along with Xanthomonadales OTU002 (74.0% of workers vs. 17.5% of larvae).

**Figure 4:**
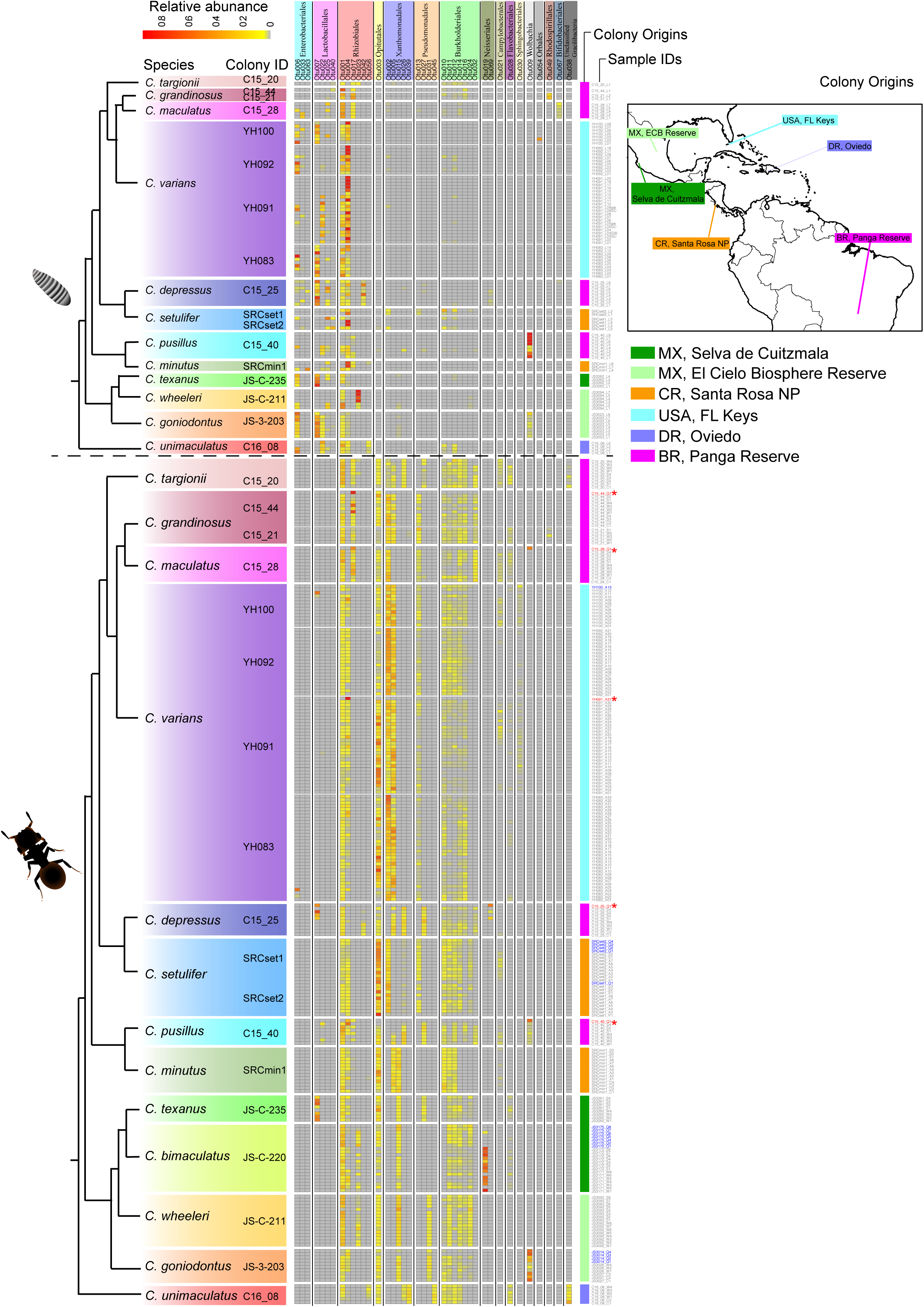
Heatmap illustrating 97% OTU composition across individual *Cephalotes* ants. Shown are geographic origins and patterns of host relatedness for all 336 sequence libraries, plus those from callow ants (n=16 samples with “_C#” in Sample ID column) and a pupa (n=1 sample with “_P#” under “Sample ID column). Libraries from alate queens (blue font) and wingless queens (red font w/asterisks) are highlighted in the Sample ID column. Upper panel illustrates data from larvae, while lower panel shows results from all other stages. Columns show each of the 40 focal OTUs (39 used for enterotyping analysis, plus *Wolbachia*). Rows show individual sequence libraries. Relative abundance data for each OTU in each sequence library are shown with a gray-yellow-red heatmap as illustrated in the upper left.

Using the same abundance threshold for presence/absence assignment, and focusing on OTUs found in at least 10% of larval samples (**Supplementary Figure 4; Supplementary Table 17**), OTUs enriched in larvae included Lactobacillales OTUs 007 (3.2% of workers vs. 36.9% of larvae), 022 (2.6% of workers vs. 28.2% of larvae), 025 (0% of workers vs. 35.9% of larvae), 082 (0% of workers vs. 11.7% of larvae), and 086 (0% of workers vs. 10.7% of larvae). Enterobacteriales OTU006 (1.3% of workers vs. 45.8% of larvae) and OTU033 (0% of adults vs. 13.6% of larvae) showed a similar trend, as did Actinobacteria OTUs 062 (Actinomycetales; 0.6% of workers vs. 10.7% of larvae) and 080 (Bifidobacteriales; 0% of workers vs. 16.5% of larvae).

Distinct from these patterns were trends for Rhizobiales symbionts, which were common in both adults and larvae. OTU001 was found, for instance, in 100% of workers and 79.6% of larvae. OTU004 was in 76.6% of workers and 83.5% of larvae. Though less prevalent, OTUs 017 (15.6% of workers vs. 19.4% of larvae) and 053 (11.0% of workers vs. 17.5% of larvae) showed similar patterns.

Maximum and average relative abundance of OTUs across development revealed qualitatively similar trends, indicating that adult-enriched symbionts with low- to intermediate-prevalence across larvae are usually found at low relative abundance within these immatures (**Supplementary Figure 4; Supplementary Table 17**). Two Alcaligenaceae OTUs, 010 and 012, provided interesting exceptions – with maximum relative abundances in larvae (0.309 and 0.157) either exceeding or approaching those in workers (0.121 and 0.188). Rhizobiales, with their dual-stage prevalence, showed similar patterns, with greater average and maximum relative abundance in larvae vs. adults, as exemplified by OTUs 001, 004, and 053, but not OTU017. Larvae-enriched symbionts were, conversely, most often rare in adults when detected, with the exceptions of OTU007 (Lactobacillales) and OTU006 (Enterobacteriales) – both reaching sporadically high maximum relative abundance in a small number of workers (0.506 and 0.578).

Many of the above general trends of larval vs. adult differences – gleaned from adult workers – were recapitulated when comparing alate queens/gynes and soldiers against larval immatures (Figs. 3**, S3; Supplementary Table 17**). But while these adult castes harbored fairly similar microbiomes there were some key exceptions. In *C. depressus* and *C. pusillus*, for instance, Lactobacillales were seemingly enriched in soldiers relative to workers. Furthermore, in the former species a rare bacterium from the Neisseriales was also common in soldiers but rare in workers. In contrast, Campylobacterales appeared more abundant in workers than in soldiers in 5 out of 7 colonies harboring this microbe, suggesting interesting avenues for future study.

When expanding our adult caste comparisons to include five mature queens, sampled from five *Cephalotes* species, we saw considerably greater microbiome divergence (**Supplementary Figure 3)**. With microbiomes bearing little resemblance to those of other adult counterparts, symbiotic communities of wingless queens instead, resembled those of some larvae, with dominance by a subset of Rhizobiales (i.e. OTUs 004 and 017). Overall, mature queen microbiomes had lower diversity, with an apparent absence of many of the adult-enriched OTUs present in their mature offspring. Representing a slight exception to the Rhizobiales-enrichment trend was a queen from *C. depressus*, possessing such bacteria at lower titers, but showing enrichment, instead, for the same Neisseriales and Lactobacillales found in abundance within their soldiers (see above). The mature queens from *C. pusillus* and *C. maculatus* colonies uniquely harbored *Wolbachia*, in addition to Rhizobiales. This was a conspicuous pattern for the latter given this symbiont’s apparent absence from the queen’s presumed larva, soldier, and worker offspring.

### Whole ant microbiome composition across castes and stages of 13 *Cephalotes* species – oligotype distributions

To understand strain-level diversity in *Cephalotes*-associated bacterial communities across developmental stages and castes, we looked at 16S rRNA oligotype distribution within the 40 focal symbiont OTUs from Fig. 4 (**Supplementary Figure 5**). Although slowly-evolving genes, like 16S rRNA, obscure higher strain-level resolution, we detected several distinct oligotypes within most dominant OTUs. Similar to our 97% OTU result, permutational multivariate Analysis of Variance revealed that larvae and adults from 11 out of 12 species harbor significantly different bacterial communities at this narrower clustering width (**Supplementary Table 16**; Adonis permANOVA tests: p < 0.05).

Despite these differences, we found identical oligotypes across siblings from these stages for a handful of trans-developmentally shared OTUs, including those from the Rhizobiales (OTUs 001, 004, 017, 023, 042 and 056). Sibling workers, soldiers and winged queens from the same colony similarly shared oligotypes, suggesting a homogenizing effect of colony on microbiome composition. Microbiomes of wingless queens remained non-diverse at the oligotype-level. And those sharing 97% OTUs with colony-mates shared oligotypes with them, as well. The sole exception came for a single oligotype belonging to OTU004, found in the mature queen, and n=1 larva of *C. varians* colony YH091, but in no other members of that social unit (**Supplementary Figure 5)**.

### Whole ant microbiome composition across castes and stages of 13 *Cephalotes* species – enterotyping analysis inferred from 97% OTU composition

With our results suggesting impacts of development and caste on microbiome composition, we aimed to further characterize symbiotic bacteria of *Cephalotes* ants through enterotyping analysis on our 336 whole ant Illumina sequencing libraries. Results suggested the presence of three community types, or “enterotypes” (Fig. 5a), with Calinski-Harabasz values reaching a peak of 0.829 at n=3 clusters, compared to CH values of 0.662 and 0.781 under models with n=2 and n=4 (**Supplementary Table 11)**. Enterotype 1 was mainly comprised of microbiomes from workers, soldiers, and 14 out of 16 alate queens/gynes (Fig. 5c). Indicator OTU analyses revealed that this cluster was dominated by 97% OTUs from the Opitutales (n=1 OTU), Burkholderiales (n=6 OTUs), Xanthomonadales (n=2 OTUs), Flavobacteriales (n=1 OTU), Campylobacterales (n=1 OTU), and Sphingobacteriales (n=1 OTU), with the latter taxon being concentrated in *C. varians* (Fig. 5b**; Supplementary Table 12**).

**Figure 5:**
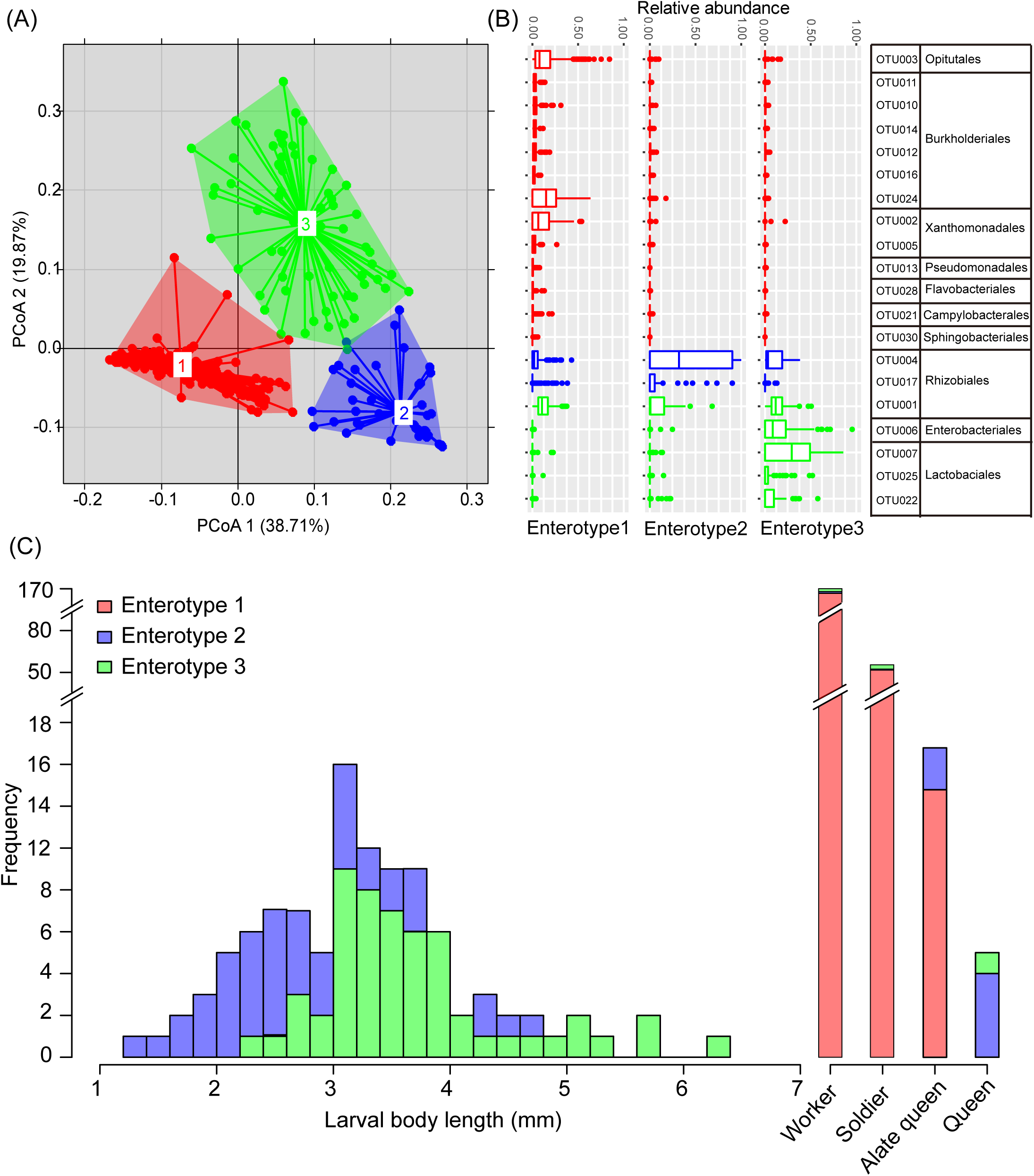
Enterotyping analysis categorizes microbiomes from 13 *Cephalotes* species into three clusters. Sequence libraries clustered at 97% OTU levels, were used to compute weighted UniFrac values. Using the resulting matrix, libraries were clustered with the partitioning around medoids (PAM) algorithm, and the Calinski-Harabasz (CH) index was used to identify the optimal number of clusters as n=3. A) Between-class analysis, performed with the package “ade4” in R, was used for this first plot, which shows similarities of sequence libraries from Enterotypes 1 vs. 2 vs. 3 across two PCoA axes. B) Stem and whisker plots illustrate medians, quartiles, and outliers for each 97% OTU found significant in our indicator OTU analyses. Relative abundances of these OTUs, hence, vary significantly across enterotypes. C) Microbiomes from larvae of varying size (normalized sizes on x-axis) binned, differentially, into enterotypes 2 vs. 3, while microbiomes of varying castes varied, to some degree, in their enterotype assignments.

Enterotype 2 included microbiomes of larvae, four out of five mature/wingless queens, and two out of sixteen alate queens/gynes, in addition to very small numbers of workers and soldiers (Fig. 5c). Characteristic of this cluster were Rhizobiales OTUs 004 and 017 (Fig. 5b). Microbiomes assigned to the third enterotype were mostly from larvae. However, one hailed from a wingless queen/gyne, while a few belonged to soldiers and workers. Rhizobiales OTU001 was identified as an indicator OTU for this enterotype, as was Enterobacteriales OTU006, along with three Lactobacillales OTUs (Fig. 5b**; Supplementary Table 12**). Estimating larval developmental stage through body length measures, we noticed that enterotype 3 was dominated by microbiomes of older (i.e. larger) larval samples, while the mature/wingless queen-including enterotype 2 was primarily comprised of young larvae (Fig. 5c).

### Correlations between host phylogeny and *cephalotine* microbiomes differ across enterotypes

Principal coordinates analysis plots (Fig. 6) suggested that host phylogeny plays a significant role in predicting community structure for ants hosting enterotype 1-clustering microbiomes, but not clearly those clustering with enterotypes 2 or 3. With an interest in understanding whether cephalotine microbiome composition is predicted by evolutionary history, we investigated the impact of host phylogeny and geographic location on community beta-diversity using partial Mantel tests. Bray-Curtis and Jaccard Index dissimilarity matrices computed for enterotype 1-classifying samples, were significantly correlated with the host phylogenetic distance matrix, at 97%, 98%, and 99% OTU clustering widths, and at the oligotype level, after controlling for the effect of geographic distance (P ≤ 0.001; Table 1).

**Figure 6:**
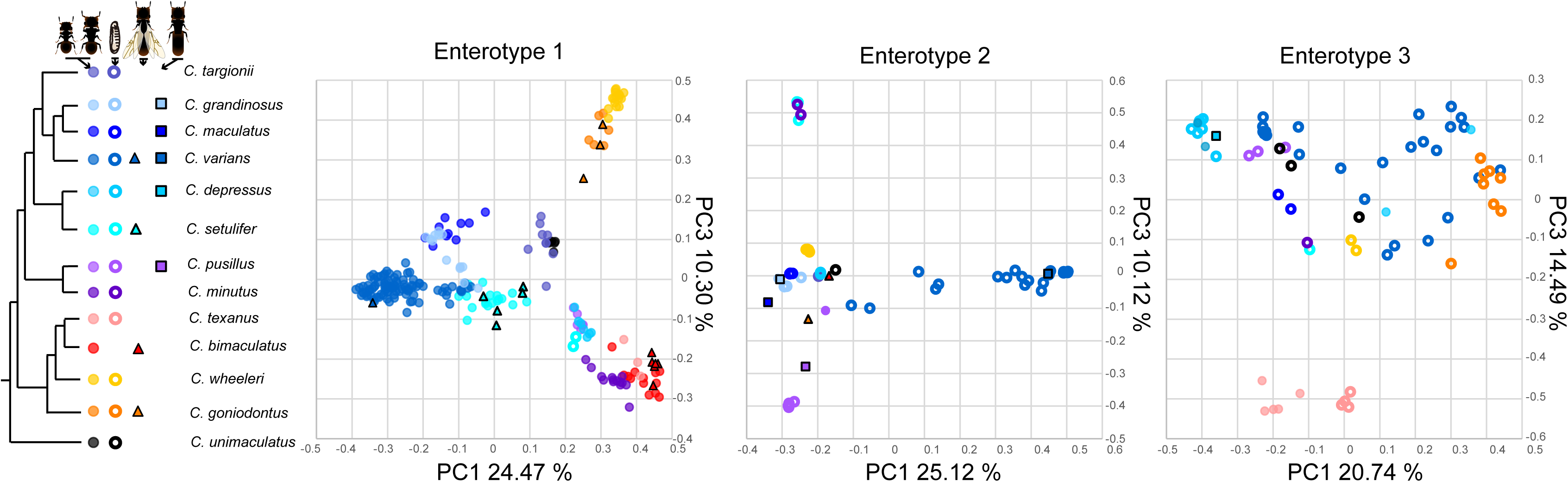
PCoA plots on microbiomes from each enterotype. Data were generated from Bray-Curtis dissimilarity measures computed on our oligotype table, with separate plots and analyses for the libraries assigning to each enterotype. Phylogeny at the left shows host relatedness, and connects different colors and symbols to varying *Cephalotes* species, castes, and developmental stages. Clustering by host relatedness is evident in panel 1, for the first enterotype – a pattern consistent with findings from our partial Mantel test results.

**Table 1.** The r value and p value of Mantel’s test showing the influence of host phylogeny in controling of geography or the influence of geography in controling of host phylogeny for each enterotype.

| Enterotype ID | Effect | Controlling for | 97% OTU |  | 98% OTU |  | 99% OTU |  | Oligotype |  |
| --- | --- | --- | --- | --- | --- | --- | --- | --- | --- | --- |
|  |  |  | Partial R<sup>2</sup> | P value | Partial R<sup>2</sup> | P value | Partial R<sup>2</sup> | P value | Partial R<sup>2</sup> | P value |
| Based on Bray-Curtis distance matrix |  |  |  |  |  |  |  |  |  |  |
| Enterotype1 | Host phylogeny | Geography | 0.457 | 0.001 | 0.515 | 0.001 | 0.496 | 0.001 | 0.519 | 0.001 |
|  | Geography | Host phylogeny | 0.297 | 0.015 | 0.447 | 0.003 | 0.284 | 0.01 | 0.157 | 0.141 |
| Enterotype2 | Host phylogeny | Geography | 0.079 | 0.537 | 0.196 | 0.143 | 0.111 | 0.353 | 0.130 | 0.081 |
|  | Geography | Host phylogeny | 0.132 | 0.220 | 0.429 | 0.005 | 0.170 | 0.151 | 0.176 | 0.15 |
| Enterotype3 | Host phylogeny | Geography | 0.017 | 0.563 | 0.026 | 0.858 | 0.068 | 0.66 | 0.045 | 0.776 |
|  | Geography | Host phylogeny | -0.129 | 0.200 | -0.034 | 0.816 | -0.066 | 0.688 | 0.338 | 0.025 |
| Based on Jaccard distance matrix |  |  |  |  |  |  |  |  |  |  |
| Enterotype1 | Host phylogeny | Geography | 0.646 | 0.001 | 0.676 | 0.001 | 0.656 | 0.001 | 0.636 | 0.001 |
|  | Geography | Host phylogeny | 0.477 | 0.005 | 0.547 | 0.001 | 0.452 | 0.003 | 0.363 | 0.008 |
| Enterotype2 | Host phylogeny | Geography | 0.363 | 0.004 | 0.434 | 0.004 | 0.562 | 0.001 | 0.262 | 0.039 |
|  | Geography | Host phylogeny | 0.033 | 0.801 | 0.252 | 0.035 | 0.048 | 0.69 | 0.131 | 0.328 |
| Enterotype3 | Host phylogeny | Geography | -0.006 | 0.959 | 0.193 | 0.225 | 0.375 | 0.021 | 0.372 | 0.02 |
|  | Geography | Host phylogeny | 0.204 | 0.190 | 0.222 | 0.176 | 0.301 | 0.051 | 0.257 | 0.106 |

Geographic distance explained a smaller proportion of community dissimilarity for enterotype 1. Specifically, when using Bray-Curtis distances the average r-squared value across clustering widths was 0.296 for geography compared versus 0.497 for phylogeny. For Jaccard index-based analyses, the average r-squared value for geography was 0.460 compared to 0.654 for phylogeny. But like phylogeny, geography did show a significant correlation with community dissimilarity at every OTU clustering width, along with the oligotype level, for Jaccard index analyses. Significant correlations between geographic distance and community dissimilarity were evident for 97%, 98%, and 99% OTUs, but not the oligotype level, when analyzing Bray-Curtis distances (P < 0.05; Table 1).

For enterotype 2, host phylogenetic distance was significantly correlated with Jaccard dissimilarity after controlling for geographic distance, at all sequence clustering widths (P < 0.05; Table 1). These correlations were not, however, significant at any level of sequence clustering when using the Bray-Curtis dissimilarity matrix. Average r-squared values for the effect of phylogeny were slightly lower, for Jaccard-based analyses on enterotype 2 (0.405) compared to those on enterotype 1 (0.654). For this second enterotype, correlations between community dissimilarity and phylogeny-controlled geographic distances were only significant at the 98% clustering level.

For enterotype 3, host phylogenetic distance and community dissimilarity were not significantly correlated after accounting for geographic distance, except for a significantly positive correlation for the Jaccard distance computed at the 99% OTU and oligotype levels. Average r-squared values, however, were again comparatively low versus those from enterotype 1 (0.374 vs. 0.646), when averaged across these significant clustering widths. In addition, the correlations between geographic distance and community dissimilarities for enterotype 3 were only significant at the oligtype level for Bray-Curtis measures (Table 1).

We experimented with partial Mantel tests on enterotype 3, repeating analyses with only specialized, adult-enriched bacteria (defined in subsequent sections) or with non-specialized bacteria likely acquired from the environment. Focusing on 99% and oligotype levels, and beta-diversity metrics computed with the Jaccard index, we found that phylogenetic distance remained a significant predictor of microbiome dissimilarity when only specialized bacteria were included. For these results r-squared values of significant findings ranging from 0.335 – 0.502. When only non-specialists were included prior to Jaccard distance matrix calculation, phylogeny was not significant (**Supplementary Table 18**). Combined with results discussed further below, these results suggest a strong effect of host phylogeny on microbiomes made up primarily by specialized *Cephalotes* symbionts (enterotypes 1 & 2), but not on those comprised largely of free-living bacteria (enterotype 3).

### Phylogenetic affinities of abundant bacterial species from *Cephalotes* larvae

To initiate a study on the evolutionary origins of larva-enriched, and in some cases, newly identified *Cephalotes*-associated symbionts, we used maximum likelihood analysis for phylogenetic inference on abundant sequences from the Rhizobiales (Analysis #1), Enterobacteriales (Analysis #2), and Lactobacillales (Analysis #3). Our Rhizobiales phylogeny indicated that 16S rRNA gene sequences from such microbes grouped into two monophyletic, *Cephalotes* ant-specific lineages (Fig. 7a). Symbiont strains from Rhizobiales OTU001 formed a distinct clade, which was sister to a ‘crown group’ clade comprised primarily of symbionts from other ants. Nested within this crown group was a second lineage comprised exclusively of *Cephalotes* associates, including OTUs 004, 017, 023, 042, and 056.

**Figure 7:**
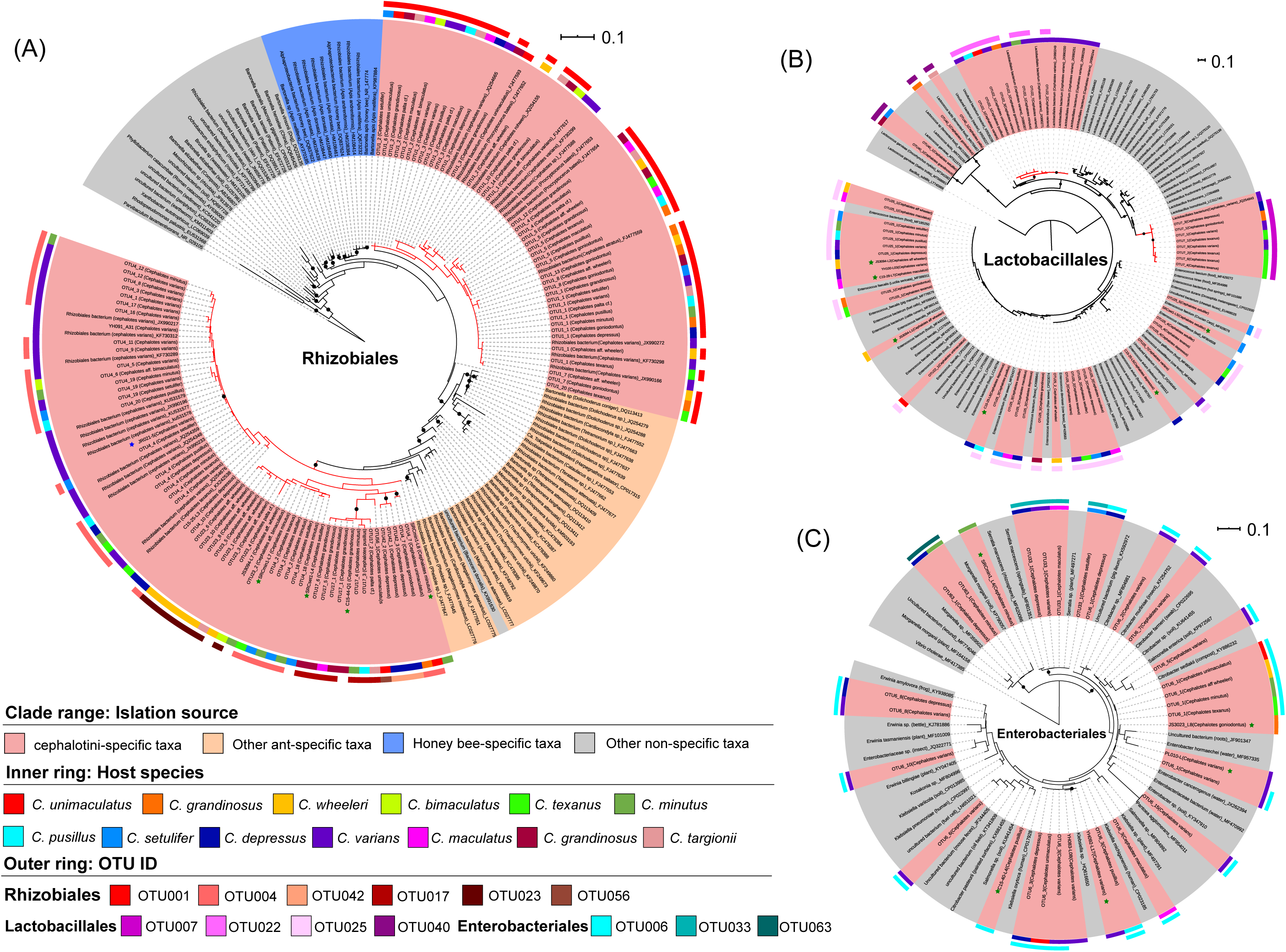
Maximum likelihood phylogenies of three taxa found in abundance in larvae across the *Cephalotes* genus. All phylogenies inferred from alignments of 16S rRNA sequences. Those from amplicon sequencing in this study are indicated with colored shading in the outer color strips, which indicate bacterial taxonomic assignments. Those produced from our Sanger sequencing are highlighted with stars (culture independent sequence from ant = green; sequence from cultured symbiont isolate = blue). The remainer were identified through BLASTn searches against the NCBI nr database. Names of sequences from cephalotine ants are enclosed in pink shading. Those from other ants are in orange. Those from other hosts or habitats are in gray. Interior color strip identifies the cephalotine host species.

Our maximum likelihood phylogeny of Lactobacillales bacteria (Fig. 7b) revealed that 16S rRNA gene sequences from the more common OTU007 (sister to *Lactobacillus homohiochii* and *L. fructivorans*) and OTU022 (sister to two uncultured bacteria) formed two distinct and well-supported *Cephalotes*-specific clades, separated from free-living bacteria by modestly long branches. While dominant in *Cephalotes* with enterotype 3-like microbiomes, bacteria from these larvae-dominant lineages were found in 23 of the 24 adult ants harboring Lactobacillales. Symbiotic strains from OTU025 Lactobacillales, also common in larvae, but seemingly absent from adults, did not form a host-specific clade, instead showing close relatedness to free-living *Enterococcus* bacteria.

In contrast to the patterns for Rhizobiales and Lactobacillales, 16S rRNA gene sequences from three Enterobacteriales OTUs, dominant in *Cephalotes* larvae with enterotype 3-like microbiomes, did not form host-specific clades, grouping instead with a range of free-living bacteria from this order (Fig. 7c). In particular, sequences from OTU006 showed relatedness to *Erwinia*, *Enterobacter*, *Salmonella*, *Pantoea*, and *Citrobacter.* OTU033 showed phylogenetic affinity to *Serratia*. The more distantly related OTU063 appeared related to *Morganella*.

### Identifying host-specialized vs. environmentally acquired bacteria – BLAST and phylogenetic analyses

Frequent relatedness to free-living, non-cephalotine associated bacteria suggested that several of the bacteria common within larval microbiome are environmentally acquired, unlike the vast majority of microbes from adults. To identify and compare the fractions of stage-enriched microbiomes that are likely acquired from the environment (i.e. highly related to free-living bacteria) vs. seemingly specialized and transmitted through evolved mechanisms for ancient timespans (e.g. Lanan et al. [41]), we began by performing BLASTn searches on the most common (“representative”) sequence found in each of the 154 97% OTUs exceeding pre-defined thresholds (i.e. 0.00294 in at least 2 libraries, or 0.05 in one; **Supplementary Table 19**). Fifty-one of these yielded a top hit to a bacterium previously determined to fall within a cephalotine-specific clade [7, 62, 66]. Across these instances the average sequence identity was 98.0%.

Of the remaining 103 OTUs we performed phylogenetic analysis on 67 of the most abundant (**Figs. S6-S10**), using inclusion criteria, and BLASTn strategies for identifying close relatives that are described in the supplementary figure legends. Analyses from the Actinobacteria (**Supplementary Figure 6**) – found commonly in larvae (**Supplementary Figure 3**) – revealed that OTU062 was related to previously identified gut- and head-associated sequences from *Cephalotes*. Showing some relatedness to an uncultured bacterium from rainwater, and more distant relatedness to *Agromyces* and *Curtobacterium*, we do not yet see strong evidence that this lineage is a long-term specialist given that its only identified cephalotine host is, thus far, *C. varians* from the Florida Keys. Furthermore, its 98.4% sequence identity to a non-cephalotine bacterium (accession #: KY874657) puts it at odds with most cephalotine-specific bacteria, previously shown to exhibit an average of 93.3% relatedness to their closest non-cephalotine match [65]. Only two other monophyletic clusters of cephalotine ant-derived sequences were observed on this tree. While both were distributed across multiple *Cephalotes* species, they were also comprised of only two unique sequences, showing close relatedness to bacteria from other habitats. These trends argue against high likelihood of ancient and specialized relationships between cephalotines and Actinobacteria, once again contrasting with results for most adult-enriched bacteria [7, 62, 66].

Using similar logic, we found minimal evidence for additional cephalotine-specialization among the Lactobacillales (**Supplementary Figure 7**). Beyond the potentially specialized OTUs 007 and 022, only one other cephalotine-specific clade was detected. Consisting of two sequences from OTU085 with 98.81% relatedness to non-cephalotine bacteria (accession #: AB934560.1), this is also unlikely to be an ancient or widespread specialist.

In contrast, the three unique sequences from the ambiguously classifying Pseudomonadales OTU0046, were nested within a cephalotine-specific lineage, albeit with low bootstrap support (62%; **Supplementary Figure 8**). Hailing from *C. wheeleri* and *C. goniodontus*, they showed relatedness to symbionts from three other *Cephalotes* species, including *C. setulifer*, *C. rohweri*, and *C. atratus*. OTU116, from the Bacteroidetes had only 85.4% identity to its top cephalotine-derived BLAST hit. Initially implicated as a plausible specialist, our phylogenetic analyses did place it near cephalotine-specific clades of bacteria in the Sphingobacteriales. But it was not part of a host-specific monophyletic unit, grouping instead with rumen-associated bacteria (**Supplementary Figure 9**).

The phylogeny inferred for Analysis #8 grouped sequences from three Flavobacteriales OTUs (60, 72, and 119) into a monophyletic clade with 85% bootstrap support (**Supplementary Figure 10**). Collectively found in four *Cephalotes* species (**Supplementary Figure 3**), and with members showing only ∼92.0% identity to their nearest non-cephalotine relatives (**Supplementary Table 18**), this lineage can be plausibly retained as a cephalotine-specialized clade. Sequences from the occasionally abundant Neisseriales in OTUs 019 and 069 were included in this same phylogeny. While only 94.5% identical to other, non-cephalotine derived bacteria (**Supplementary Table 18**), we did not see strong evidence for specialization due to non-monophyly of these two OTUs and confinement of each to a single host species (**Supplementary Figure 3**). Also of interest were sequences from the Oxalobacteriaceae, including OUT041, found in three monophyletic Mexican *Cephalotes*, and OTU073, present in the early-branching *C. unimaculatus* (Dominican Republic). While related, monophyly of these sequences was interrupted by *Olivibacterium flavum*, a bacterium from a non-cephalotine source. We saw no additional evidence for host-specific lineages on this phylogeny.

### Proportions of adult vs. larval microbiomes comprised by host-specialist bacteria

Using our phylogenetic results we expanded our list of cephalotine-specialized symbionts to include Lactobacillales (OTUs 007 and 022), Pseudomonadales (OTU046), and Flavobacteriales (OTUs 060, 072, 119) OTUs. With this finalized classification scheme in hand (**Supplementary Table 19**), we computed the proportion of each individual ants’ microbiome comprised by specialized bacteria, assessing these values across stages and castes from each of our 18 studied colonies (**Supplementary Figure 11**). In adults, microbiomes of pre- and non-reproductive castes contained similarly high fractions of bacteria from cephalotine-specialized clades, i.e. those found exclusively in *Cephalotes* and/or *Procryptocerus* ants. Specifically, the median summed relative abundances for such specialized bacteria were, 0.984, 0.983 and 0.987 across across alate queens/gynes, soldiers, and workers, respectively. Averages for alate queens trended slightly downward compared to the other two castes (0.869 AQ, 0.940 S, and 0.956 W) due to the high prevalence of putatively non-specialized *Wolbachia* in the former caste for *C. goniodontus*.

In contrast to these patterns, symbionts from cephalotine-specialist clades comprised a median of 0.469 of mature/wingless queens’ microbiomes, and an average of 0.661. These lower fractions were driven by an abundance of *Wolbachia* or Neisseriales in a subset of these n=5 samples, and by the paucity or absence of specialized symbiont OTUs in the Xanthomonadales, Burkholderiales, Pseudomonadales (i.e. assigning to the genus *Ventosimonas*), Campylobacterales, Flavobacteriales, Gracillibacteria, and Sphingobacteriales. Along similar lines, *Cephaloticoccus* symbionts from the Opitutales – typically the most dominant microbes across adult ants – were abundant in only one mature queen. Indeed, in focusing on the non-Rhizobiales portion of wingless queens’ specialized core microbiomes, we saw that these, and the few other, remaining specialists made up a median of 0 and an average of 0.109 of their microbiomes. Proportions comprised by these non-Rhizobiales specialized bacteria were, contrastingly, high for alate queens/gynes, soldiers, and workers. In particular, these respective averages equaled 0.756, 0.717, and 0.719, illustrating their less Rhizobiales-dominated composition. It is interesting to note that most Rhizobiales found in mature/wingless queens hailed from only one of the two deep-branching cephalotine-associated lineages in this order (Fig. 7), with an absence of OTU001, found in all other castes and stages of every colony in our study (**Supplementary Figure 3**).

Bearing some resemblance to wingless queens, the fractions of the larval microbiome made up by bacteria from cephalotine-specialist lineages was considerably lower than that for alate queens/gynes, soldiers, and workers, with a median and average of 0.700 and 0.675. Removal of Rhizobiales from consideration pushed these values down to 0.162 and 0.206. Additional removal of Lactobacillales OTUs 007 and 022 yielded median and average values of 0.023 and 0.060 for the specialized fraction of the larval microbiome, suggesting the rarity of other core adult-enriched specialist symbionts at this immature stage. Of notable exception were five colonies where the remaining specialized, adult-enriched symbionts comprised over 0.070 of the symbiotic community. In these cases we found a tendency toward modest abundance of specialized Burkholderiales OTUs from the Alcaligenaceae, which comprised between 0.061 to 0.279 of the total microbiome. Specialized Xanthomonales were generally rare, though they did constitute an average relative abundance of 0.065 in one colony from *C. setulifer*. Non-specialized portions of the larval microbiome were most often made up of Lactobacillales from beyond OTUs 007 and 022, Enterobacteriales, and sporadically abundant bacteria, including those from the orders Actinomycetales and Bifidobacteriales in the Actinobacteria (**Supplementary Figure 3**). Bacteria from this phylum made up an average of 0.019 of the larval microbiome, reaching a maximum average of 0.121 in larvae of *C. maculatus*.

### Assessing microbiome variation across larval development

With: 1) established definitions for specialist bacterial lineages (**Supplementary Table 19**); 2) identified patterns of high larval abundance of non-specialized symbionts from the Actinobacteria, Enterobacteriales, and Lactobacillales (**Supplementary Figure 3**); 3) discovery of two divergent specialized Rhizobiales linages (Fig. 7a); and 4) suggestions that microbiomes of young vs. old larvae often classify into distinct enterotypes (Fig. 5c), we assessed trends of symbiont shifts across larval development by binning individuals into 11 normalized size categories, used to infer the relative age of 103 larvae. ANOVA statistics, focused on arcsin transformed relative abundances from each larval sequence library (**Supplementary Tables S20-21**) revealed significant differences in the titers of Rhizobiales OTU001 across larval development (df=10, sum of squares=0.42865, F-value=2.0602, p=0.03573), with a trend of increasing relative abundance with increasing larval size – used here as our proxy for age (**Supplementary Figure 12**). Results were highly significant for the second cephalotine-specific Rhizobiales lineage containing OTUs 004, 017, 023, 042, and 056 (df=10, sum of squares=7.9636, F-value=5.5194, p-value=2.15 e-06); but in this case, relative abundance decreased with size. A combined pool of specialized and non-specialized Lactobacillales showed a trend of greater prevalence in larger and, hence, presumably older larvae, with similarly strong statistical support for a relationship between size and titer (df=10, sum of squares = 2.2033, F-value=3.6587, p-value=3.87 e-04). While non-specialized Enterobacteriales showed a similar trend of increasing relative abundance with size, larval size categories did not have a significant effect in our ANOVA model (df=10, sum of squares=0.7507, F-value=1.63, p-value=0.1103), suggesting a need for future study on this taxon. No other bacterial groups considered here showed significant differences in arcsin transformed relative abundance across these ranked size classes (**Supplementary Table 21**).

### Cophylogeny analysis – *Cephalotes* species vs. *Cephaloticoccus symbionts*

Finally, to assess whether the earlier described phylosymbiosis signal might extend from cospeciation between turtle ants and their symbionts, we initiated what will be a deeper, more systematic investigation into host-symbiont phylogenetic congruence (Fig. 8; **Supplementary Figure 13**). For this purpose, we used sequence data from a prior metagenomic study [7], inferring phylogenies of adult-enriched *Cephaloticoccus* (Opitutales) symbionts from separate analyses of two non-laterally transferred protein-coding genes. Bootstrap support vales for the nodes in each phylogeny were generally high, exceeding 80 for 10/16 internal nodes of the *uvrB* tree and for 9/16 internal nodes of the *rpoB* tree.

**Figure 8:**
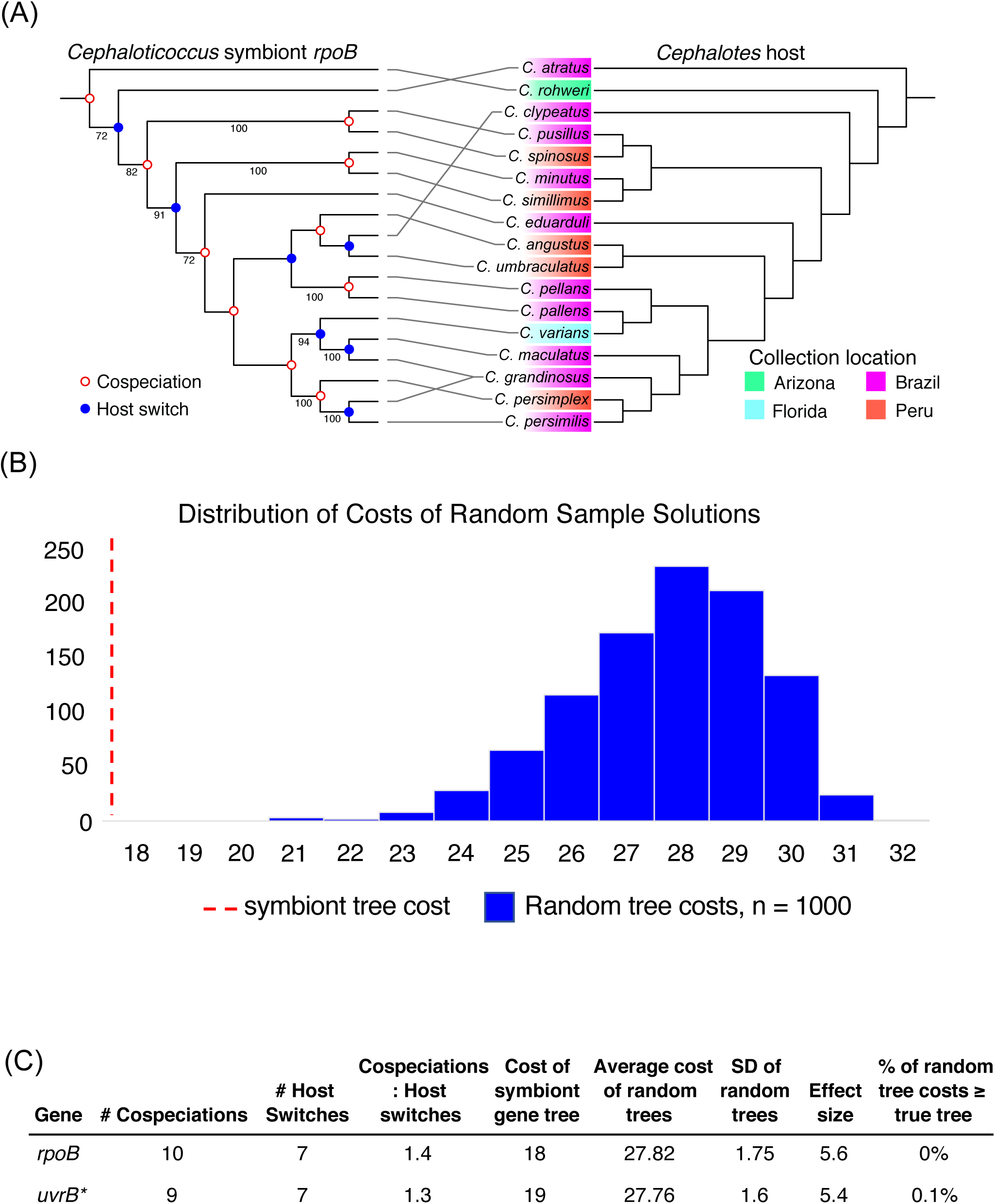
Cophylogeny analyses for *Cephaloticoccus* symbionts. A. Tanglegram of *rpoB* symbiont gene tree (from *Cephaloticoccus* symbionts extracted from metagenomes of Hu et al. [7]) vs. *Cephalotes* host tree (from Price et al. [93]). For the symbiont gene tree, we list all bootstrap values ≥70. Cospeciation and host switching events are indicated at each symbiont node and were inferred based on the top solution from Jane4. B. Cost histogram from 1000 randomized tip mapping permutations in Jane4. A lower cost indicates fewer inferred host switching events, and therefore stronger phylogenetic congruence. C. Table of Jane4 analysis statistics. *The *uvrB* tanglegram and histogram are available in **Supplementary Figure 12**.

We compared these two phylogenies against the host *Cephalotes* phylogeny inferred by Price et al. [93], using Jane4. Both symbiont gene trees exhibited a greater degree of phylogenetic congruence in relation to the host phylogeny than expected by chance (Fig. 8**; Supplementary Figure 13**). In particular, the event-based mapping of the Jane4 program estimated 10 cospeciation events compared to 7 host switches for the *rpoB* vs. host tree comparison. Additionally, the overall ‘cost’ of our inferred *rpoB* tree – 18 – was substantially lower than the all costs estimated for 1000 randomized datasets (average randomized cost = 27.82). Results were similar for the *uvrB* tree – with 9 cospeciation events compared to 7 host switching events, and an actual cost of 19 compared to an average of 27.74 for randomized datasets. Among the several nodes suggestive of cospeciation events, two pairs of sister symbiont taxa recapitulated sister taxa relationships on the *Cephalotes* phylogeny in ways that were not confounded by geography. This was true for both gene trees. Specifically, symbionts from *C. pusillus* (Brazil) and *C. spinosus* (Peru) grouped together with high bootstrap support (*uvrB* = 95; *rpoB* = 100), as did symbionts from *C. minutus* (Brazil) and *C. simillimus* (Peru) (*uvrB* = 80; *rpoB* = 100). In a distinct instance, the sister pairing of Peru-collected *C. angustus* and *C. umbraculatus* was nearly recapitulated on the *rpoB* gene tree, with the symbiont clade’s monophyly being broken up only by an apparent horizontal transfer from this symbiont lineage to the Brazil-derived *C. clypeatus.* Collectively, these patterns suggest that while the symbionts used for this analysis came from ants in a small number of geographic locales, host evolutionary history appears to be a geographically robust predictor of symbiont gene tree topology for at least one cephalotine-specific, adult-enriched symbiont.

## Discussion

Motivated by prior findings of microbiome shifts across holometabolous insect development [31]; by discoveries and expectations that some social insect castes do, or should, harbor microbes with divergent function [105]; by the recent discoveries of the actual and potential functions of turtle ant gut microbiomes [7, 70, 73, 106]; and by our understanding of mechanisms promoting beneficial symbioses [26], we set out to establish a comprehensive, colony-level view of the turtle ant microbiome across the *Cephalotes* phylogeny. Expanding upon foundational work on these social, holometabolous insects, our study provides several insights with broader relevance for understanding microbiomes of animals.

We observe that workers and soldiers harbor similar microbiomes, with a few exceptions, and that workers may undergo microbiome stabilization during early phases of adulthood (Fig. 3). We also find that microbiomes of winged queens (or “gynes”) resemble those of workers and soldiers, with the vast majority of their microbes coming from cephalotine-specific clades (**Supplementary Figure 3**). Differing from prior work in social hymenopterans like honeybees [51–53], this discovery suggests gynes acquire the parent colony’s microbiome prior to dispersal and mating – indicating a likely means for transgenerational symbiont passage and partner fidelity (Fig. 9). But intriguingly, mature, wingless queens from established colonies possessed simple microbiomes comprised of Rhizobiales bacteria abundant in young larvae. This, in turn, suggests a process of microbiome succession – or, more specifically, winnowing – sometime after colony founding (Fig. 9). Future studies on young colonies will be necessary to further understand this hypothesized transition.

**Figure 9:**
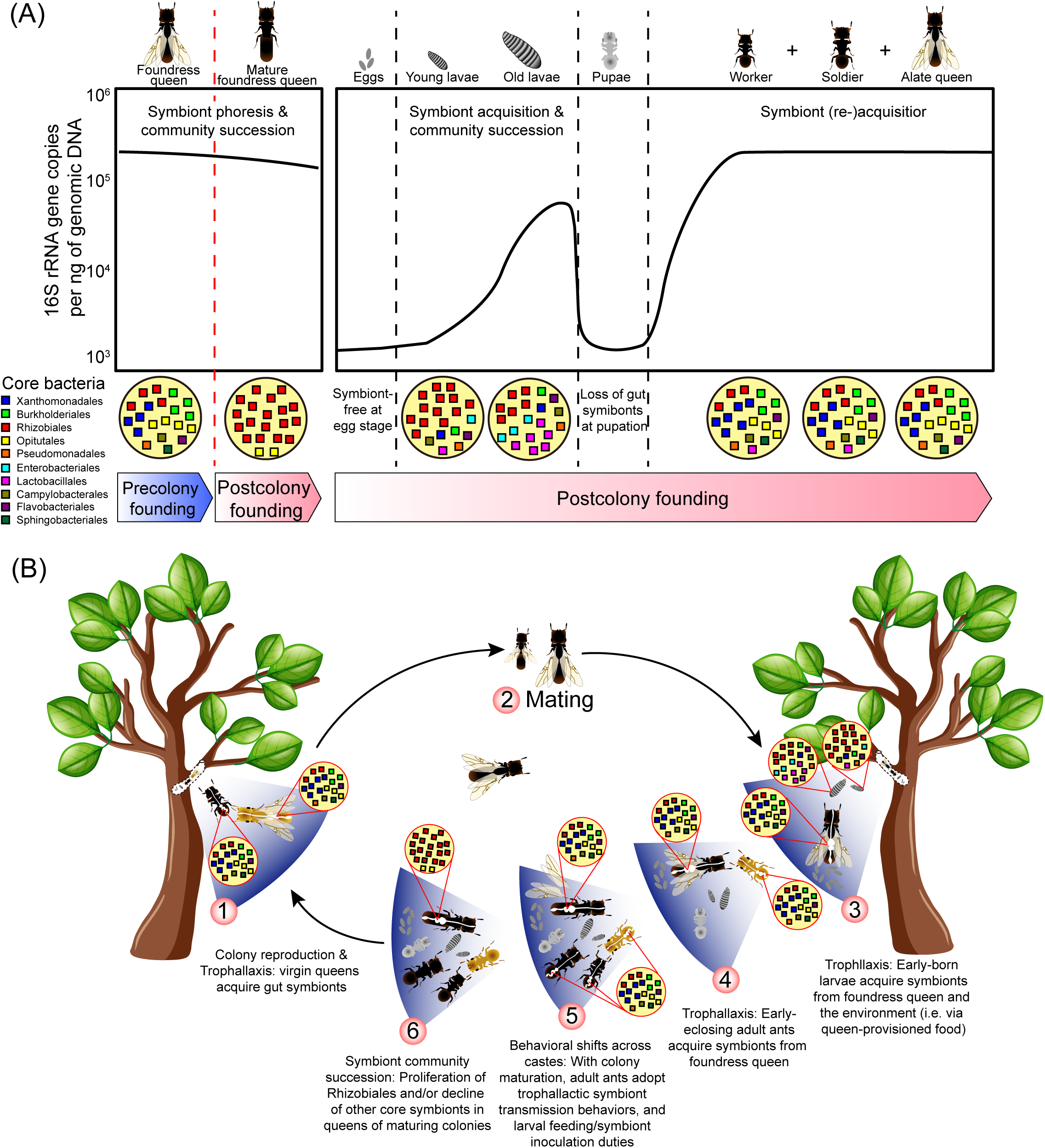
Summary figure, illustrating mechanisms of microbiome acquisition, community succession, and the impacts of caste and development. A) Symbiot dynamics over the lifespan of an individual al ant across its development with the adult-stage symbiont transition proposed in queens at left, and the typical progression for all ants, regardless of caste, on the right. B) Hypothesized lifecycle of the microbiome in relation to the *Cephalotes* ant colony lifecycle. Process begins with production of winged (alate) queens, which mate on the wing with males, and found new colonies – often individually. Presence of worker and soldier like microbiome, known to be transmitted through oral-anal trophallaxis (and “walled-in” by a proventricular filter) in alate queens implies that they bring the specialist microbiomes with them upon mating and new colony founding, which would indicate a route for vertical symbiont transfer. Young queens would then need to be the inoculation sources for the specialist microbiomes (e.g. Rhizobiales in larvae; and the diverse range of OTUs in adults) in early-born larvae and adults. As colonies mature, workers and soldiers would take on trophallactic inoculation roles. Queens are then hypothesized to undergo a microbiome succession toward a lower-diversity, often Rhizobiales-enriched community. Bacteria from non-cephalotine specialized taxa are typically very closely related to free-living microbes. It is proposed that these are selectively acquired, and retained, by larvae from food.

As for adults, we find larval microbiomes to be highly conserved across 40 million years of *Cephalotes* evolution. But larval and adult microbiomes differ in both quantitative and qualitative regards. While both stages begin – at egg hatching and eclosion – with individuals harboring very few bacteria (Fig. 2), the Rhizobiales symbionts dominating the guts of young larvae comprise just one of the nearly one dozen adult-enriched symbiont clades found in most adults past the early callow stage (Figs. 3, 4). As larvae grow, however, their microbiomes undergo a stereotyped pattern of community succession, as inferred from our discovery that late-stage (i.e. large) larvae harbor a mixture of adult-enriched specialists, and bacteria of likely environmental origin (Figs. 3, 5, 7).

Our statistical analyses indicate that similarity of the adult microbiome among *Cephalotes* species (Fig. 6) recapitulates host relatedness (Table 1), with at least one symbiont exhibiting partial cospeciation (Fig. 8) with its cephalotine hosts. Host phylogeny does a poorer job of predicting microbiome similarity in older larvae (Table 1), a reflection of the conserved nature of this microbiome and of a likely lack of coevolution between turtle ants and late-larva-dominant, free-living bacteria. As discussed at the end of this section, we conclude that the highly conserved symbiont compositions of larval and adult cephalotine ants have been governed by distinct mechanisms – partner fidelity and environmental filtering – with differing relative impacts across development.

### The impacts of stage on holometabolous insect microbiomes

Symbiotic gut communities of larvae and adults are expected to differ for most holometabolous insects (Hammer & Moran 2019). This stems not only from the distinct ecology of larvae and adults, but also from the blank slate gut environments that often occur upon egg hatching and pupal eclosion (e.g. Fig. 2; Roche and Wheeler [71]; Moll et al. [32]; Wang and Rozen [107]). Opposing forces, however, may partially homogenize microbiomes harbored on opposite sides of metamorphosis, including maternal symbiont deposition upon eggs in their pre-hatched stage; larval development in habitats seeded by adult microbes; social transfer; developmental persistence of symbiont-housing tissues and their contained symbionts; symbiont migration and tissue recolonization within pupating hosts; and insect consumption of symbiont-housing body contents shed prior to pupation [16, 37, 107–109].

While our prior findings suggested that turtle ant adults and larvae share some specialized symbionts [65], we show here that the microbiomes from these cephalotine stages exhibit only partial overlap. In particular, after removal of likely cross-contaminants, we estimate that an average of 31.5 bacterial 97% OTUs are housed, per colony, when assessed among workers and larvae for the 12 *Cephalotes* species censused at both stages (**Supplementary Table 22**). Among these, 10.8 OTUs are shared; 7.9 are found at detectable levels only in workers; while the remaining 12.8 OTUs are confined to larvae.

This discovery resembles findings from other holometabolous insects, including the emerald ash borer beetle. In this insect, 13 of 20 bacteria detected in at least 2 larval and/or adult samples were shared across these stages, which differentially feed on the cambium and phloem, versus leaves, of ash trees [110]. Like the emerald ash borer, substantial proportions of larval midgut bacteria are harbored in the guts of adult forest cockchafers, pest scarab beetles exhibiting distinct root- and leaf-feeding habits as larvae and adults [111]. Fitting a similar mold is the butterfly *Heliconius erato*. Amplicon sequencing of the 16S rRNA gene from this species showed that three of the top 10 bacterial phylotypes were shared across adults and larvae, despite adult consumption of pollen and plant nectar compared to the leaf feeding habits of larvae [34].

Contrasting with these findings are discoveries from the dipteran *Drosophila melanogaster*, in which five bacterial species comprising ≥90% of most sampled microbiomes were each shared across adults and larvae in one lab-based study [36]. It has been traditionally thought that gut microbes of *Drosophila* do not establish residency, and that they are, instead, transient members of the fly diet that are (re-)seeded in food after gut passage [112, 113]. Work here, contrastingly argues that juvenile-associated turtle ant symbionts associate with larval gut tissue (**Supplementary Figure 2)**. And by virtue of their residence in a blind gut, disconnected from the larval anus, the symbionts of turtle ant larvae cannot be viewed as rapid, passthrough transients.

In a slightly different vein, recent research on *D. melanogaster* demonstrated a capacity for at least a small fraction of their symbionts to stably reside within the adult crop. The stable crop resident identified in that study, *Acetobacter thailandicus*, increased larval fitness, being acquired by larvae after inoculation into food substrate from its adult crop staging ground [109]. This supports a broader trend in which holometabolous insect adults carry and vector microbes of benefit to their young offspring, despite the possible lack of direct benefits to the adults, themselves (e.g. Kaltenpoth et al. [114]; Shukla et al. [115]). In the *Cephalotes* system, it is certainly possible that a subset of the specialized symbionts found in alate queens, soldiers, or workers do not directly benefit their carriers, but that they are helpful, instead, to larvae. Benefits to larvae may unfold through inoculation of symbionts with activity in larval guts, or through trophallactic feeding of symbiont-derived metabolites produced in adult “nutrient factories”.

Among other social insects, like *Cephalotes*, there are likely numerous opportunities for cross-stage symbiont exchange given the close proximity of larvae and adults, and the capacities for trophallactic, social transmission. Accordingly, partial overlap between adult and larval gut microbiomes has been reported for honeybees and bumblebees [116]. Studies have shown variable presence of adult-enriched symbionts, like *Gilliamella* and *Snodgrassella*, in larvae [40, 117–119]. But consistent among investigations are observations that larval bacterial loads are lower than those of adults, with some evidence for an increase in community size with larval age [40, 118, 119], and for parallel succession in larval gut communities [117, 118], with both patterns resembling those reported here for *Cephalotes* (Figs. 2; 3).

Among other bacteria in honeybees, several investigations have uncovered an abundance of the adult-associated “Alpha 2.2” symbiont in larvae [40], which has now been formally named *Parasaccharibacter apium* [120]. One study, for example, found this bacterium to dominate the cultivable fraction of gut bacteria in young larvae [117]. Having been detected in conditioned and stored pollen in earlier work [40], *P. apium* was subsequently shown to be present in royal jelly, and the hypopharyngeal glands and crops of nurses, indicating a flexible suite of lifestyles and the likely concentration of this microbe in sources of larval inoculation. This may be key to successful early development, as some *P. apium* strains boost larval fitness [120]. This species, furthermore, is found in mature and possibly virgin queens, but is, contrastingly, rare in workers [51, 119].

### The impacts of caste on social (holometabolous) insect microbiomes

Distributional trends of *P. apium* comprise part of a broader trend of caste-correlated microbiome differentiation in honeybees [52, 121, 122]. Caste differences have been demonstrated or predicted in other social insect systems, though this topic remains somewhat unexplored [105]. We observed subtle symbiont differences among cephalotine workers and soldiers, including tendencies toward specialist Campylobacterales enrichment in workers, and episodic enrichment of Neisseriales and Lactobacillales OTUs in soldiers. These latter patterns, however, unfolded in only 1-2 turtle ant species. Notwithstanding the below-discussed microbiome divergence in mature queens, the absence of consistent microbiome differences among other adult castes of *Cephalotes* raises interesting questions.

To date, the roles of the turtle ant soldiers – a derived caste not present in ancestral *Cephalotes* – appears linked to ecological specialization in nesting habits, with soldier head size being a finely selected trait linked to the use of tree nest cavities with entrances of a specific diameter [123, 124]. While this distinction does not lead to *a priori* expectations of inter-caste microbiome differences, further study on the roles of soldiers, versus workers, in inoculating larvae or young adults with core symbionts [41, 69] could shed light on the importance of soldier microbiomes. Also of importance will be an understanding on the direct impacts of symbionts on the fitness of hosts, or indirect benefits received through boosts to the colony’s nutrient economy or defensive status. Metatranscriptome or metabolomics studies would enable complementary lines of inquiry into realized, *in vivo* symbiont function across castes. Indeed, gut metabolome analysis in honeybees, across a series of bacterial manipulations, has demonstrated contributions of gut bacteria to the metabolism of several components of the pollen diet [46]. These capacities are intriguing in light of slight variation in microbiomes among honeybee foragers and nurses [122], and given also that, among honeybee castes, foragers consume less pollen, and more nectar, than nurses [125].

### Symbiont function and implications for variable symbioses within colonies

Gut microbiomes of insects can improve host fitness through a variety of mechanisms, ranging from detoxification [126, 127], to defense [4], and to a series of nutrient-themed functions. These latter benefits include increased digestive efficiency [128–132], improved nitrogen economies achieved through the fixation or recycling of nitrogen [133–135], and the provisioning of essential, limiting nutrients like amino acids and B-vitamins [136–138].

In turtle ants, members of the core, specialized adult microbiome have been shown to recycle waste nitrogen, with worker ant hosts obtaining large quantities of this recycled nitrogen in the form of amino acids [7]. Through a more recent metagenome-focused effort, a broader suite of likely functions has been identified for adult-enriched specialist bacteria, and for those enriched in larvae [73]. Discovered among adult-associated symbionts, in addition to the aforementioned N-metabolic functions, were capacities for B-vitamin synthesis and breakdown of recalcitrant plant cell wall fibers, including xylan, some forms of pectin, and possibly lignin. Among larvae, Enterobacteriales and Lactobacillales – likely originating from the environment – encode similar functions, in addition to capacities to catabolize cellulose, and a seemingly greater capacity to metabolize pectin, found in abundance in pollen cell walls. It is, thus, highly plausible that turtle ant gut symbionts help to digest fiber-rich diets, providing usable forms of carbon to the colony. The fact that candidate digestive symbionts of larvae live in other habitats (Fig. 7), suggests a need to demonstrate the expression of key genes, and the resulting impacts on host nutrient/energy budgets, through further study.

This functional study [73] begins to provide insight into how various symbiotic functions could diverge or overlap across adults and larvae. Larvae are often viewed as the colony’s digestive caste (e.g. Erthal et al. [57]), at least in ant species with a narrow proventriculus [139]. As such, an abundance of plant cell wall degrading bacteria in pollen-feeding ants like *Cephalotes* could improve the efficiency of plant fiber digestion, while providing increased access to the nutrients sequestered behind the pollen cell wall. A range of larval gut bacteria also encode fermentative capacities, including the core, specialist symbionts from the Rhizobiales that are dominant in young larvae [73]. With an additional potential to synthesize vitamin B12 (cobalamin), it is plausible that this symbiont could benefit larvae from early through late stages.

Nitrogen-recycling, and amino acid biosynthetic capacities of larval symbionts could be of further importance in ant development. With ant larvae requiring more protein than adults [72, 140, 141], the question arises as to why worker-associated symbionts invest in N-recycling and amino acid synthesis [7]. While we cannot rule out impacts on queen reproduction or direct aspects of worker fitness, it is certainly possible that these functions provide a greater direct benefit to larvae. Prior studies, for example, suggest that storage proteins can accumulate in workers, and that these protein levels are depleted in the presence of larvae [142, 143]. Through trophallaxis, and regulated worker physiology, it is thus conceivable that a fraction of symbiont-fueled N-metabolism in workers [7] supports the larval N-economy. Whether the digestive, fermentative abilities of larval symbionts are of symmetrical use to the energy budgets of adult workers (e.g. Grover et al. [144]) is an intriguing possibility awaiting similar investigation.

Also worthwhile will be investigations into the function of the low-diversity microbiomes of mature turtle ant queens. In a recent study, symbionts of *Temnothorax nylanderi* ant queens were found to impact egg production [55]. Might mature queens’ gut symbionts directly sustain reproduction in turtle ants? In addressing this, it is of interest to note that a bacterium with close relatedness to mature queen-enriched Rhizobiales symbionts has had its genome sequenced [7]. Interestingly, this symbiont – strain JR021-5 – lacks abilities to recycle nitrogen and to synthesize most amino acids, as does a close relative sampled recently through metagenomics [73]. The absence of extensive N-metabolism does not rule out a supportive role for queen reproduction, but given the longevity of ant queens and the known N-requirements to support female insect reproductive function [145], this finding does at least begin to question this symbiont’s importance in reproductive support.

But the impacts of Rhizobiales symbionts on turtle ant queens could alternatively be realized through protective, rather than nutritive or energetic functions. In *Temnothorax* ants, patterns of queen gene expression shift from enrichment in anti-pathogen/parasite defensive function to investment in antioxidant activity, suggesting two protective functions that symbionts might fulfill [146]. Oxidative damage is of further importance to aging queens of honeybees, with evidence for a series of evolved host- and, possibly, microbe-mediated mechanisms to mitigate such damage [52]. Might Rhizobiales protect queens from senescence-inducing oxidative damage? According to a recent publication, such a possibility is not far-fetched [147], as at least some turtle ant Rhizobiales were seen to encode the biosynthetic machinery to make arylpolyenes – carotenoid-like molecules protecting against oxidative damage [148].

Up-regulated immune defense would also be advantageous for ant queens that have long life spans, inhabit fixed nests, and establish colonies independently in pathogen-rich habitats after mating [149, 150]. And, indeed, higher pathogen resistance in *Lasius* and *Formica* queens, after mating, suggests an eventual up-regulation of protective mechanisms during the process of queen maturation and aging [150]. Building off of these ideas, we return to the earlier described honeybee symbiont, *Parasaccharibacter apium.* Notably, this crop-, larval-, and queen-abundant symbiont improves colony-level resistance against microsporidian *Nosema* parasites [151]. Strong parallels to the distributions of JR021-5-like Rhizobiales symbionts, as shown here and elsewhere [69, 74], raise questions as to whether defense could be similarly enacted by these cephalotine ant symbionts. Indeed, might selection favor symbiont-encoded defense in mature queens after they have nourished early-born offspring and inoculate first-borns with core symbionts (Fig. 9)? And might defensive functions be crucial in protecting microbe-free young larvae? Addressing these questions will be of clear importance in the further development of a colony-level view for turtle ant symbioses and symbiont function.

### Symbiont conservation and mechanisms promoting beneficial microbiomes

While symbiosis is an important part of eukaryotic biology [152], we have intriguingly learned much on its significance from studying ‘exceptions to the rules’ – including cases in which symbionts are harbored at low densities [153] or those in which they have been lost, or replaced, after ancient associations [154]. Emerging from such work is the discovery that highly integrated and ancient symbioses can be hard to escape [155, 156]; that fixes to old symbioses may sometimes involve slap-dash solutions [157]; and that the origins of novel resource acquisition or defensive strategies [158, 159], or the acquisition of key genes via lateral gene transfer [160], can eliminate the need for symbionts or at least some symbiont functions.

In light of this, and observations that microbes are harbored at low densities across many ants [81, 161–163], the cephalotine-gut microbe symbiosis comes across as fairly conspicuous. With the *Cephalotes* genus dating in excess of 46 million years, the conservation of symbionts in adults and larvae is noteworthy – all the moreso in the face of habitat and caste differentiation exhibited across this clade [93]. Enrichment in symbiotic gut bacteria, harbored at high abundance, is reported to be concentrated at the low ends of the food chain among tree-dwelling ants like *Cephalotes* [162], with the internal housing of specialized, symbiotic bacteria appearing as a derived state in many ant lineages (Russell et al. [161]; but see Jackson et al. [164]). A prior study noted a concentration of specialized symbiotic bacteria in ants feeding primarily on combinations of spores, pollen, vertebrate excreta, insect honeydew, extrafloral nectar, and plant wound secretions [65, 161, 165]. Within some of these groups, symbiosis appears ancient. This is true for ants in the Camponotini and their intracellular *Blochmannia* symbionts, with their estimated 40 million years of association history [166].

Extracellular gut microbiomes can show levels of conservation approaching, or exceeding that, of cephalotine ants in other insect systems. This is true, to a degree, for the termites. While the diversity of gut microbiomes in this system has complicated evolutionary histories, these social, hemimetabolous insects have long engaged in interactions with a range of specialized bacteria and protists, with some relationships stretching back in arguable excess of 150 million years [167, 168]. Coming from less diverse bee gut microbiomes are several core symbiont specialists, including *Gilliamella* and *Snodgrassella*, which are argued to have been acquired by social, corbiculate bees nearly 80 million years before present [169]. As such the age of the *Cephalotes* microbiome association is by no means unprecedented, nor is our discovery that symbionts from this group may cospeciate with their hosts (Fig. 8). Hinted at through indirect measures in a prior turtle ant study [63], cospeciation has previously been evidenced for extracellular gut symbioses in bees [48] and termites (e.g. Noda et al. [170]).

The known means of social symbiont transmission in these above systems [40, 42, 48] argue, in part, for microbiome conservation through partner fidelity (e.g. Kaltenpoth et al. [13]). And in our study, the presence of the core microbiome in pre-mated queens suggests a means by which a parent colony can propagate its microbes to the next generation (Fig. 9). Over long periods of time, such transmission should select for beneficial microbes [10].

Such a process has unfolded for numerous, transovarially transferred intracellular symbionts, which have evolved streamlined genomes uniquely suited to meet host needs [171]. It has also occurred for a range of extracellular symbionts [13, 172, 173], including farmed symbionts not housed within or on the host body. Perhaps most famous among such latter associations are the attine ants and their cultivated fungi. Resembling *Cephalotes* ants in their mode of colony founding, new attine ant colonies are founded by queens – after mating flights – without assistance of non-reproductives. Ensuring the propagation of their externally digestive fungal cultivars, these queens transport a fungal inoculum in a cephalic cavity known as infrabuccal pocket [45]. As hypothesized for *Cephalotes*, this provides a means for vertical symbiont transfer, while necessitating that queens at some point associate with core symbionts. Evidence for occasional host switching between attines and their fungi – and for domestication of novel cultivar lineages – argues that short-term partner fidelity does not always equate to faithful cospeciation [174, 175]. In parallel with this finding, potential cospeciation between *Cephaloticoccus* symbionts and turtle ant hosts is imperfect, with evidence for some host switching (Fig. 8). For the remaining adult-enriched symbionts, inferences on the rates of host-switching, loss, duplication, and cospeciation await more rigorous phylogenetic analysis.

While non-negligible portions of the larval microbiome owe their conservation to partner fidelity, a second mechanism – environmental filtering – has likely driven long-term retention of Enterobacteriales, some Lactobacillales, and Actinobacteria. In particular, we propose that some component of the *Cephalotes* larval gut physiology, or perhaps members of their gut microbiome (*sensu* Scheuring and Yu [176]), could favor the persistence of particular microbes acquired from the environment. Enterobacteriales and Lactobacillales microbes have been found previously in pollen, and in particular pollen that has been collected and stored by hymenopterans [177], providing a plausible source that could explain their presence in turtle ant larval guts. We note, further, that tendencies of the larval gut to support the growth of some adult-enriched bacteria (e.g. Rhizobiales, Alcaligenaceae, Xanthomonadales), while seemingly excluding others (e.g. *Cephaloticoccus* from the Opitutales, Campylobacterales) suggest that filtering mechanisms shape more than just the environmentally acquired inoculate encountered by larvae.

Environmental filtering is thought to play a common role in shaping the variety of bacteria that can survive and thrive in the guts of various insects (e.g. Chandler et al. [19]; Ravenscraft et al. [18]; Kennedy et al. [178]). It has, indeed, promoted long term homogenization in the guts of some such animals, including those of the genus *Drosophila* which retain a conserved range of bacteria from the Enterobacteriaceae, Orbales, Acetobacteriaceae, Lactobacillales. Evidence for this mechanism, and against partner fidelity, begins to accrue when considering there is little phylosymbiosis signal in *Drosophila* [179], including a quartet of mushroom-feeding flies sharing common ancestry ∼28 million years before present [180]. The continued presence of bacteria related to those from the environment, and the acquisition of similar food-borne taxa by distantly related *Drosophila*, further support this idea [19].

Beyond this system, some studies have revealed high specificity of symbionts acquired by insects from the environment, showing also that these hosts preferentially acquire beneficial symbionts, excluding those that are less beneficial. Evidence for this phenomenon, termed partner choice, has been obtained for the beewolf-antennal *Streoptomyces* symbiosis [13]. It is also documented for the bean bug *Riptortus pedestris*, due to evolved anatomical gut features, the corkscrew motility of their preferred symbionts, and the heightened competitive abilities of their beneficial gut symbionts [26]. Whether *Cephalotes* ants have evolved precise means to select beneficial symbionts for growth in the larval gut is not yet established, thought it remains an intriguing prospect.

## Conclusions

In conclusion, this study characterized the gut microbiomes across life stages and castes from 13 species of *Cephalotes* turtle ants. We found that the partner fidelity, which is promoted by transgenerational symbiont transfer through colony-founding queens, plays a central role on the stable and conserved symbiosis between adult ants and gut microbes. In contrast, *Cephalotes* larvae harbor distinct but also conserved microbiomes. The weaker phylosymbiosis patterns and close relatedness to free-living bacteria of larval microbiomes indicates environmental filtering as a primary mechanism behind such larval microbiome conservation. Our findings expand our vision beyond the previous work building mainly upon *Cephalotes* workers, and provide a major change in our understanding of acquisition and persistence of beneficial symbionts across metamorphosis in holometabolous insects.

## Funding

This study was supported by NSF grant 1050360 and 1442144 to J.A.R., NSF grant 1110515 to J.G.S., NSF grant 1442256 to S.P. Funding was also provided by National Natural Science Foundation of China 32070401 to Y. H., Beijing Advanced Innovation Program for Land Surface Science and the “111” Program of Introducing Talents of Discipline to Universities (B13008).

## Notes

### Competing Interest Statement

The authors have declared no competing interest.

