## Supplementary figures for "The secrets to domestic bliss – Partner fidelity and environmental filtering preserve stage-specific turtle ant gut symbioses for over 40 million years"

(1) Supplementary Figure 1: Photos of all ants included in this study.
(2) Supplementary Figure 2: Bacterial composition among gut tissues and gut contents of *Cephalotes varians* larvae.
(3) Supplementary Figure 3: Heatmap illustrating relative abundance of common bacterial 97% OTUs across castes and developmental stages – with an expanded view of bacteria occasionally abundant in larvae.
(4) Supplementary Figure 4: Prevalence and maximum relative abundance for common OTUs across larvae vs. adult workers.
(5) Supplementary Figure 5: Heatmap illustrating oligotype composition across individual *Cephalotes* ants.
(6) Supplementary Figure 6: Maximum likelihood phylogenetic analysis of the Actinobacteria (Analysis #4).
(7) Supplementary Figure 7: Maximum likelihood phylogenetic analysis of the Lactobacillales (Analysis #5).
(8) Supplementary Figure 8: Maximum likelihood phylogenetic analysis of ambiguously classifying Pseudomonadales OTU046 (Analysis #6).
(9) Supplementary Figure 9: Maximum likelihood phylogenetic analysis of ambiguously classifying Sphingobacteriales OTU116 (Analysis #7).
(10) Supplementary Figure 10: Maximum likelihood phylogenetic analysis of other sequences with high abundance, and their relatives (Analysis #8).
(11) Supplementary Figure 11: Proportions of microbiomes comprised by bacteria from cephalotine-specialist lineages.
(12) Supplementary Figure 12: Bacterial abundance across larvae of varying age.
(13) Supplementary Figure 13: Cophylogeny analyses for *Cephaloticoccus* symbionts and *Cephalotes* ant hosts.

**Supplementary Figure 1: Photos of all ants included in this study.**

*Cephalotes* aff. *bimaculatus*: colony JS-C-220

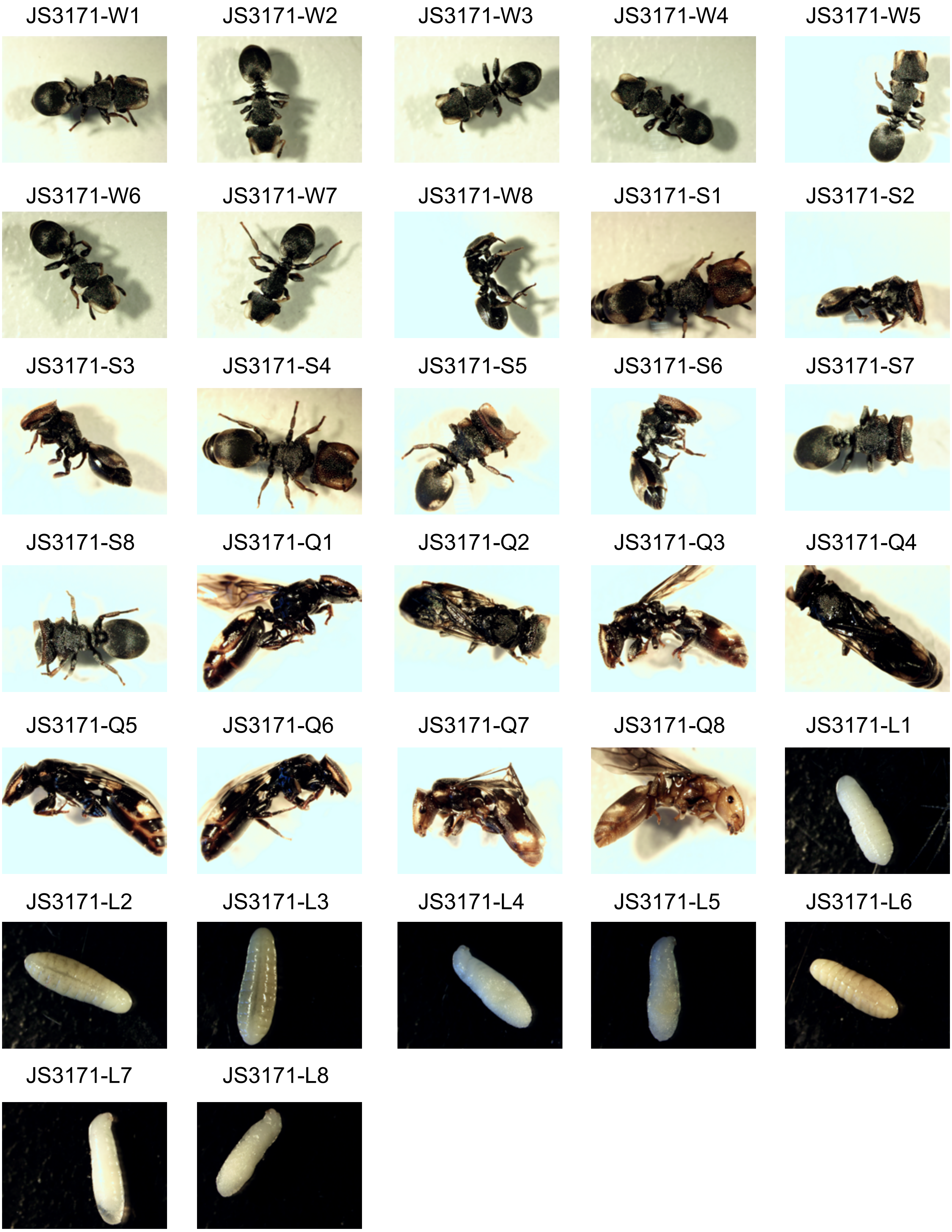

*Cephalotes* aff. *wheeleri*: colony JS-C-211

JS3092-W1

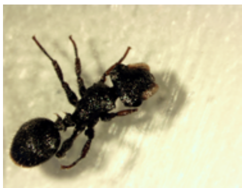

JS3092-W2

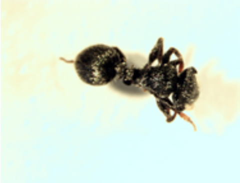

JS3092-W3

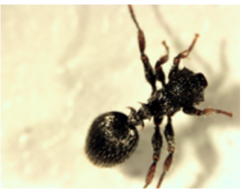

JS3092-W4

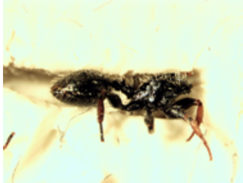

JS3092-W5

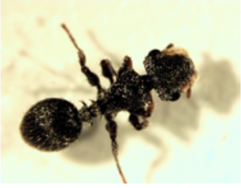

JS3092-W6

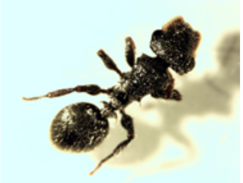

JS3092-W7

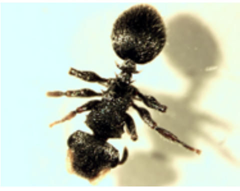

JS3092-W8

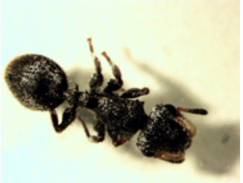

JS3093-S1

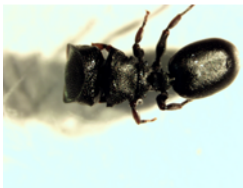

JS3093-S2

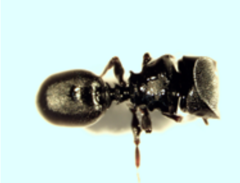

JS3093-S3

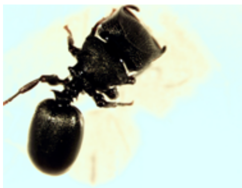

JS3093-S4

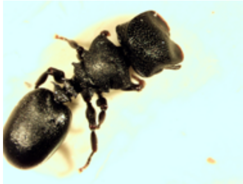

JS3093-S5

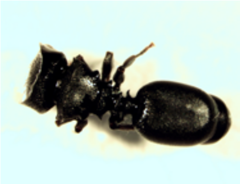

JS3093-S6

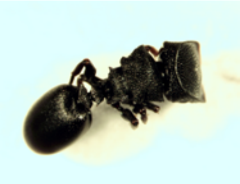

JS3093-S7

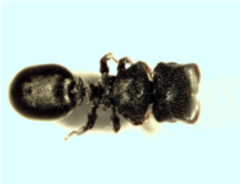

JS3093-S8

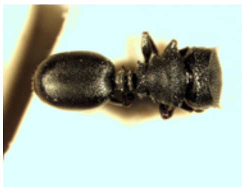

JS3094-L1

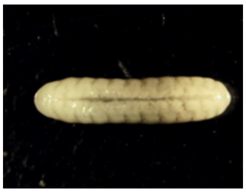

JS3094-L2

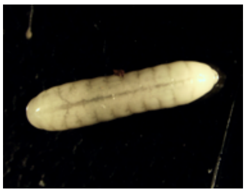

JS3094-L3

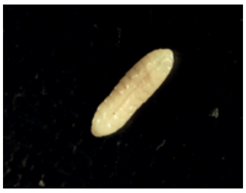

JS3094-L4

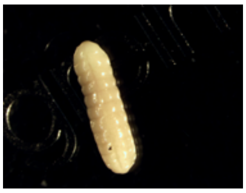

JS3094-L5

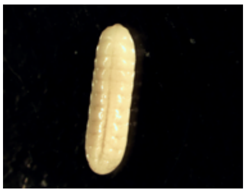

JS3094-L6

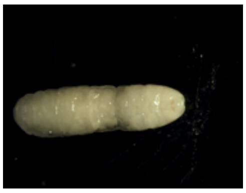

JS3094-L7

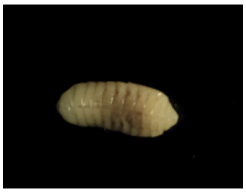

JS3094-L8

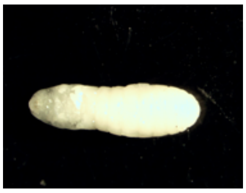

*Cephalotes depressus*: colony C15-25

C15-25-W1

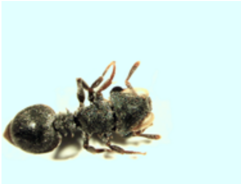

C15-25-W2

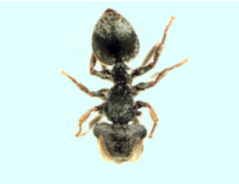

C15-25-W3

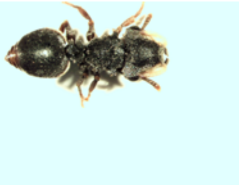

C15-25-W4

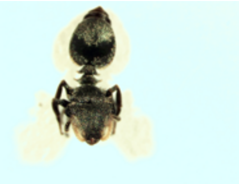

C15-25-S1

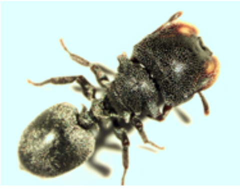

C15-25-S2

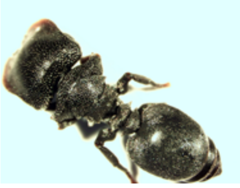

C15-25-S3

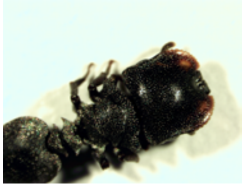

C15-25-S4

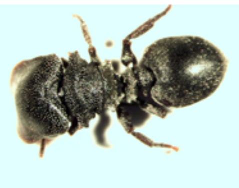

C15-25-C1

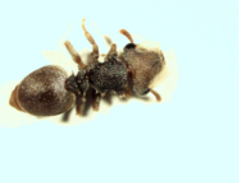

C15-25-C2

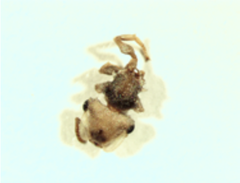

C15-25-Q1

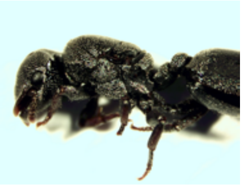

C15-25-L1

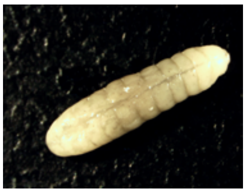

C15-25-L2

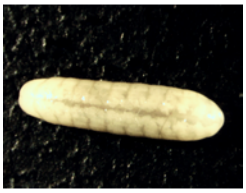

C15-25-L3

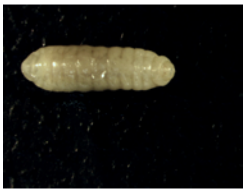

C15-25-L4

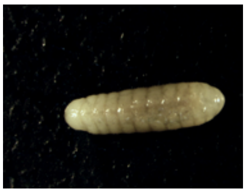

C15-25-L5

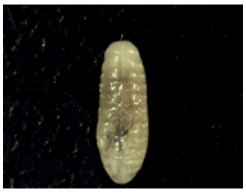

C15-25-L6

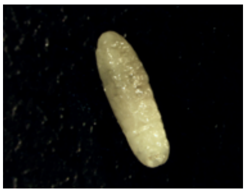

C15-25-L7

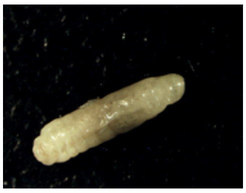

C15-25-L8

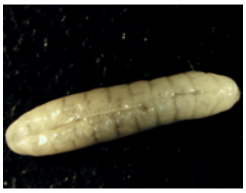

*Cephalotes goniodontus*: colony JS-3-203

JS3026-W1

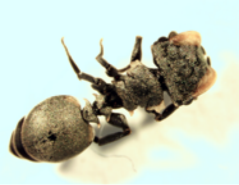

JS3026-W2

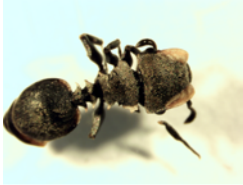

JS3026-W3

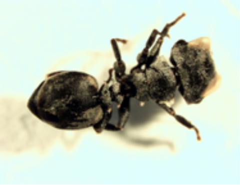

JS3026-W4

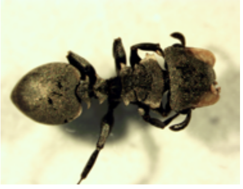

JS3026-C1

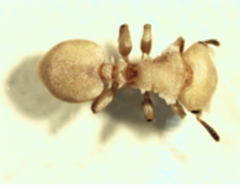

JS3026-C2

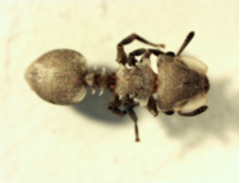

JS3026-C3

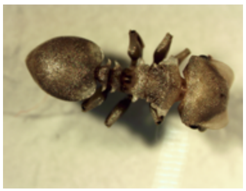

JS3024-Q1

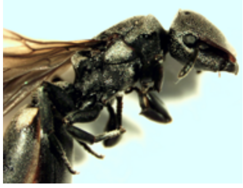

JS3024-Q2

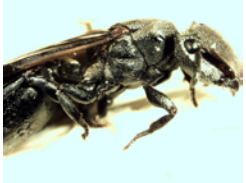

JS3024-Q3

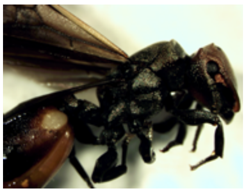

JS3024-Q4

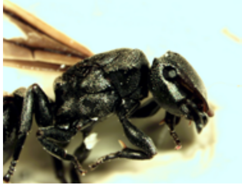

JS3024-M1

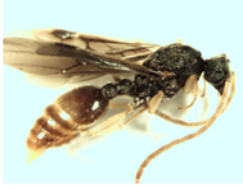

JS3024-M2

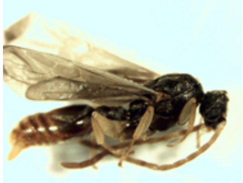

JS3024-M3

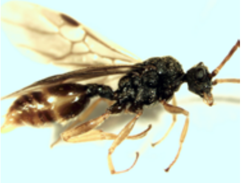

JS3023-L1

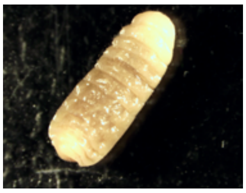

JS3023-L2

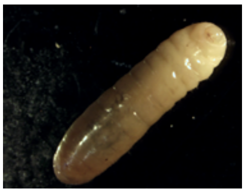

JS3023-L3

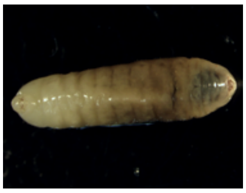

JS3023-L4

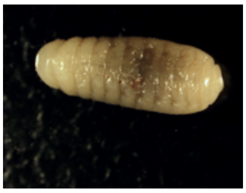

JS3023-L5

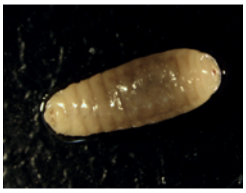

JS3023-L6

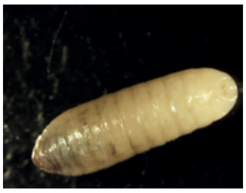

JS3023-L7

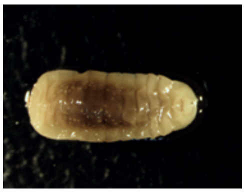

JS3023-L8

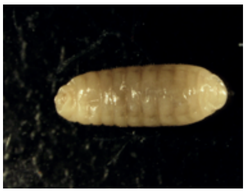

*Cephalotes grandinosus*: colony C15-21

C15-21-W1
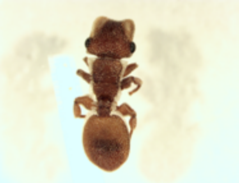

C15-21-W2
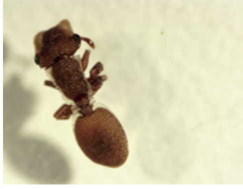

C15-21-W3
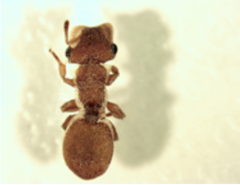

C15-21-W4
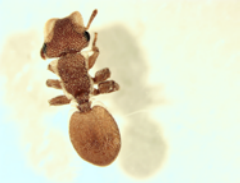

C15-21-W5
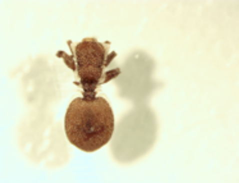

C15-21-S1
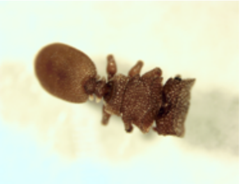

C15-21-L1
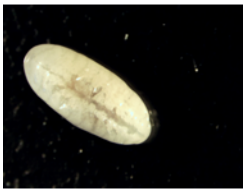

C15-21-L2
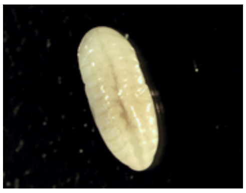

C15-21-L4
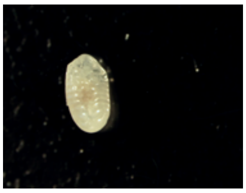

C15-21-L5
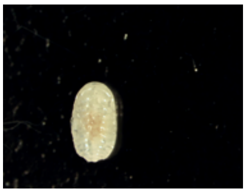

*Cephalotes grandinosus*: colony C15-44

C15-44-W1
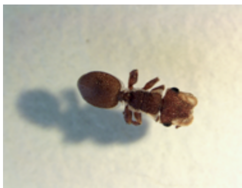

C15-44-W2
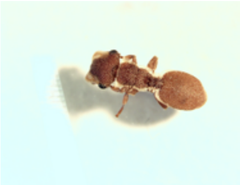

C15-44-W3
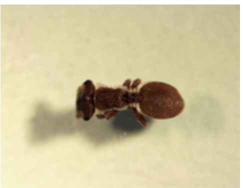

C15-44-W4
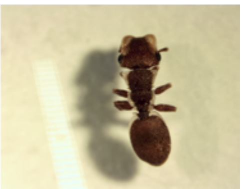

C15-44-C1
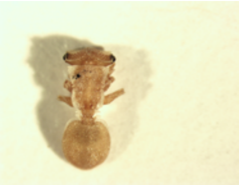

C15-44-C2
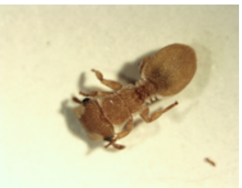

C15-44-S1
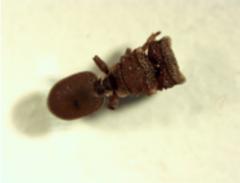

C15-44-S2
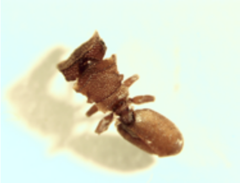

C15-44-S3
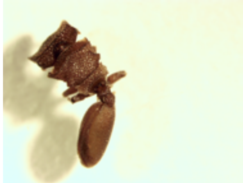

C15-44-S4
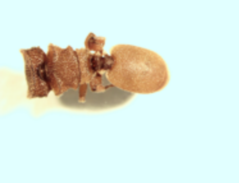

C15-44-Q1
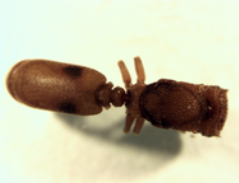

C15-44-L1
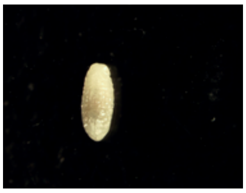

C15-44-L2
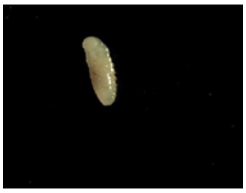

C15-44-L3
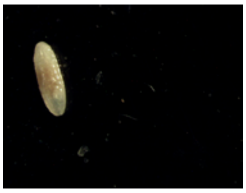

C15-44-L4
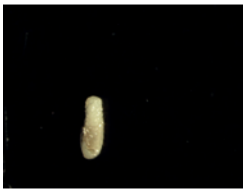

C15-44-L5
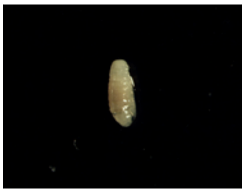

C15-44-L6
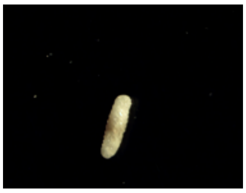

*Cephalotes maculatus*: colony C15-28

C15-28-W1

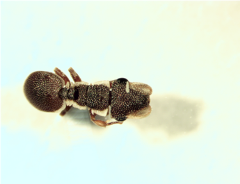

C15-28-W2

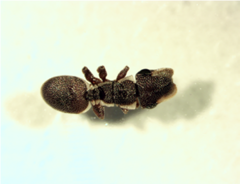

C15-28-W3

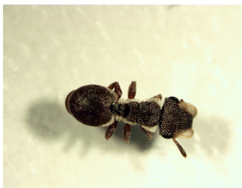

C15-28-W4

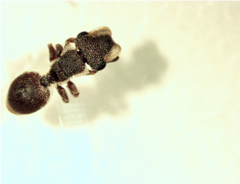

C15-28-S1

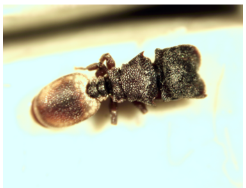

C15-28-S2

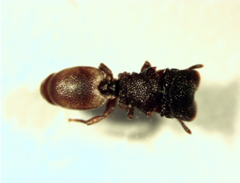

C15-28-S3

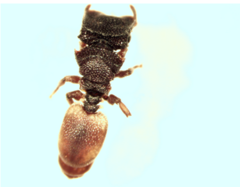

C15-28-S4

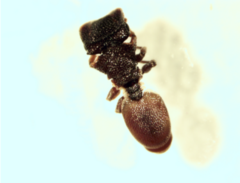

C15-28-C1

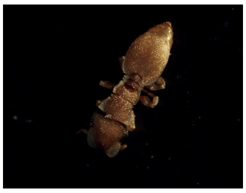

C15-28-C2

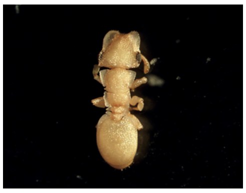

C15-28-Q1

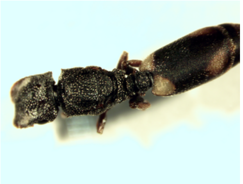

C15-28-L1

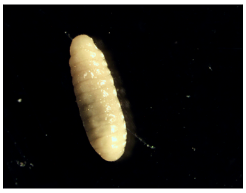

C15-28-L2

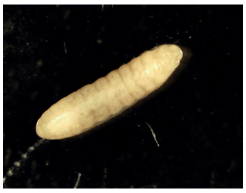

C15-28-L3

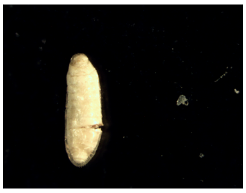

C15-28-L4

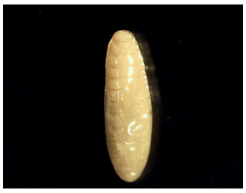

C15-28-L5

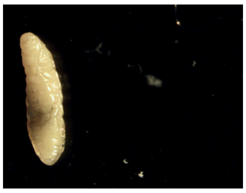

C15-28-L6

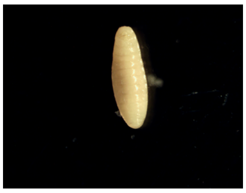

C15-28-L7

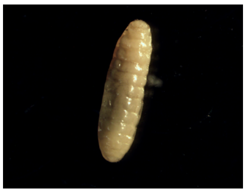

C15-28-L8

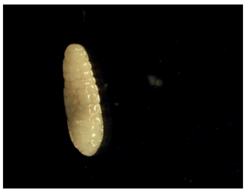

*Cephalotes minutus*: colony SRCmin1

SRCmin1-A1

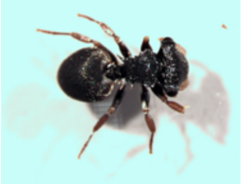

SRCmin1-A2

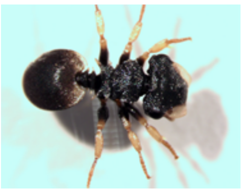

SRCmin1-A3

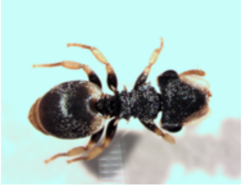

SRCmin1-A4

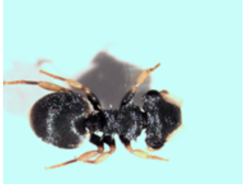

SRCmin1-A5

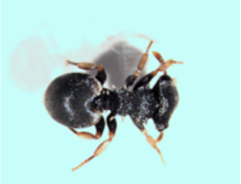

SRCmin1-A6

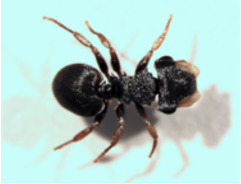

SRCmin1-A7

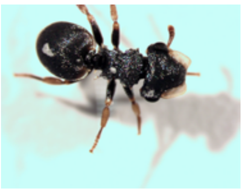

SRCmin1-A8

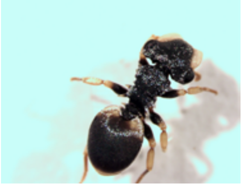

SRCmin1-C1

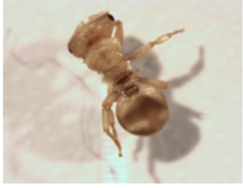

SRCmin1-C2

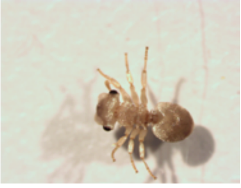

SRCmin1-C3

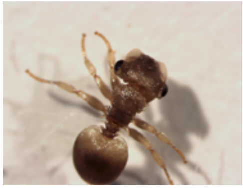

SRCmin1-C4

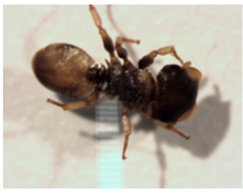

SRCmin1-S1

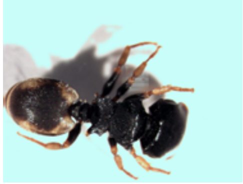

SRCmin1-S2

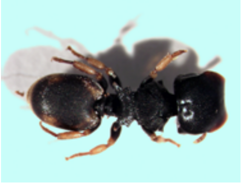

SRCmin1-S3

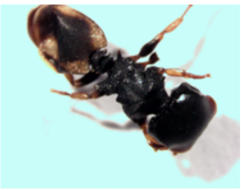

SRCmin1-L1

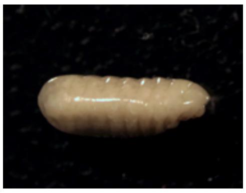

SRCmin1-L2

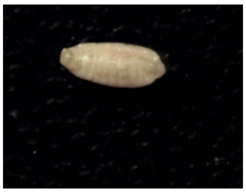

SRCmin1-L3

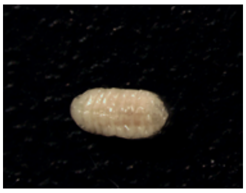

SRCmin1-L4

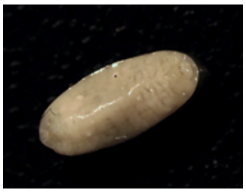

SRCmin1-L5

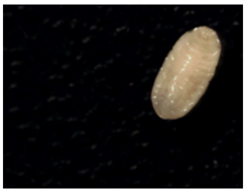

SRCmin1-L6

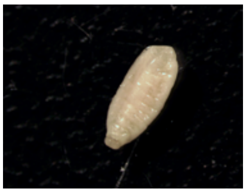

SRCmin1-L7

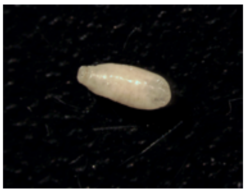

SRCmin1-L8

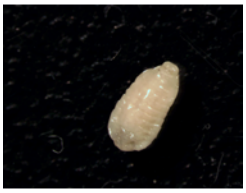

SRCmin1-PP1

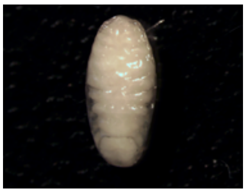

SRCmin1-PP2

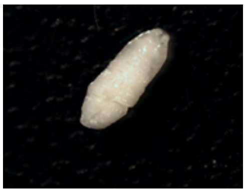

SRCmin1-PP3

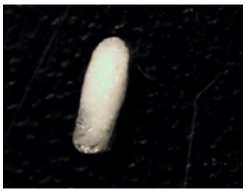

SRCmin1-PP4

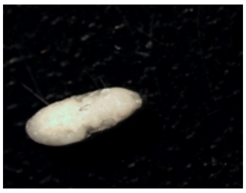

SRCmin1-PP5

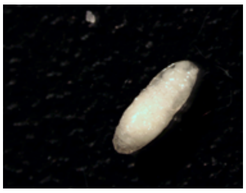

SRCmin1-P1

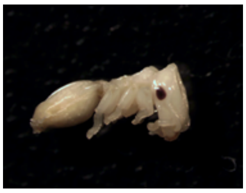

SRCmin1-P2

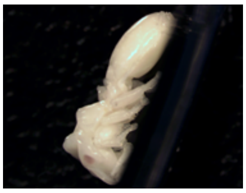

SRCmin1-P3

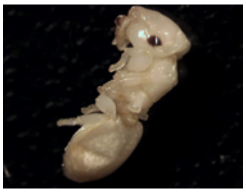

SRCmin1-P4

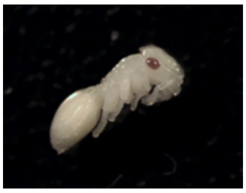

SRCmin1-P5

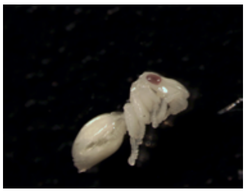

SRCmin1-P6

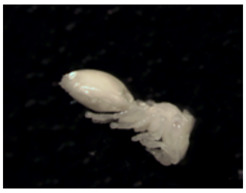

SRCmin1-P7

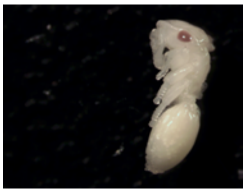

SRCmin1-P8

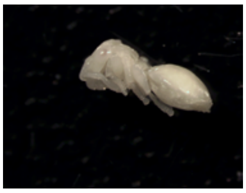

*Cephalotes pusillus*: colony C15-40

C15-40-W1

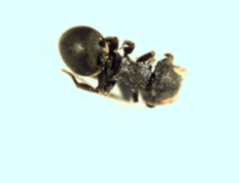

C15-40-W2

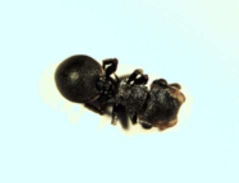

C15-40-W3

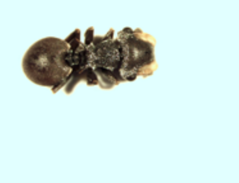

C15-40-W4

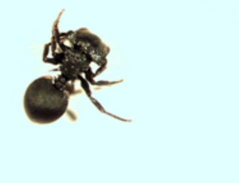

C15-40-S1

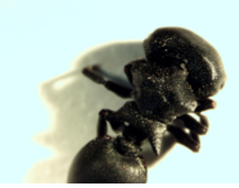

C15-40-S2

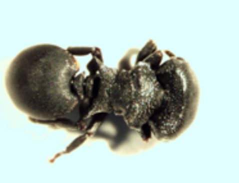

C15-40-S3

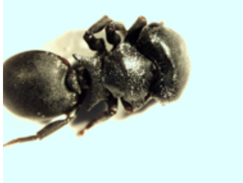

C15-40-S4

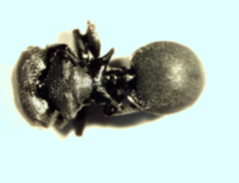

C15-40-Q1

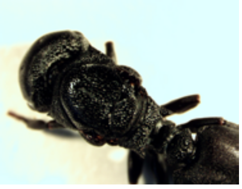

C15-40-L1

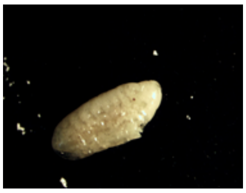

C15-40-L2

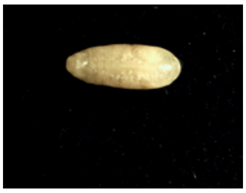

C15-40-L3

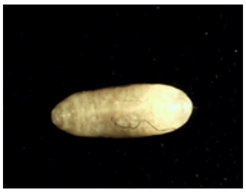

C15-40-L4

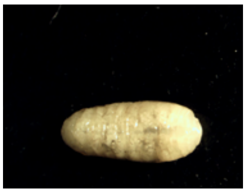

C15-40-L5

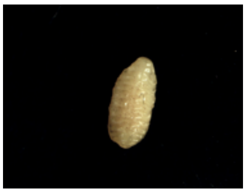

C15-40-L6

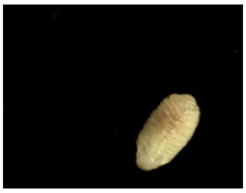

C15-40-L7

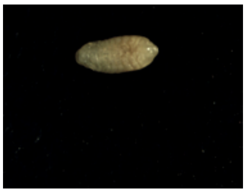

C15-40-L8

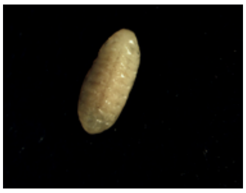

*Cephalotes setulifer*: colony SRCset1

SRCset1-A1

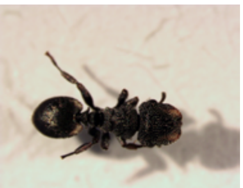

SRCset1-A2

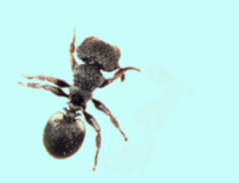

SRCset1-A3

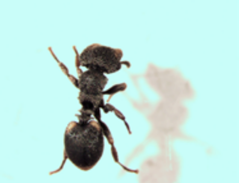

SRCset1-A4

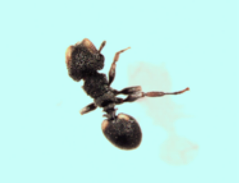

SRCset1-A5

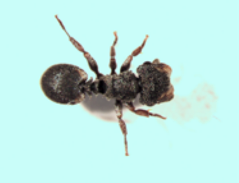

SRCset1-A6

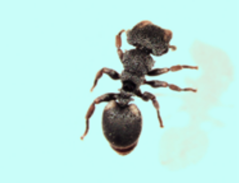

SRCset1-A7

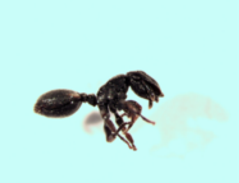

SRCset1-A8

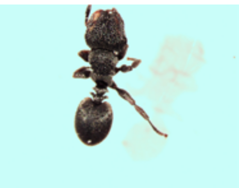

SRCset1-S1

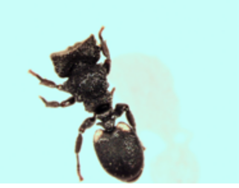

SRCset1-S2

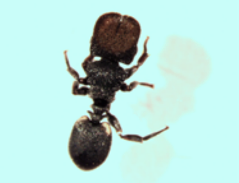

SRCset1-S3

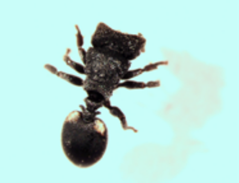

SRCset1-Q1

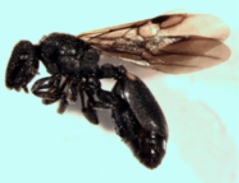

SRCset1-L1

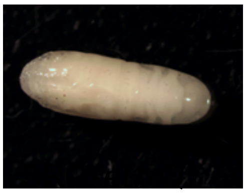

SRCset1-L2

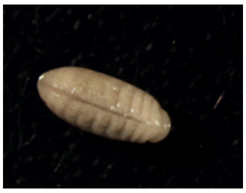

SRCset1-L3

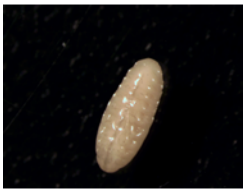

SRCset1-L4

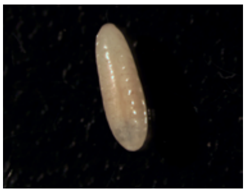

SRCset1-L5

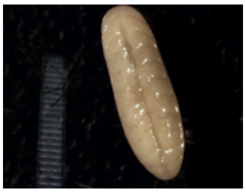

SRCset1-P1

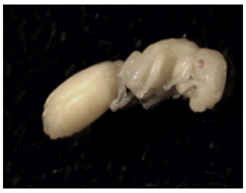

SRCset1-P2

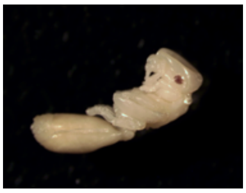

SRCset1-P3

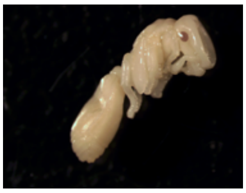

SRCset1-P4

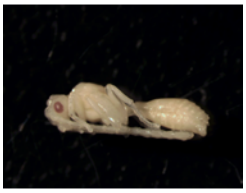

SRCset1-P5

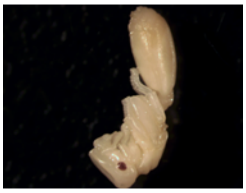

*Cephalotes setulifer*: colony SRCset2

SRCset2-A1
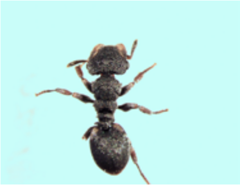
SRCset2-A2
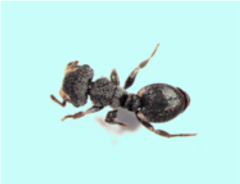
SRCset2-A3
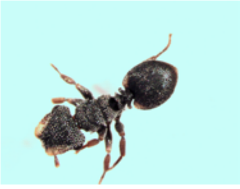
SRCset2-A4
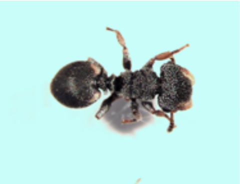
SRCset2-A5
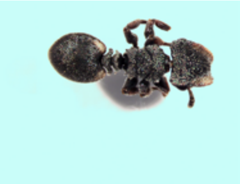

SRCset2-A6
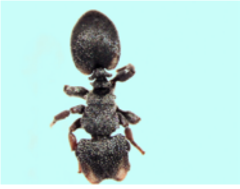
SRCset2-A7
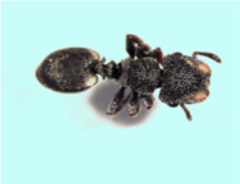
SRCset2-A8
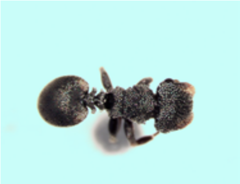
SRCset2-S1
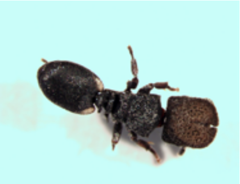
SRCset2-S2
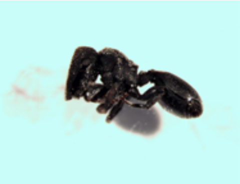

SRCset2-Q1
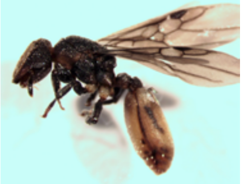
SRCset2-Q2
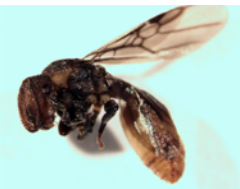
SRCset2-Q3
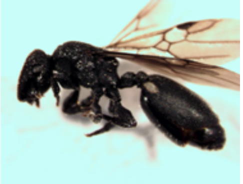
SRCset2-Q4
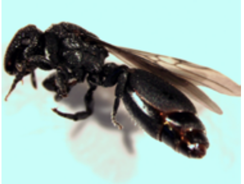
SRCset2-L1
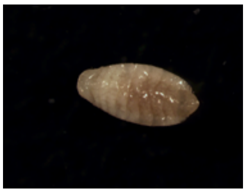

SRCset2-L2
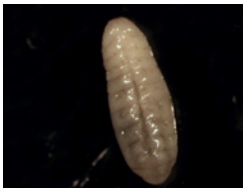
SRCset2-L3
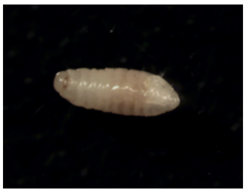
SRCset2-L4
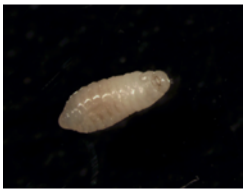
SRCset2-L5
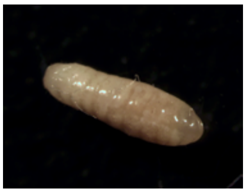
SRCset2-L6
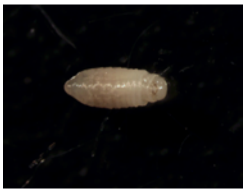

SRCset2-L7
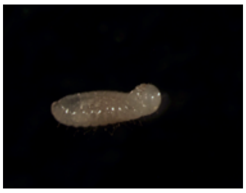
SRCset2-L8
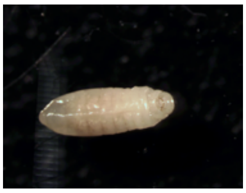
SRCset2-P1
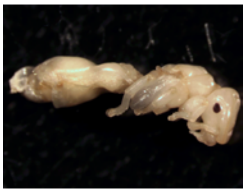
SRCset2-P2
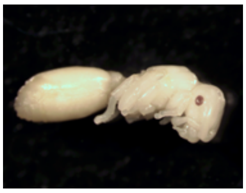
SRCset2-P3
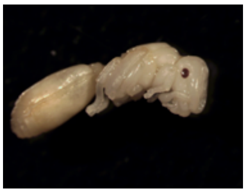

*Cephalotes targionii*: colony C15-20

C15-20-W1

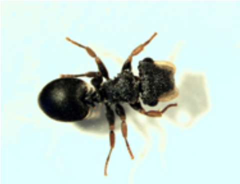

C15-20-W2

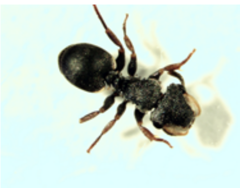

C15-20-W3

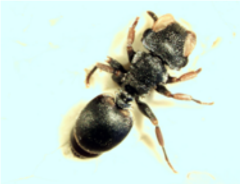

C15-20-W4

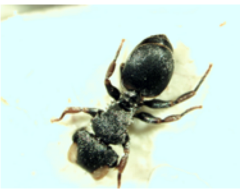

C15-20-C1

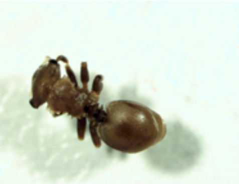

C15-20-S1

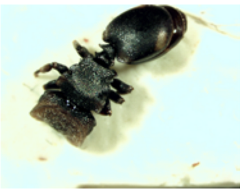

C15-20-S2

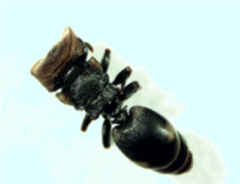

C15-20-S3

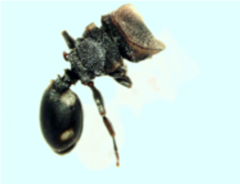

C15-20-S4

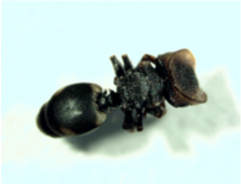

C15-20-L1

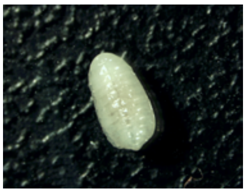

C15-20-L2

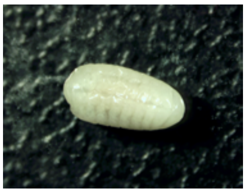

C15-20-L3

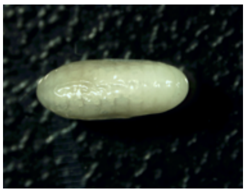

C15-20-L4

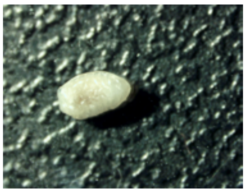

C15-20-L5

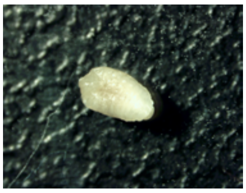

C15-20-L6

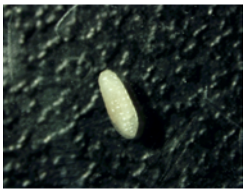

*Cephalotes texanus*: colony JS-C-235

JS3260-W1

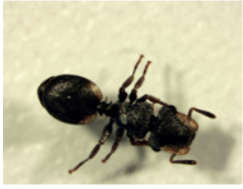

JS3260-W2

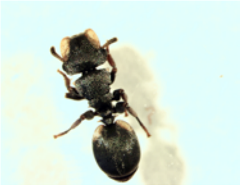

JS3260-W3

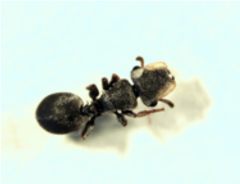

JS3260-W4

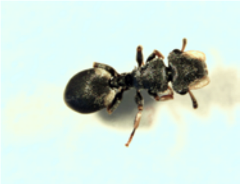

JS3261-S1

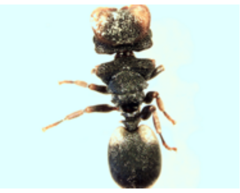

JS3261-S2

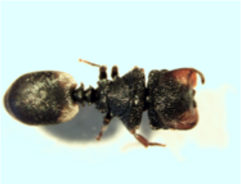

JS3261-S3

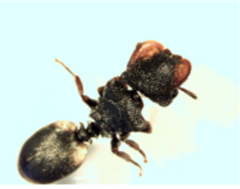

JS3261-S4

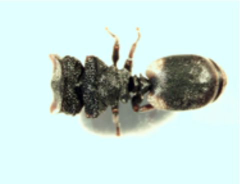

JS3262-L1

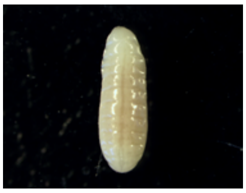

JS3262-L2

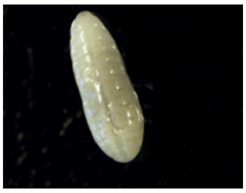

JS3262-L3

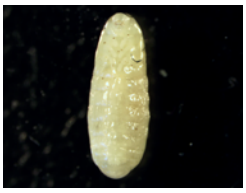

JS3262-L4

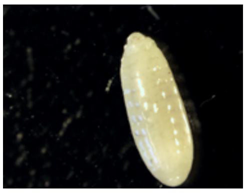

*Cephalotes unimaculatus*: colony C16-08

C16-08-W1

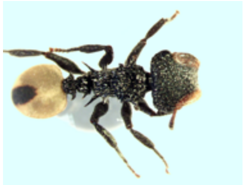

C16-08-W2

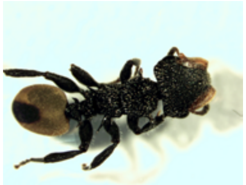

C16-08-W3

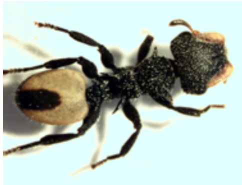

C16-08-W4

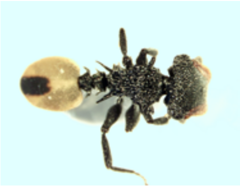

C16-08-C1

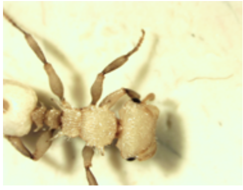

C16-08-C2

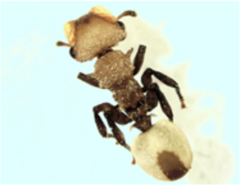

C16-08-L1

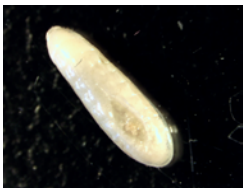

C16-08-L2

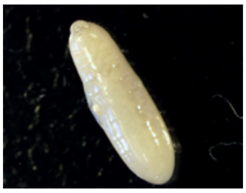

C16-08-L3

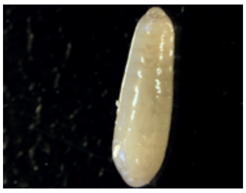

C16-08-L4

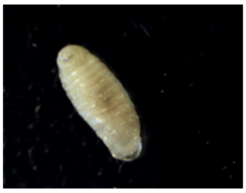

C16-08-L5

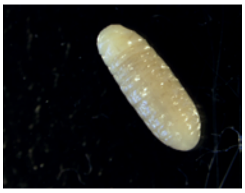

C16-08-L6

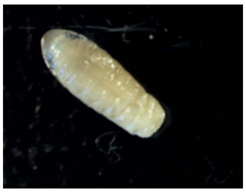

C16-08-L7

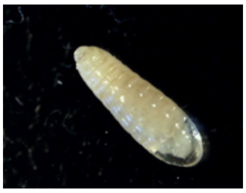

C16-08-L8

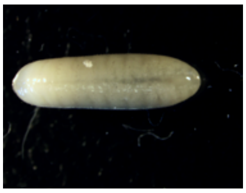

#### *Cephalotes varians*: colony YH083

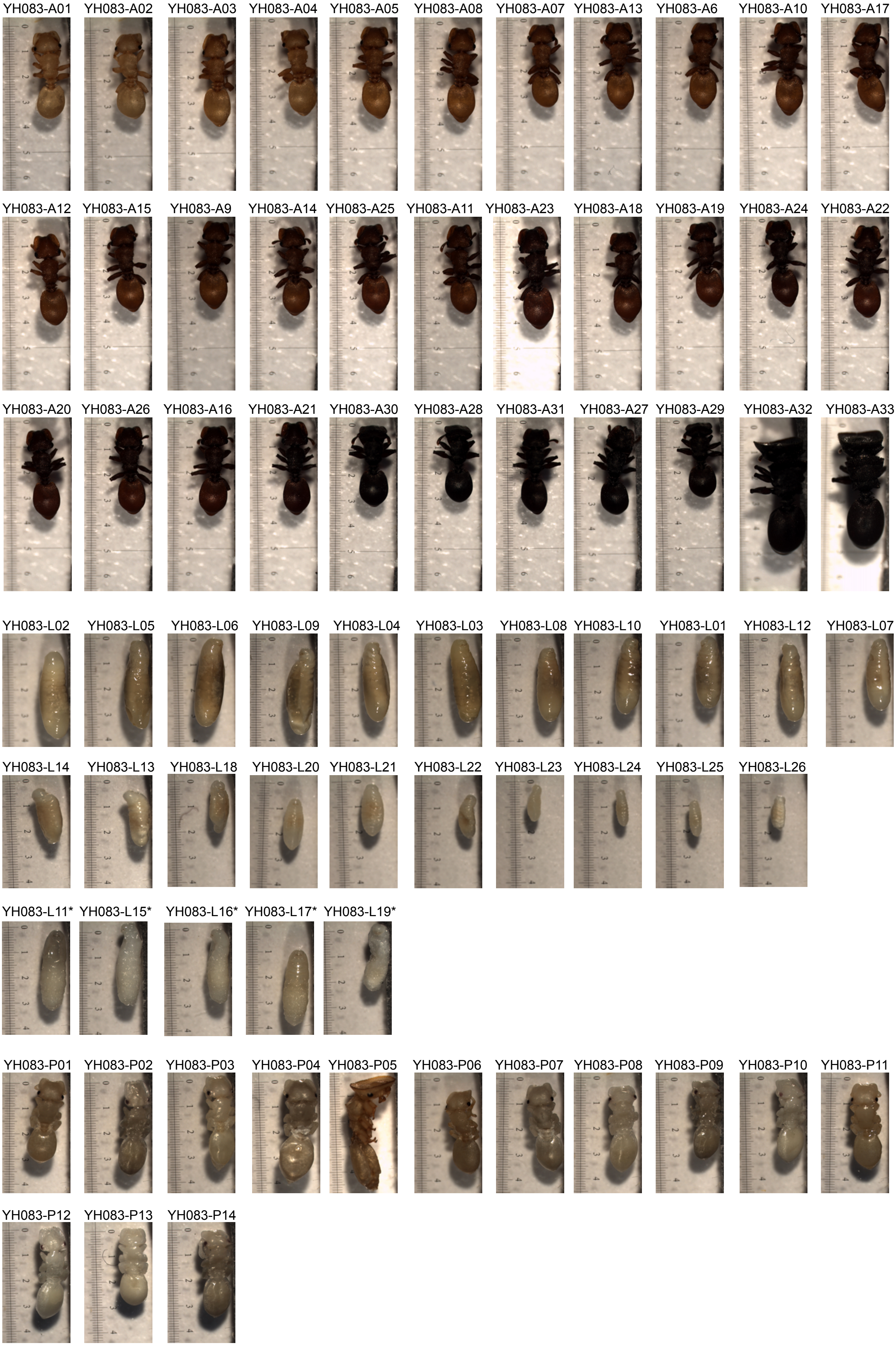

### *Cephalotes varians*: colony YH091

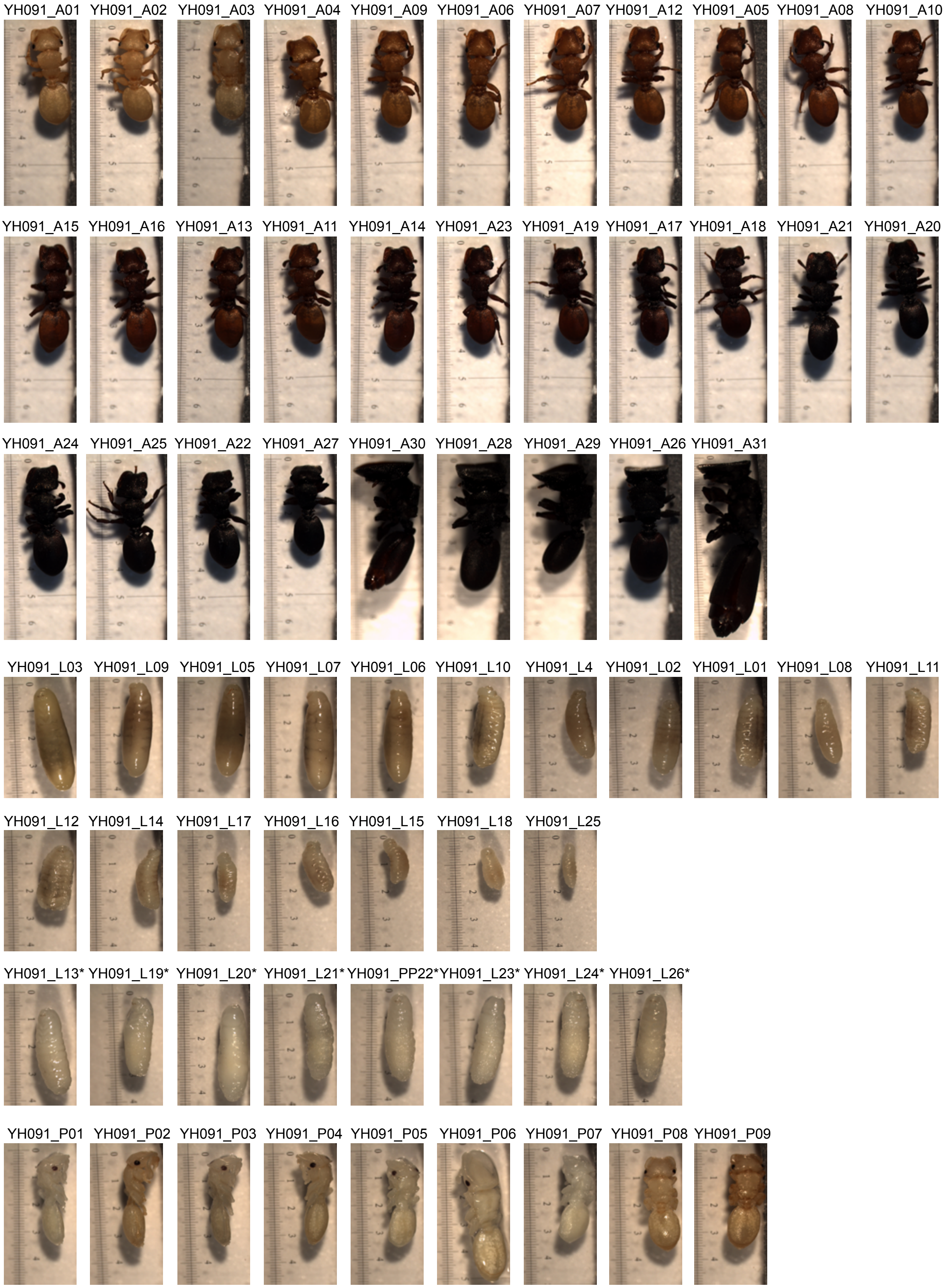

*Cephalotes varians*: colony YH092

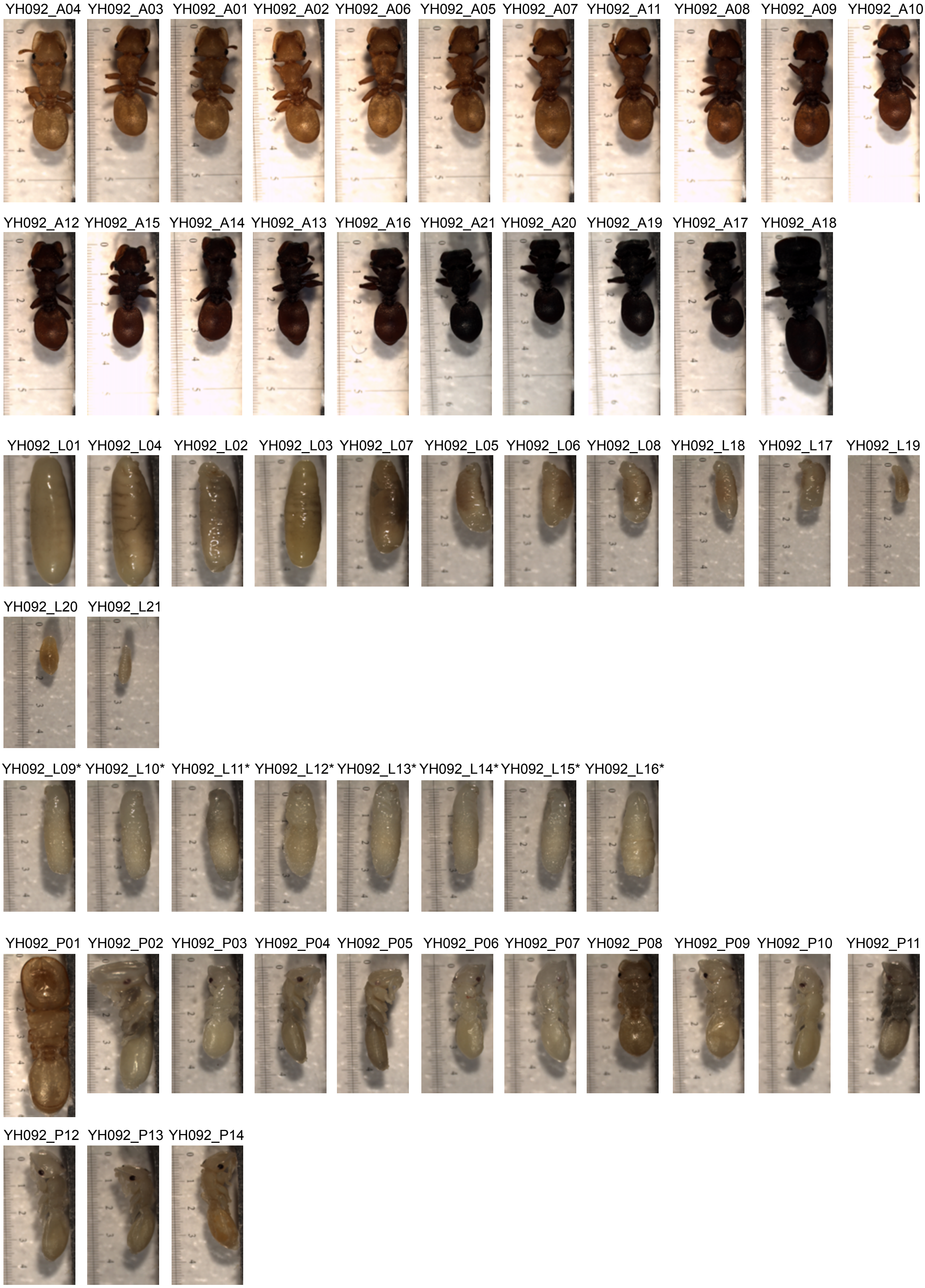

*Cephalotes varians*: colony YH100

YH100_A03 YH100_A01 YH100_A02 YH100_A04 YH100_A05 YH100_A07 YH100_A06 YH100_A08 YH100_A09 YH100_A12 YH100_A11

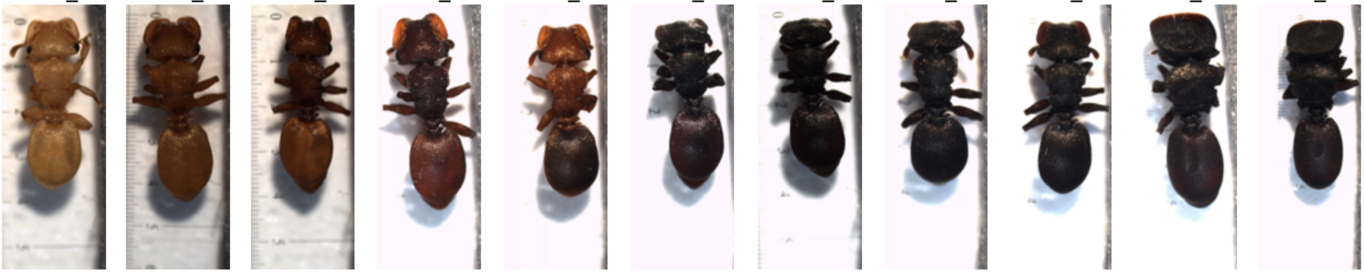

YH100_A10 YH100_A13

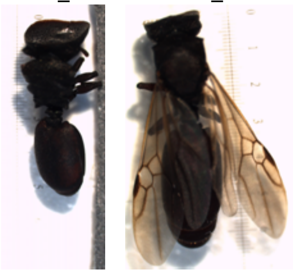

YH100_L01 YH100_L02 YH100_L03 YH100_L05 YH100_L07 YH100_L06 YH100_L08 YH100_L09 YH100_L10 YH100_L11 YH100_L12

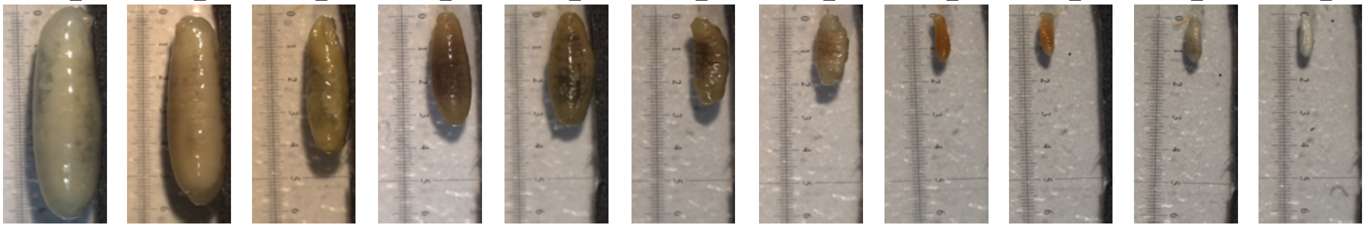

YH100_L13 YH100_L14

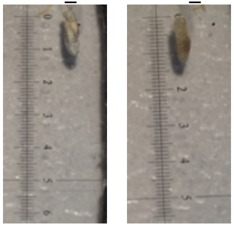

YH100_L04*

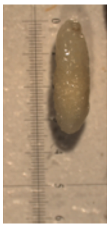

YH100_P01 YH100_P02 YH100_P03 YH100_P04 YH100_P05

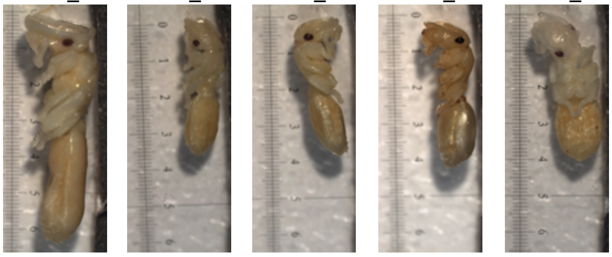

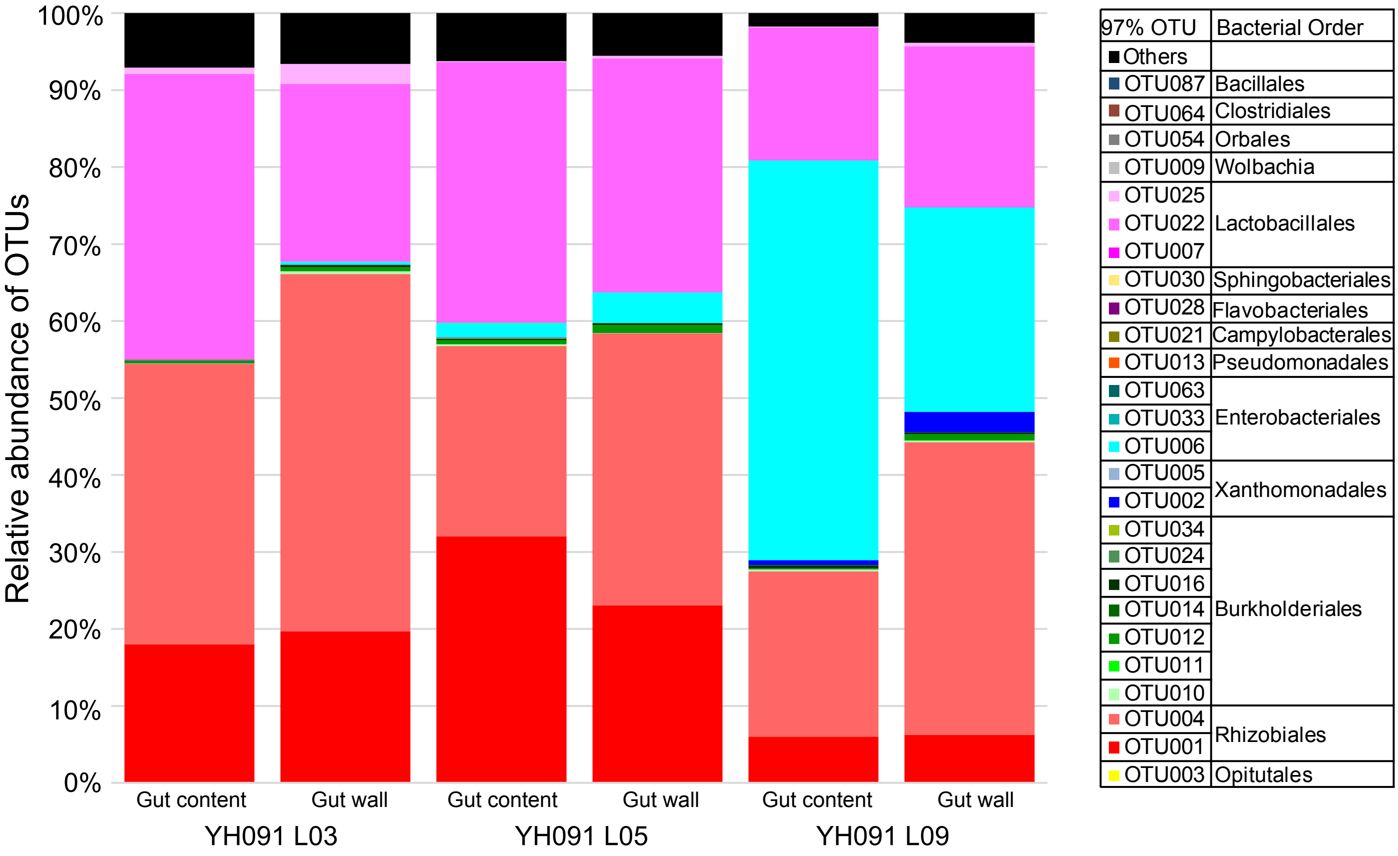


**Supplementary Figure 2: Bacterial composition among gut tissues and gut contents of** ***Cephalotes varians*** **larvae.** Stacked bar graphs illustrate bacterial composition at the 97% OTU level.

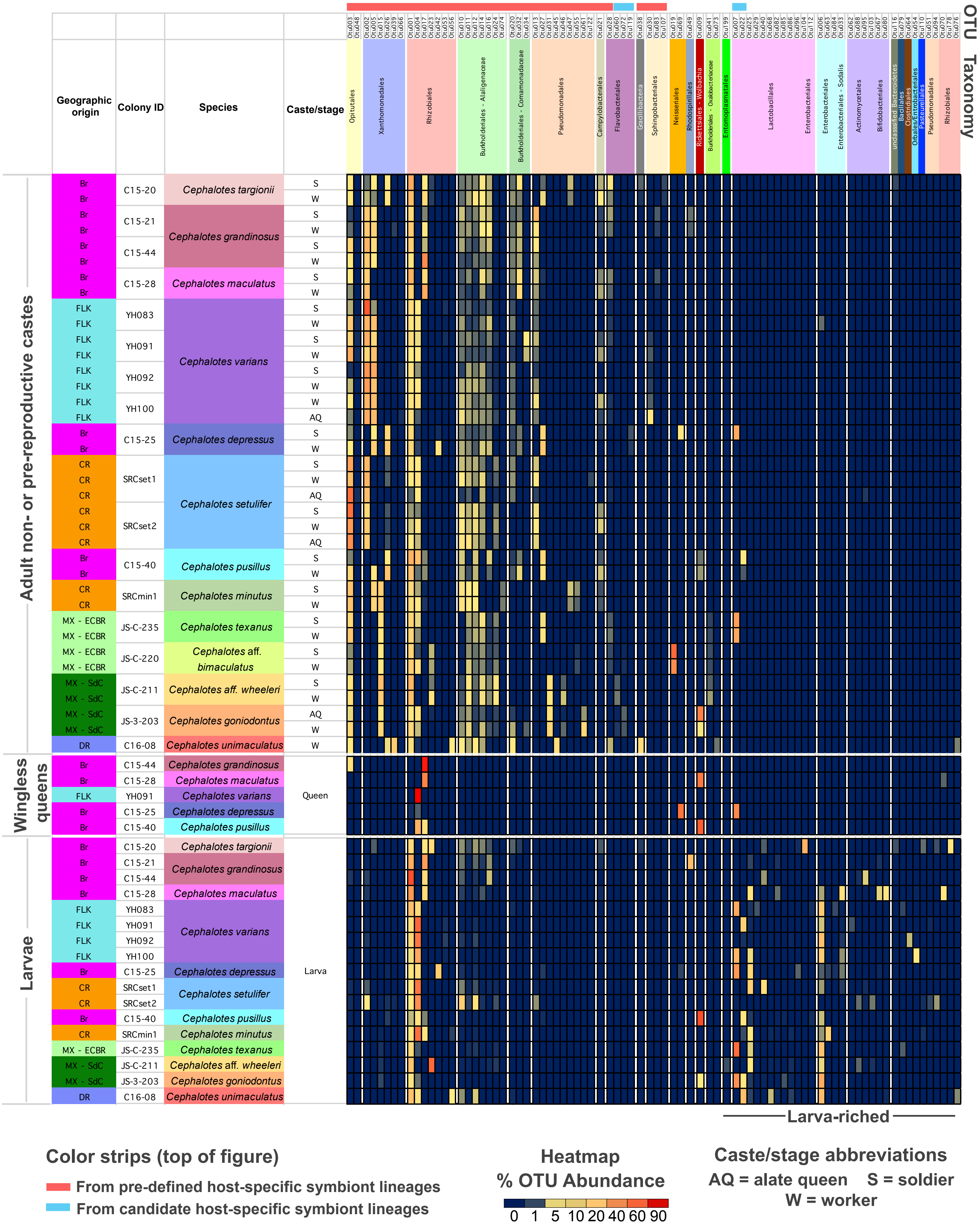

OTU
Taxonomy
Geographic origin
Colony ID
Species
Caste/stage
Adult non- or pre-reproductive castes
Wingless queens
Larvae
Cephalotes targionii
Cephalotes grandinosus
Cephalotes maculatus
Cephalotes varians
Cephalotes depressus
Cephalotes setulifer
Cephalotes pusillus
Cephalotes minutus
Cephalotes texanus
Cephalotes aff. bimaculatus
Cephalotes aff. wheeleri
Cephalotes goniodontus
Cephalotes unimaculatus
Queen
Larva
Opitutales
Xanthomonadales
Rhizobiales
Burkholderiales - Alaligenaceae
Burkholderiales - Comamonadaceae
Pseudomonadales
Campylobacterales
Flavobacteriales
Gracilibacteria
Sphingobacteriales
Neisseriales
Rhodospirillales
Rickettsiales - Wolbachia
Entomoplasmatales
Lactobacillales
Enterobacteriales
Enterobacteriales - Sodalis
Actinomycetales
Bifidobacteriales
Larva-riched
Color strips (top of figure)
From pre-defined host-specific symbiont lineages
From candidate host-specific symbiont lineages
Heatmap
% OTU Abundance
0 1 5 10 20 40 60 90
Caste/stage abbreviations
AQ = alate queen S = soldier
W = worker

**Supplementary Figure 3: Relative abundance of common bacterial 97% OTUs across castes and developmental stages – with an expanded view of bacteria at least occasionally abundant in larvae.** This heatmap figure builds upon patterns shown in Figure 3, but instead illustrates averages for each developmental stage or caste (worker, soldier, alate queen, wingless queen, or larva) within each studied colony, as opposed to the individual sequence library values shown in Fig. 3. Shown here are a subset of the 154 OTUs that exceeded 0.002994 relative abundance in at least two sequence libraries, or at least 0.05 in one. Of the 99 such OTUs illustrated here, all either exhibited a pooled average of at least 0.01 across the individuals of at least one caste or stage in a given colony, or exceeded 0.002994 average relative abundance in a caste or stage across 2 or more colonies. Figure illustrates how: (1) Non-reproductive adult microbiomes are dominated by OTUs from apparently *Cephalotes*-specific bacterial lineages (red & blue color strips atop figure). (2) Mature, wingless queens have low diversity microbiomes often dominated by Rhizobiales. (3) Larvae are enriched for Rhizobiales, Enterobacteriales, and Lactobacillales. Bacteria from the latter two orders are rarely found in adults. (4) Larvae commonly harbor additional bacteria from outside specialized *Cephalotes*-associated lineages, including Actinobacteria. (5) These immatures can also harbor high titers of bacteria from a subset of adult-enriched, specialized *Cephalotes* symbionts, including Burkholderiales from the Alcaligenaceae, but rarely Opitutales, Campylobacterales, Flavobacteriales, and Sphingobacteriales, for example. Colors used for geography and for both host & bacterial taxonomy match those used in Figure 3.

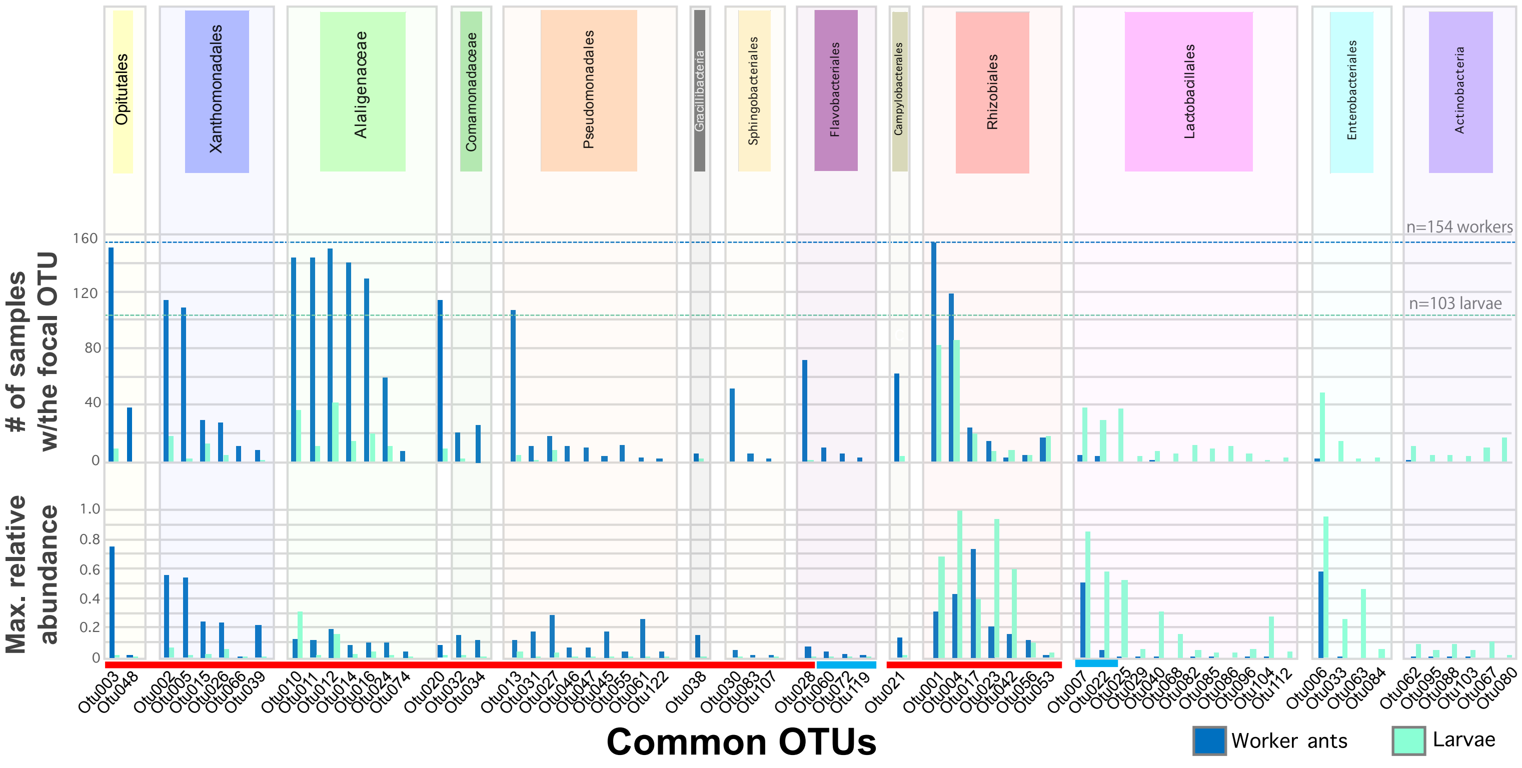


**Supplementary Figure 4: Comparing the prevalence and maximum relative abundance for common OTUs across larvae vs. adult workers.** Data derived from amplicon sequencing of 16S rRNA from 103 larvae (n=12 *Cephalotes* species) and 154 adult workers (n=13 *Cephalotes* species). Bacteria from previously recognized cephalotine-specialist lineages are emphasized with a red line above their OTU names, while those newly recognized here are emphasized with blue. The remainder showed close relatedness to free-living bacteria.

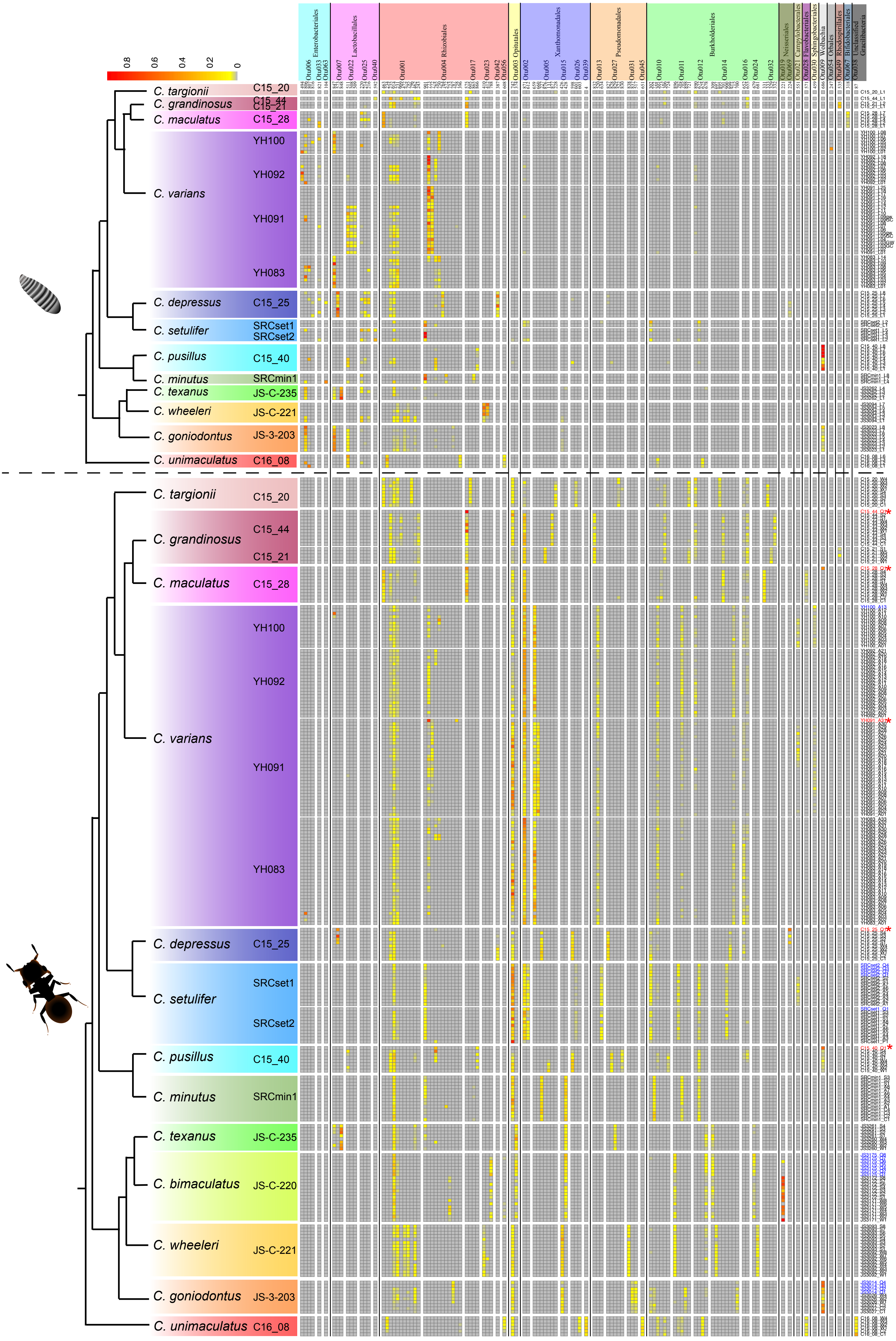

0.8
0.6
0.4
0.2
0
Enterobacteriales
Lactobacillales
Rhizobiales
Opitutales
Xanthomonadales
Pseudomonadales
Burkholderiales
Neisseriales
Campylobacterales
Flavobacteriales
Sphingobacteriales
Wolbachia
Orbales
Rhodospirillales
Bifidobacteriales
Unclassified Gracilibacteria
Otu006 Otu033 Otu063
Otu007 Otu022 Otu025 Otu040
Otu001 Otu004 Otu017 Otu023 Otu042 Otu056
Otu003
Otu002 Otu005 Otu015 Otu026 Otu039
Otu013 Otu027 Otu031 Otu045
Otu010 Otu011 Otu012 Otu014 Otu016 Otu024 Otu032
Otu019 Otu069 Otu021 Otu028 Otu030 Otu009 Otu054 Otu049 Otu067 Otu038
C. targionii C15_20
C. grandinosus C15_44 C15_21
C. maculatus C15_28
C. varians YH100 YH092 YH091 YH083
C. depressus C15_25
C. setulifer SRCset1 SRCset2
C. pusillus C15_40
C. minutus SRCmin1
C. texanus JS-C-235
C. bimaculatus JS-C-220
C. wheeleri JS-C-221
C. goniodontus JS-3-203
C. unimaculatus C16_08

**Figure S5: Heatmap illustrating oligotype composition across individual *Cephalotes* ants.** Shown are data from all 336 focal sequence libraries, plus those from callow ants (n=16 samples with "_C#" in Sample ID column) and a pupa (n=1 sample with "_P#" under "Sample ID column). Libraries from alate queens (blue font) and wingless queens (red font w/asterisks) are highlighted in the Sample ID column. Upper panel illustrates data from larvae, while lower panel shows results from all other stages. Columns show each of the 40 focal OTUs (39 used for enterotyping analysis, plus *Wolbachia*), and their constituent oligotypes. Rows show individual libraries. Relative abundances are shown with a gray-yellow-red heatmap as illustrated in the upper left.

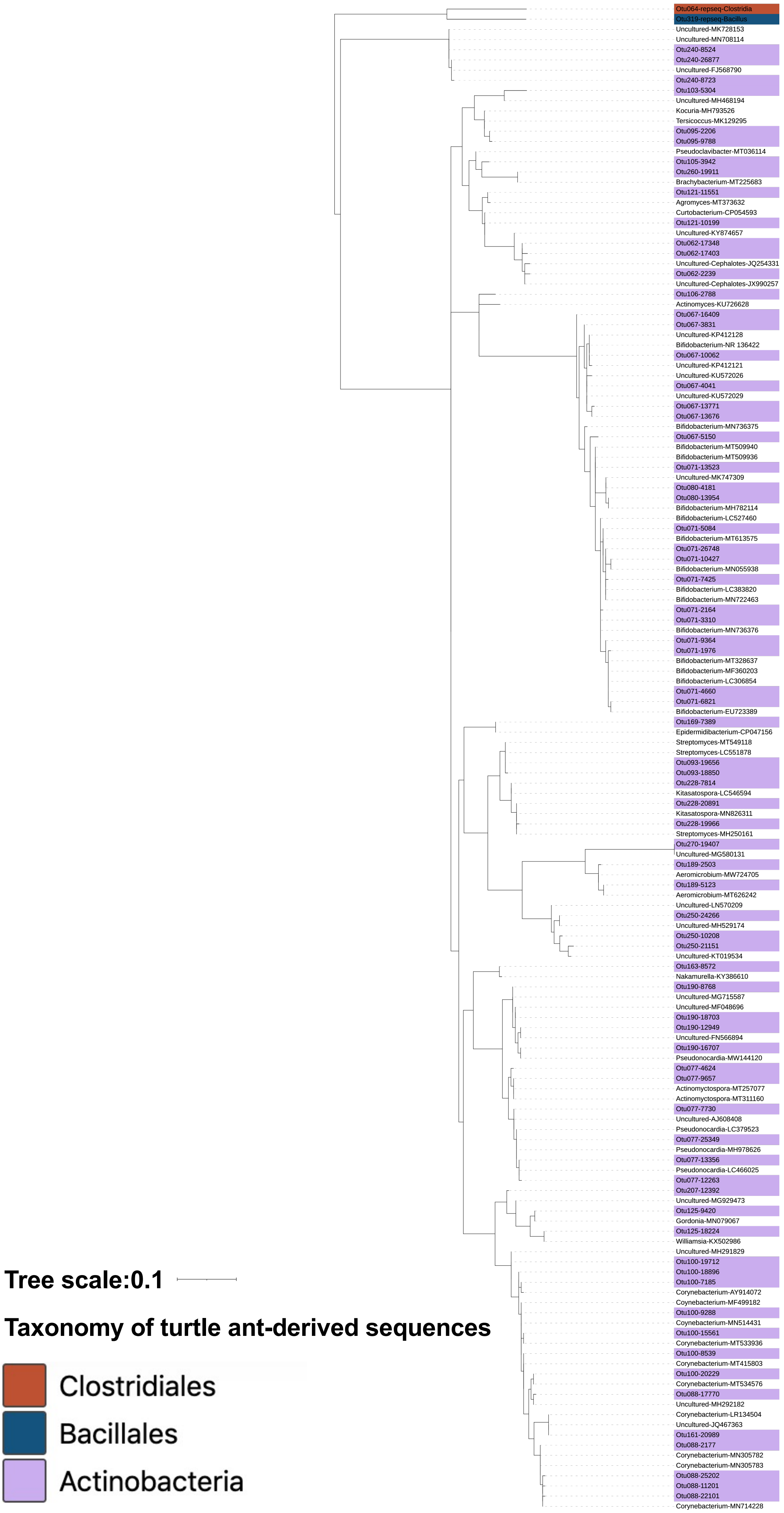
Otu064-repseq-Clostridia
Otu319-repseq-Bacillus
Uncultured-MK728153
Uncultured-MN708114
Otu240-8524
Otu240-26877
Uncultured-FJ568790
Otu240-8723
Otu103-5304
Uncultured-MH468194
Kocuria-MH793526
Tersicoccus-MK129295
Otu095-2206
Otu095-9788
Pseudoclavibacter-MT036114
Otu105-3942
Otu260-19911
Brachybacterium-MT225683
Otu121-11551
Agromyces-MT373632
Curtobacterium-CP054593
Otu121-10199
Uncultured-KY874657
Otu062-17348
Otu062-17403
Uncultured-Cephalotes-JQ254331
Otu062-2239
Uncultured-Cephalotes-JX990257
Otu106-2788
Actinomyces-KU726628
Otu067-16409
Otu067-3831
Uncultured-KP412128
Bifidobacterium-NR 136422
Otu067-10062
Uncultured-KP412121
Uncultured-KU572026
Otu067-4041
Uncultured-KU572029
Otu067-13771
Otu067-13676
Bifidobacterium-MN736375
Otu067-5150
Bifidobacterium-MT509940
Bifidobacterium-MT509936
Otu071-13523
Uncultured-MK747309
Otu080-4181
Otu080-13954
Bifidobacterium-MH782114
Bifidobacterium-LC527460
Otu071-5084
Bifidobacterium-MT613575
Otu071-26748
Otu071-10427
Bifidobacterium-MN055938
Otu071-7425
Bifidobacterium-LC383820
Bifidobacterium-MN722463
Otu071-2164
Otu071-3310
Bifidobacterium-MN736376
Otu071-9364
Otu071-1976
Bifidobacterium-MT328637
Bifidobacterium-MF360203
Bifidobacterium-LC306854
Otu071-4660
Otu071-6821
Bifidobacterium-EU723389
Otu169-7389
Epidermidibacterium-CP047156
Streptomyces-MT549118
Streptomyces-LC551878
Otu093-19656
Otu093-18850
Otu228-7814
Kitasatospora-LC546594
Otu228-20891
Kitasatospora-MN826311
Otu228-19966
Streptomyces-MH250161
Otu270-19407
Uncultured-MG580131
Otu189-2503
Aeromicrobium-MW724705
Otu189-5123
Aeromicrobium-MT626242
Uncultured-LN570209
Otu250-24266
Uncultured-MH529174
Otu250-10208
Otu250-21151
Uncultured-KT019534
Otu163-8572
Nakamurella-KY386610
Otu190-8768
Uncultured-MG715587
Uncultured-MF048696
Otu190-18703
Otu190-12949
Uncultured-FN566894
Otu190-16707
Pseudonocardia-MW144120
Otu077-4624
Otu077-9657
Actinomyctospora-MT257077
Actinomyctospora-MT311160
Otu077-7730
Uncultured-AJ608408
Pseudonocardia-LC379523
Otu077-25349
Pseudonocardia-MH978626
Otu077-13356
Pseudonocardia-LC466025
Otu077-12263
Otu207-12392
Uncultured-MG929473
Otu125-9420
Gordonia-MN079067
Otu125-18224
Williamsia-KX502986
Uncultured-MH291829
Otu100-19712
Otu100-18896
Otu100-7185
Corynebacterium-AY914072
Corynebacterium-MF499182
Otu100-9288
Corynebacterium-MN514431
Otu100-15561
Corynebacterium-MT533936
Otu100-8539
Corynebacterium-MT415803
Otu100-20229
Corynebacterium-MT534576
Otu088-17770
Uncultured-MH292182
Corynebacterium-LR134504
Uncultured-JQ467363
Otu161-20989
Otu088-2177
Corynebacterium-MN305782
Corynebacterium-MN305783
Otu088-25202
Otu088-11201
Otu088-22101
Corynebacterium-MN714228
Tree scale:0.1
Taxonomy of turtle ant-derived sequences
Clostridiales
Bacillales
Actinobacteria

**Supplementary Figure 6: Maximum likelihood phylogenetic analysis of the Actinobacteria (Analysis #4).** For this analysis (#4) we gathered all unique 16S rRNA sequences (n=73) coming from the 25 OTUs with unambiguous classifications to the phylum Actinobacteria. Top BLASTn hits were obtained for each unique sequence (vs. NCBI's nr database). If redundant with a prior top BLAST hit, we moved down the list to choose the most highly related, non-redundant match (as done for Analysis #s 5-9). After sequence alignment, we applied a Maximum Likelihood framework in Seaview, with the PhyML method and 100 bootstrap replicates. Phylogenies were visualized using the interactive Tree of Life website, and were inspected for monophyletic, cephalotine ant-specific clades showing distributions across multiple *Cephalotes* and/or *Procryptocerus* ant species.

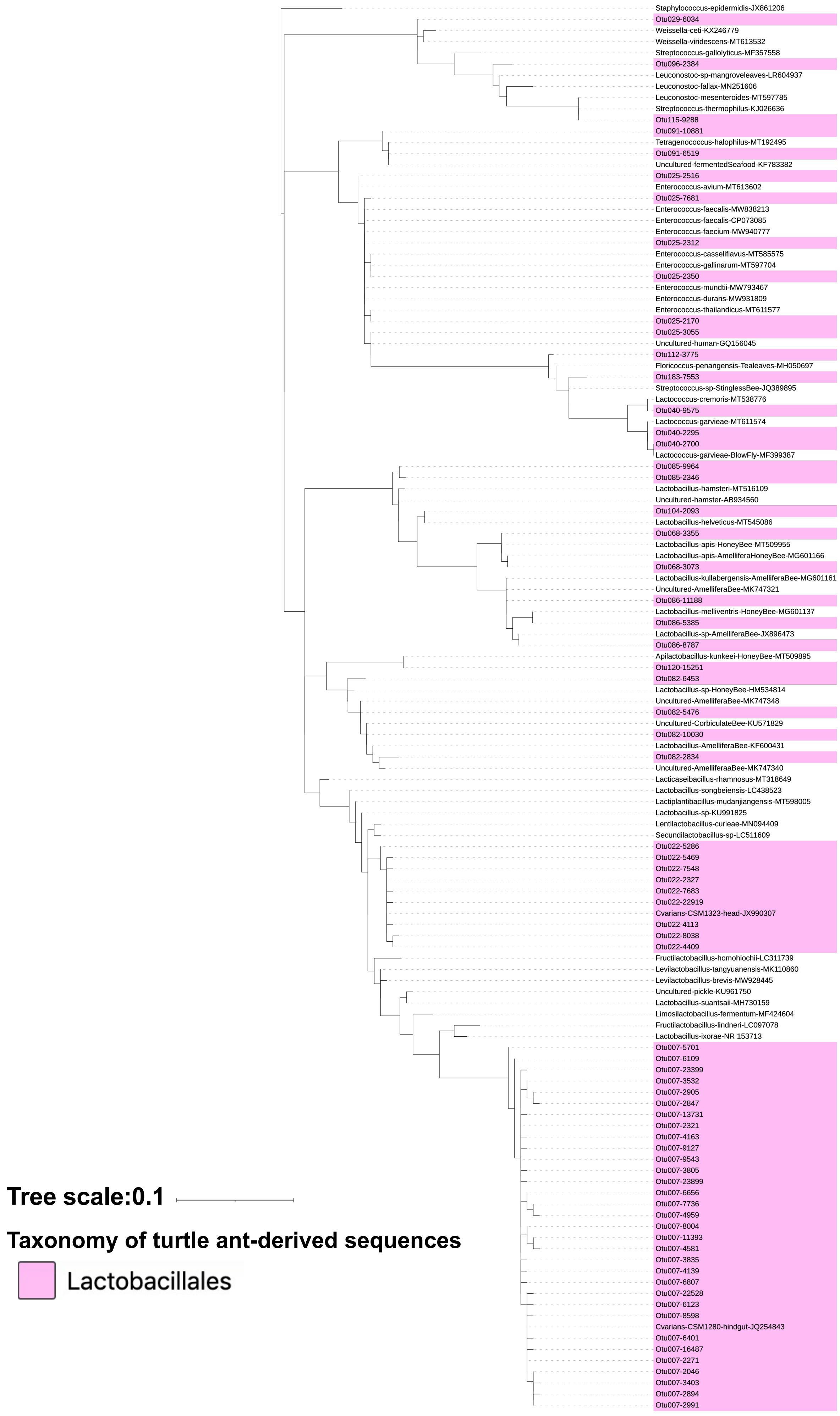

Staphylococcus-epidermidis-JX861206
Otu029-6034
Weissella-ceti-KX246779
Weissella-viridescens-MT613532
Streptococcus-gallolyticus-MF357558
Otu096-2384
Leuconostoc-sp-mangroveleaves-LR604937
Leuconostoc-fallax-MN251606
Leuconostoc-mesenteroides-MT597785
Streptococcus-thermophilus-KJ026636
Otu115-9288
Otu091-10881
Tetragenococcus-halophilus-MT192495
Otu091-6519
Uncultured-fermentedSeafood-KF783382
Otu025-2516
Enterococcus-avium-MT613602
Otu025-7681
Enterococcus-faecalis-MW838213
Enterococcus-faecalis-CP073085
Enterococcus-faecium-MW940777
Otu025-2312
Enterococcus-casseliflavus-MT585575
Enterococcus-gallinarum-MT597704
Otu025-2350
Enterococcus-mundtii-MW793467
Enterococcus-durans-MW931809
Enterococcus-thailandicus-MT611577
Otu025-2170
Otu025-3055
Uncultured-human-GQ156045
Otu112-3775
Floricoccus-penangensis-Tealeaves-MH050697
Otu183-7553
Streptococcus-sp-StinglessBee-JQ389895
Lactococcus-cremoris-MT538776
Otu040-9575
Lactococcus-garvieae-MT611574
Otu040-2295
Otu040-2700
Lactococcus-garvieae-BlowFly-MF399387
Otu085-9964
Otu085-2346
Lactobacillus-hamsteri-MT516109
Uncultured-hamster-AB934560
Otu104-2093
Lactobacillus-helveticus-MT545086
Otu068-3355
Lactobacillus-apis-HoneyBee-MT509955
Lactobacillus-apis-AmelliferaHoneyBee-MG601166
Otu068-3073
Lactobacillus-kullabergensis-AmelliferaBee-MG601161
Uncultured-AmelliferaBee-MK747321
Otu086-11188
Lactobacillus-melliventris-HoneyBee-MG601137
Otu086-5385
Lactobacillus-sp-AmelliferaBee-JX896473
Otu086-8787
Apilactobacillus-kunkeei-HoneyBee-MT509895
Otu120-15251
Otu082-6453
Lactobacillus-sp-HoneyBee-HM534814
Uncultured-AmelliferaBee-MK747348
Otu082-5476
Uncultured-CorbiculateBee-KU571829
Otu082-10030
Lactobacillus-AmelliferaBee-KF600431
Otu082-2834
Uncultured-AmelliferaaBee-MK747340
Lacticaseibacillus-rhamnosus-MT318649
Lactobacillus-songbeiensis-LC438523
Lactiplantibacillus-mudanjiangensis-MT598005
Lactobacillus-sp-KU991825
Lentilactobacillus-curieae-MN094409
Secundilactobacillus-sp-LC511609
Otu022-5286
Otu022-5469
Otu022-7548
Otu022-2327
Otu022-7683
Otu022-22919
Cvarians-CSM1323-head-JX990307
Otu022-4113
Otu022-8038
Otu022-4409
Fructilactobacillus-homohiochii-LC311739
Levilactobacillus-tangyuanensis-MK110860
Levilactobacillus-brevis-MW928445
Uncultured-pickle-KU961750
Lactobacillus-suantsaii-MH730159
Limosilactobacillus-fermentum-MF424604
Fructilactobacillus-lindneri-LC097078
Lactobacillus-ixorae-NR 153713
Otu007-5701
Otu007-6109
Otu007-23399
Otu007-3532
Otu007-2905
Otu007-2847
Otu007-13731
Otu007-2321
Otu007-4163
Otu007-9127
Otu007-9543
Otu007-3805
Otu007-23899
Otu007-6656
Otu007-7736
Otu007-4959
Otu007-8004
Otu007-11393
Otu007-4581
Otu007-3835
Otu007-4139
Otu007-6807
Otu007-22528
Otu007-6123
Otu007-8598
Cvarians-CSM1280-hindgut-JQ254843
Otu007-6401
Otu007-16487
Otu007-2271
Otu007-2046
Otu007-3403
Otu007-2894
Otu007-2991
Tree scale:0.1
Taxonomy of turtle ant-derived sequences
Lactobacillales

**Supplementary Figure 7: Maximum likelihood phylogenetic analysis of the Lactobacillales (Analysis #5).** For this analysis we broadened our focus to encompass less abundant Lactobacillales sequences not part of initial phylogenetic inference for this order (Analysis #3). Included were all unique 16S rRNA sequences from the order with relative abundance >0.002994 in at least two sequence libraries or >0.05 in one library. Top BLASTn hits (vs. NCBI's nr database) were obtained for each unique sequence, and if redundant with a prior top BLAST hit, we moved down the list to choose the most highly related, non-redundant match. After sequence alignment, we applied a Maximum Likelihood framework in Seaview, with the PhyML method and 100 bootstrap replicates. Phylogenies were visualized using the interactive Tree of Life website, and were inspected for monophyletic, cephalotine ant-specific clades showing distributions across multiple *Cephalotes* and/or *Procryptocerus* ant species.

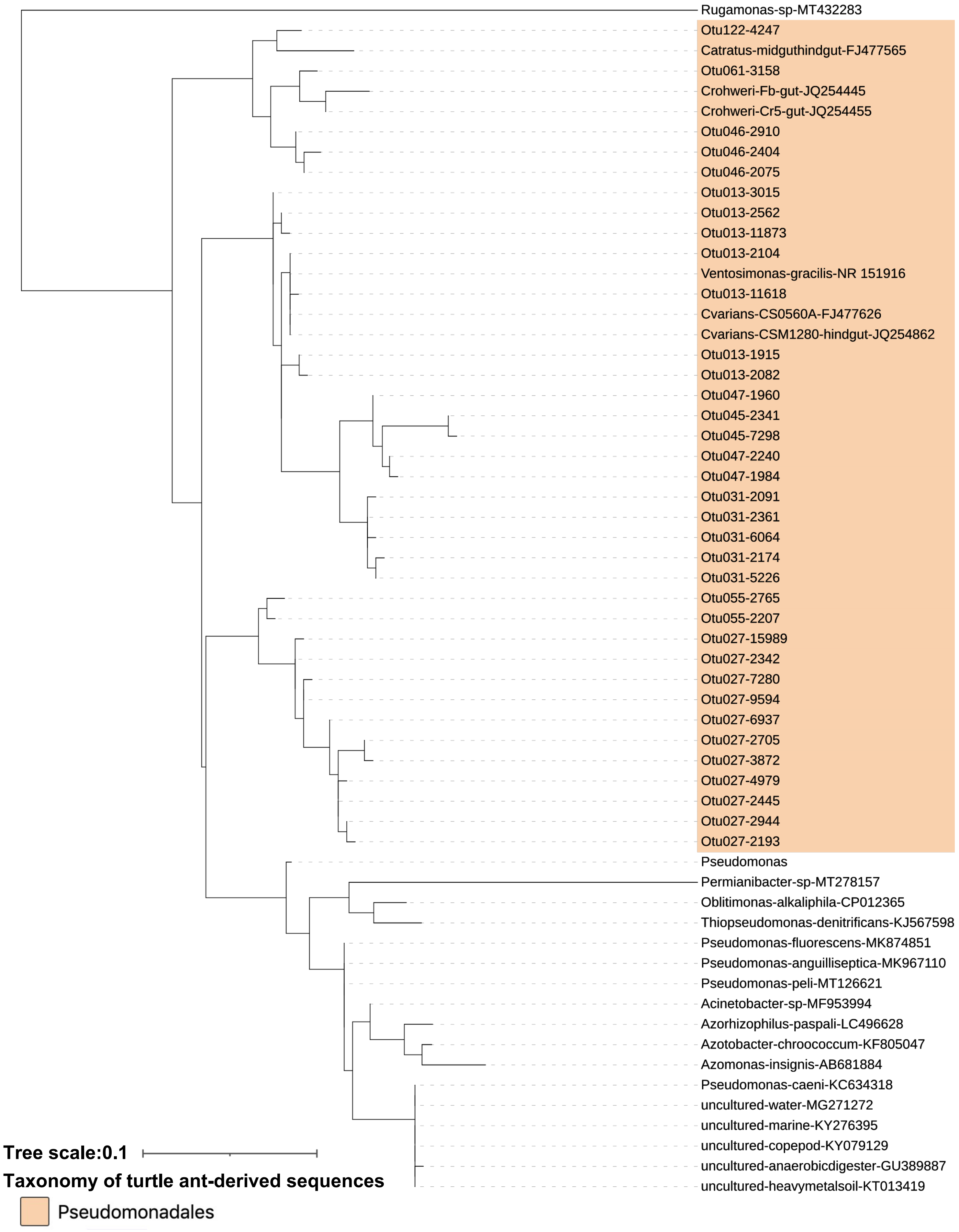
Rugamonas-sp-MT432283
Otu122-4247
Catratus-midguthindgut-FJ477565
Otu061-3158
Crohweri-Fb-gut-JQ254445
Crohweri-Cr5-gut-JQ254455
Otu046-2910
Otu046-2404
Otu046-2075
Otu013-3015
Otu013-2562
Otu013-11873
Otu013-2104
Ventosimonas-gracilis-NR 151916
Otu013-11618
Cvarians-CS0560A-FJ477626
Cvarians-CSM1280-hindgut-JQ254862
Otu013-1915
Otu013-2082
Otu047-1960
Otu045-2341
Otu045-7298
Otu047-2240
Otu047-1984
Otu031-2091
Otu031-2361
Otu031-6064
Otu031-2174
Otu031-5226
Otu055-2765
Otu055-2207
Otu027-15989
Otu027-2342
Otu027-7280
Otu027-9594
Otu027-6937
Otu027-2705
Otu027-3872
Otu027-4979
Otu027-2445
Otu027-2944
Otu027-2193
Pseudomonas
Permianibacter-sp-MT278157
Oblitimonas-alkaliphila-CP012365
Thiopseudomonas-denitrificans-KJ567598
Pseudomonas-fluorescens-MK874851
Pseudomonas-anguilliseptica-MK967110
Pseudomonas-peli-MT126621
Acinetobacter-sp-MF953994
Azorhizophilus-paspali-LC496628
Azotobacter-chroococcum-KF805047
Azomonas-insignis-AB681884
Pseudomonas-caeni-KC634318
uncultured-water-MG271272
uncultured-marine-KY276395
uncultured-copepod-KY079129
uncultured-anaerobicdigester-GU389887
uncultured-heavymetalsoil-KT013419
Tree scale:0.1
Taxonomy of turtle ant-derived sequences
Pseudomonadales

**Supplementary Figure 8: Maximum likelihood phylogenetic analysis of ambiguously classifying Pseudomonadales OTU046 (Analysis #6).** To ascertain whether OTU046 clustered with cephalotine ant-specialized bacteria from the Pseudomonadales, we first selected all unique 16S rRNA sequences from this OTU with relative abundance >0.002994 in at least two sequence libraries, or >0.05 in one library. With these sequences as queries, we performed BLASTn searches against the NCBI nr database, and against the genera from the Pseudomonadaceae family found to be highly related to OTU046 based on initial BLAST results. In addition to our inclusion of top BLASTn hits, our phylogenetic analysis also included unique sequences from OTUs more confidently assigning as cephalotine-specialized Pseudomonadales through prior BLASTn searches. After sequence alignment, we applied a Maximum Likelihood framework in Seaview, with the PhyML method and 100 bootstrap replicates. Phylogenies were visualized using the interactive Tree of Life website, and were inspected for monophyletic, cephalotine ant-specific clades showing distributions across multiple *Cephalotes* and/or *Procryptocerus* ant species.

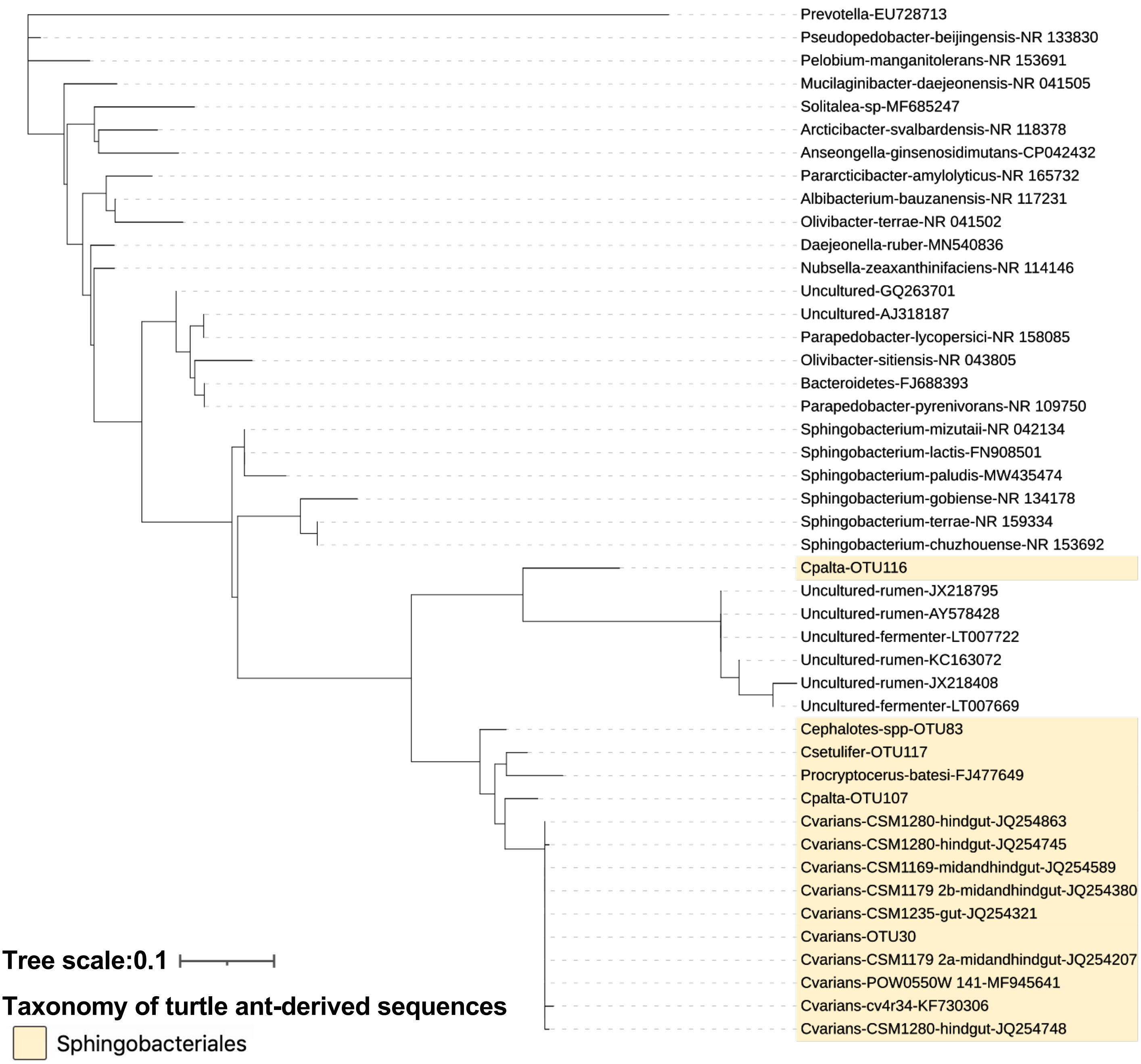

Prevotella-EU728713
Pseudopedobacter-beijingensis-NR 133830
Pelobium-manganitolerans-NR 153691
Mucilaginibacter-daejeonensis-NR 041505
Solitalea-sp-MF685247
Arcticibacter-svalbardensis-NR 118378
Anseongella-ginsenosidimutans-CP042432
Pararcticibacter-amylolyticus-NR 165732
Albibacterium-bauzanensis-NR 117231
Olivibacter-terrae-NR 041502
Daejeonella-ruber-MN540836
Nubsella-zeaxanthinifaciens-NR 114146
Uncultured-GQ263701
Uncultured-AJ318187
Parapedobacter-lycopersici-NR 158085
Olivibacter-sitiensis-NR 043805
Bacteroidetes-FJ688393
Parapedobacter-pyrenivorans-NR 109750
Sphingobacterium-mizutaii-NR 042134
Sphingobacterium-lactis-FN908501
Sphingobacterium-paludis-MW435474
Sphingobacterium-gobiense-NR 134178
Sphingobacterium-terrae-NR 159334
Sphingobacterium-chuzhouense-NR 153692
Cpalta-OTU116
Uncultured-rumen-JX218795
Uncultured-rumen-AY578428
Uncultured-fermenter-LT007722
Uncultured-rumen-KC163072
Uncultured-rumen-JX218408
Uncultured-fermenter-LT007669
Cephalotes-spp-OTU83
Csetulifer-OTU117
Procryptocerus-batesi-FJ477649
Cpalta-OTU107
Cvarians-CSM1280-hindgut-JQ254863
Cvarians-CSM1280-hindgut-JQ254745
Cvarians-CSM1169-midandhindgut-JQ254589
Cvarians-CSM1179 2b-midandhindgut-JQ254380
Cvarians-CSM1235-gut-JQ254321
Cvarians-OTU30
Cvarians-CSM1179 2a-midandhindgut-JQ254207
Cvarians-POW0550W 141-MF945641
Cvarians-cv4r34-KF730306
Cvarians-CSM1280-hindgut-JQ254748
Tree scale:0.1
Taxonomy of turtle ant-derived sequences
Sphingobacteriales

**Supplementary Figure 9: Maximum likelihood phylogenetic analysis of ambiguously classifying Sphingobacteriales OTU116 (Analysis #7).** To characterize the evolution of an ambiguously classifying Sphingobacteriales OTU with a top BLASTn hit showing only ~85% sequence identity to a specialized Sphingobacteriales symbiont from *Cephalotes* ants, we performed additional phylogenetic analyses. First, we selected all unique 16S rRNA sequences from this OTU showing relative abundance >0.002994 in at least two sequence libraries, or >0.05 in one library – this equated to exactly n=1 sequence. We used this sequence for a BLASTn search against the NCBI nr database. We also included 1) related OTUs from the cephalotine-associated Sphingobacteriales identified in the present study (OTUs 038, 083, and 107); 2) top BLASTn hits to these 3 putatively related, cephalotine-specific Sphingobacteriales OTUs; and 3) relatives identified via BLASTn results targeting a range of genera and families in the Sphingobacteriales. After sequence alignment, we applied a Maximum Likelihood framework in Seaview, with the PhyML method and 100 bootstrap replicates. Phylogenies were visualized using the interactive Tree of Life website, and were inspected for monophyletic, cephalotine ant-specific clades showing distributions across multiple *Cephalotes* and/or *Procryptocerus* ant species.

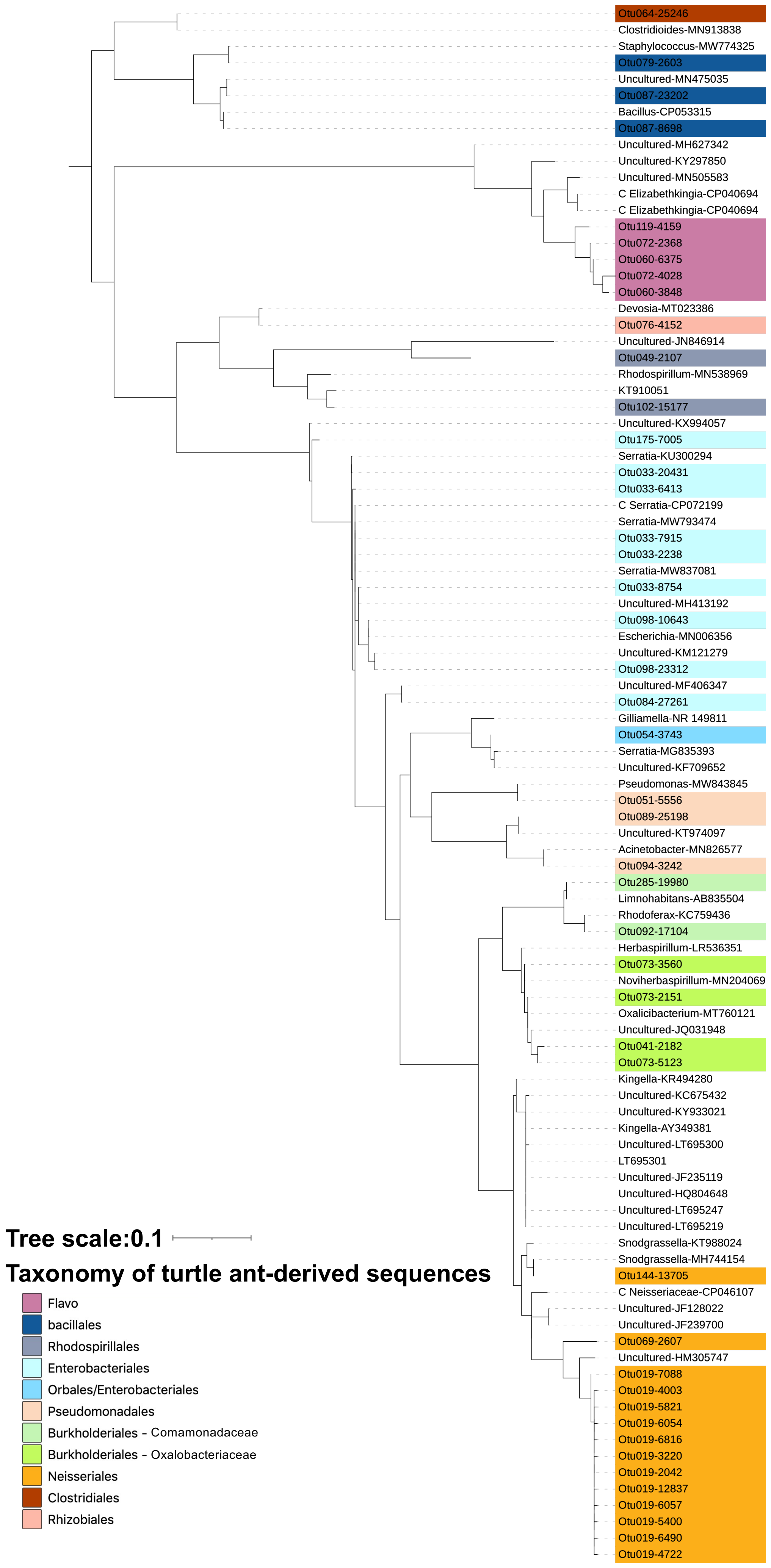
Otu064-25246
Clostridioides-MN913838
Staphylococcus-MW774325
Otu079-2603
Uncultured-MN475035
Otu087-23202
Bacillus-CP053315
Otu087-8698
Uncultured-MH627342
Uncultured-KY297850
Uncultured-MN505583
C Elizabethkingia-CP040694
C Elizabethkingia-CP040694
Otu119-4159
Otu072-2368
Otu060-6375
Otu072-4028
Otu060-3848
Devosia-MT023386
Otu076-4152
Uncultured-JN846914
Otu049-2107
Rhodospirillum-MN538969
KT910051
Otu102-15177
Uncultured-KX994057
Otu175-7005
Serratia-KU300294
Otu033-20431
Otu033-6413
C Serratia-CP072199
Serratia-MW793474
Otu033-7915
Otu033-2238
Serratia-MW837081
Otu033-8754
Uncultured-MH413192
Otu098-10643
Escherichia-MN006356
Uncultured-KM121279
Otu098-23312
Uncultured-MF406347
Otu084-27261
Gilliamella-NR 149811
Otu054-3743
Serratia-MG835393
Uncultured-KF709652
Pseudomonas-MW843845
Otu051-5556
Otu089-25198
Uncultured-KT974097
Acinetobacter-MN826577
Otu094-3242
Otu285-19980
Limnohabitans-AB835504
Rhodoferax-KC759436
Otu092-17104
Herbaspirillum-LR536351
Otu073-3560
Noviherbaspirillum-MN204069
Otu073-2151
Oxalicibacterium-MT760121
Uncultured-JQ031948
Otu041-2182
Otu073-5123
Kingella-KR494280
Uncultured-KC675432
Uncultured-KY933021
Kingella-AY349381
Uncultured-LT695300
LT695301
Uncultured-JF235119
Uncultured-HQ804648
Uncultured-LT695247
Uncultured-LT695219
Snodgrassella-KT988024
Snodgrassella-MH744154
Otu144-13705
C Neisseriaceae-CP046107
Uncultured-JF128022
Uncultured-JF239700
Otu069-2607
Uncultured-HM305747
Otu019-7088
Otu019-4003
Otu019-5821
Otu019-6054
Otu019-6816
Otu019-3220
Otu019-2042
Otu019-12837
Otu019-6057
Otu019-5400
Otu019-6490
Otu019-4722
Tree scale:0.1
Taxonomy of turtle ant-derived sequences
Flavo
bacillales
Rhodospirillales
Enterobacteriales
Orbales/Enterobacteriales
Pseudomonadales
Burkholderiales - Comamonadaceae
Burkholderiales - Oxalobacteriaceae
Neisseriales
Clostridiales
Rhizobiales

**Supplementaruy Figure S10: Maximum likelihood phylogenetic analysis of other sequences with high abundance, and their relatives (Analysis #8).** For this final phylogenetic analysis aimed at identifying specialized bacterial symbionts of cephalotine ants, we focused on subset of the 154 OTUs exceeding the required abundance threshold in ant sequence libraries – i.e. the most abundant left-overs, not analyzed in Analysis #1-7 above, with OTU IDs falling between #s 001 –100. These sequences did not belong to an OTU assigning as a specialized core cephalotine bacterium via BLAST, and had not been studied extensively in previously published phylogenetic analyses on ant-associated bacteria (e.g. *Wolbachia*, Entomoplasmatales – Russell et al. 2009; Funaro et al. 2011). This amounted to 19 selected OTUs, hailing from 8 orders. To maximize our chances of finding cephalotine-specific linages we analyzed unique sequences from all OTUs in these 8 orders (i.e. not just the 19 selected OTUs) if they exhibited >0.002994 relative abundance in at least 2 *Cephalotes* sequence libraries, or if they were present in >0.05 relative abundance in at least one library. Top BLASTn hits were downloaded for each of these sequences. After sequence alignment, we applied a Maximum Likelihood framework in Seaview, with the PhyML method and 100 bootstrap replicates. Phylogenies were visualized using the interactive Tree of Life website, and were inspected for monophyletic, cephalotine ant-specific clades showing distributions across multiple *Cephalotes* and/or *Procryptocerus* ant species.

| Geography | Colony ID | Species | Caste | % of microbiome comprised by OTUs from specialized *Cephalotes* clades | % of specialized (host-specific) microbes not including Rhizobiales | % of microbiome comprised by OTUs from specialized, adult-enriched *Cephalotes* clades | % of microbiome comprised by OTUs from specialized, adult-enriched *Cephalotes* clades not including Rhizobiales | Notes on abberrations |
|---|---|---|---|---|---|---|---|---|
| Br | C15-20 | *Cephalotes targionii* | S | 0.9864 | 0.6129 | 0.9864 | 0.6129 | |
| Br | | | W | 0.9901 | 0.6172 | 0.9901 | 0.6172 | |
| Br | C15-21 | *Cephalotes grandinosus* | S | 0.9744 | 0.8194 | 0.9744 | 0.8194 | |
| Br | | | W | 0.9818 | 0.7312 | 0.9818 | 0.7312 | |
| Br | C15-44 | | S | 0.9960 | 0.7883 | 0.9960 | 0.7883 | |
| Br | | | W | 0.9915 | 0.4738 | 0.9908 | 0.4730 | |
| Br | C15-28 | *Cephalotes maculatus* | S | 0.9792 | 0.5734 | 0.9792 | 0.5734 | |
| Br | | | W | 0.9964 | 0.5282 | 0.9964 | 0.5282 | |
| FLK | YH083 | *Cephalotes varians* | S | 0.9900 | 0.8113 | 0.9900 | 0.8113 | |
| FLK | | | W | 0.9712 | 0.8377 | 0.9710 | 0.8376 | |
| FLK | YH091 | | S | 0.9878 | 0.8177 | 0.9878 | 0.8177 | |
| FLK | | | W | 0.9692 | 0.8493 | 0.9684 | 0.8486 | |
| FLK | YH092 | | S | 0.9911 | 0.8733 | 0.9911 | 0.8733 | |
| FLK | | | W | 0.9890 | 0.8430 | 0.9890 | 0.8430 | |
| FLK | YH100 | | W | 0.9904 | 0.8174 | 0.9904 | 0.8174 | |
| FLK | | | AQ | 0.9901 | 0.8751 | 0.9901 | 0.8751 | |
| Br | C15-25 | *Cephalotes depressus* | S | 0.9300 | 0.7984 | 0.6151 | 0.4836 | Neisseriales & Lactobacillales abundance |
| Br | | | W | 0.9963 | 0.7828 | 0.9963 | 0.7828 | |
| CR | SRCset1 | *Cephalotes setulifer* | S | 0.9799 | 0.8394 | 0.9799 | 0.8394 | |
| CR | | | W | 0.9935 | 0.8337 | 0.9935 | 0.8337 | |
| CR | | | AQ | 0.9778 | 0.8968 | 0.9778 | 0.8968 | |
| CR | SRCset2 | | S | 0.9889 | 0.9342 | 0.9889 | 0.9342 | |
| CR | | | W | 0.9926 | 0.8704 | 0.9926 | 0.8704 | |
| CR | | | AQ | 0.9945 | 0.8989 | 0.9945 | 0.8989 | |
| Br | C15-40 | *Cephalotes pusillus* | S | 0.9729 | 0.4315 | 0.9234 | 0.3820 | Lactobacillales abundance |
| Br | | | W | 0.9676 | 0.7018 | 0.9533 | 0.6874 | |
| CR | SRCmin1 | *Cephalotes minutus* | S | 0.9968 | 0.8350 | 0.9968 | 0.8350 | |
| CR | | | W | 0.9930 | 0.8028 | 0.9930 | 0.8028 | |
| MX - ECBR | JS-C-235 | *Cephalotes texanus* | S | 0.9891 | 0.7970 | 0.6268 | 0.4347 | Lactobacillales abundance |
| MX - ECBR | | | W | 0.9854 | 0.8207 | 0.6571 | 0.4924 | Lactobacillales abundance |
| MX - ECBR | JS-C-220 | *Cephalotes* aff. *bimaculatus* | S | 0.4050 | 0.2111 | 0.4042 | 0.2103 | Neisseriales abundance |
| MX - ECBR | | | W | 0.5474 | 0.2467 | 0.5474 | 0.2467 | Neisseriales abundance |
| MX - SdC | JS-C-211 | *Cephalotes* aff. *wheeleri* | S | 0.9782 | 0.6881 | 0.9782 | 0.6881 | |
| MX - SdC | | | W | 0.9653 | 0.6516 | 0.9653 | 0.6516 | |
| MX - SdC | JS-3-203 | *Cephalotes goniodontus* | AQ | 0.5144 | 0.3521 | 0.5144 | 0.3521 | Wolbachia abundance |
| MX - SdC | | | W | 0.9376 | 0.7615 | 0.9376 | 0.7615 | |
| DR | C16-08 | *Cephalotes unimaculatus* | W | 0.9507 | 0.7802 | 0.9496 | 0.7791 | |
| Br | C15-44 | *Cephalotes grandinosus* | Queen | 0.9971 | 0.0963 | 0.9971 | 0.0963 | |
| Br | C15-28 | *Cephalotes maculatus* | | 0.4677 | 0.0000 | 0.4677 | 0.0000 | Wolbachia abundance |
| FLK | YH091 | *Cephalotes varians* | | 0.9981 | 0.0000 | 0.9981 | 0.0000 | |
| Br | C15-25 | *Cephalotes depressus* | | 0.4694 | 0.4506 | 0.0188 | 0.0000 | Neisseriales abundance |
| Br | C15-40 | *Cephalotes pusillus* | | 0.3739 | 0.0000 | 0.3739 | 0.0000 | Wolbachia abundance |
| Br | C15-20 | *Cephalotes targionii* | Larva | 0.4657 | 0.1636 | 0.4657 | 0.1636 | Burkholderiales (Alcaligenaceae) abundance |
| Br | C15-21 | *Cephalotes grandinosus* | | 0.7326 | 0.0829 | 0.7326 | 0.0829 | Burkholderiales (Alcaligenaceae) abundance |
| Br | C15-44 | | | 0.8689 | 0.0745 | 0.8689 | 0.0745 | Burkholderiales (Alcaligenaceae) abundance |
| Br | C15-28 | *Cephalotes maculatus* | | 0.5347 | 0.0401 | 0.5347 | 0.0401 | |
| FLK | YH083 | *Cephalotes varians* | | 0.6759 | 0.3264 | 0.3544 | 0.0049 | |
| FLK | YH091 | | | 0.8500 | 0.1619 | 0.6968 | 0.0087 | |
| FLK | YH092 | | | 0.6998 | 0.0292 | 0.6955 | 0.0249 | |
| FLK | YH100 | | | 0.6375 | 0.2635 | 0.3780 | 0.0041 | |
| Br | C15-25 | *Cephalotes depressus* | | 0.7685 | 0.4076 | 0.3798 | 0.0190 | |
| CR | SRCset1 | *Cephalotes setulifer* | | 0.6522 | 0.0653 | 0.6522 | 0.0653 | |
| CR | SRCset2 | | | 0.8567 | 0.3782 | 0.8552 | 0.3766 | Burkholderia (Alcaligenaceae) & Xanthomonadales abundance |
| Br | C15-40 | *Cephalotes pusillus* | | 0.2441 | 0.0907 | 0.1708 | 0.0173 | Wolbachia abundance |
| CR | SRCmin1 | *Cephalotes minutus* | | 0.7136 | 0.0189 | 0.7025 | 0.0077 | |
| MX - ECBR | JS-C-235 | *Cephalotes texanus* | | 0.7164 | 0.5844 | 0.1550 | 0.0230 | |
| MX - SdC | JS-C-211 | *Cephalotes* aff. *wheeleri* | | 0.8071 | 0.0212 | 0.8071 | 0.0212 | |
| MX - SdC | JS-3-203 | *Cephalotes goniodontus* | | 0.5935 | 0.4901 | 0.1139 | 0.0105 | Wolbachia abundance |
| DR | C16-08 | *Cephalotes unimaculatus* | | 0.6630 | 0.3054 | 0.4319 | 0.0743 | Burkholderiales (Alcaligenaceae) abundance |

**Supplementary Figure S11: Proportions of microbiomes comprised by bacteria from cephalotine -specialist lineages.** Data focus on the 154 OTUs with >0.002994 in 2 or more sequence libraries or >0.05 in at least one. After BLASTn analyses and phylogenetics (Analysis #s 1-8), each OTU was classified as belonging to a lineage comprised of bacteria found only in *Cephalotes* or *Procryptocerus* ants, separated from other lineages by long branches; or, clustering with bacteria from other hosts or habitats, including free-living bacteria. Proportions of the microbiome for each sequence library were averaged, by caste and stage, separately for each *Cephalotes* ant colony. Averages were separated for pre- and non-reproductive ants (alate queens "AQ", vs. soldiers and workers, "S" and "W"), wingless (mature) queens, and larvae. Averages were shown after including or excluding Rhizobiales bacteria, which made up a large proportion of the specialized larval microbiome, and with vs. without the newly identified candidate Lactobacillales specialists from OTU007 and OTU002, also rare in adults and without clear transmission mechanisms enabling partner fidelity. Heatmap ranges from white to red, to show low to high average relative abundances. Notes on abberrations are provided for stages/castes from colonies exhibiting exceptions to general trends across other colonies. Colors used for geographic origins and host taxonomy are the same as those in Figure 3.

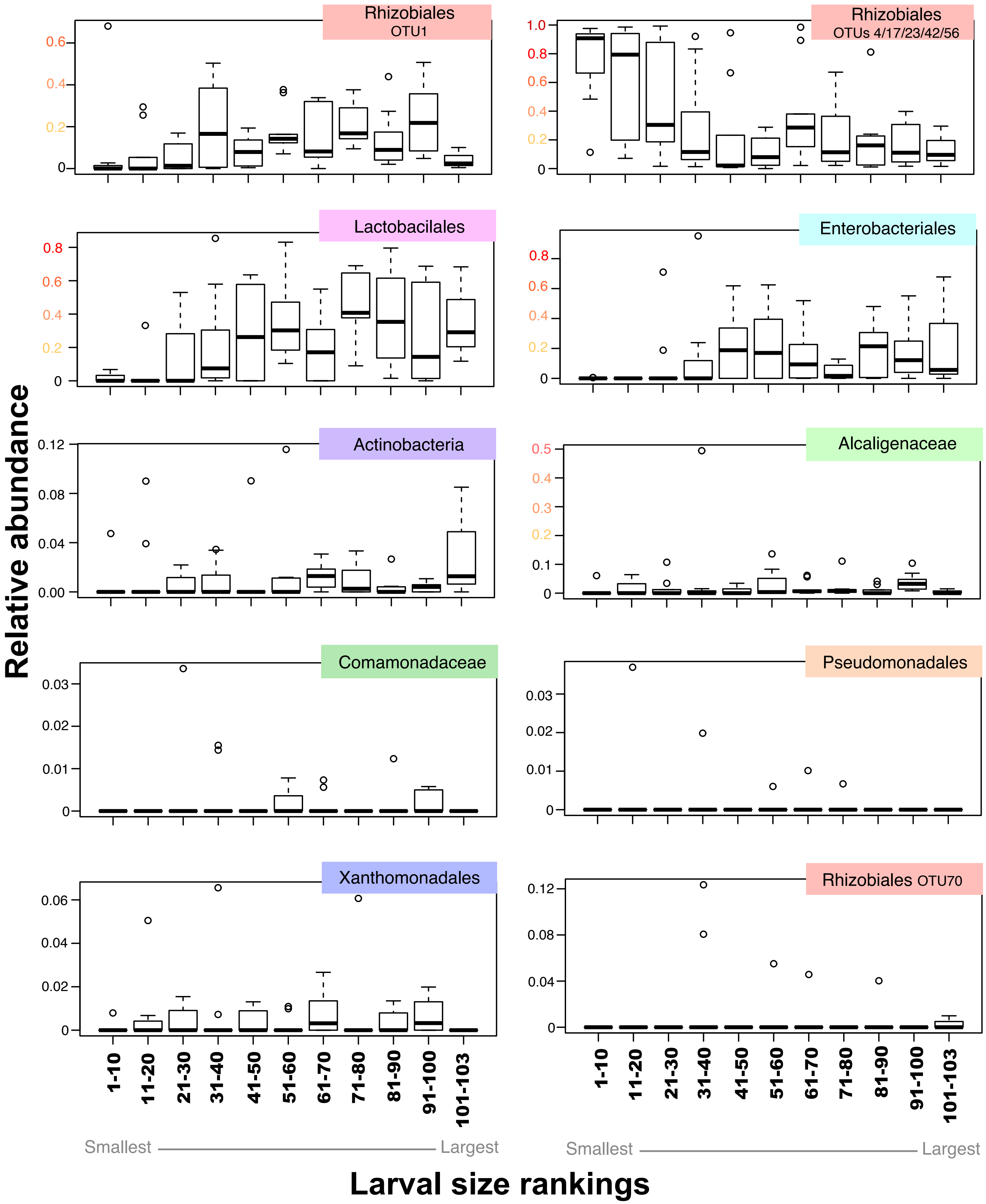
Rhizobiales OTU1
Rhizobiales OTUs 4/17/23/42/56
Lactobacilales
Enterobacteriales
Actinobacteria
Alcaligenaceae
Comamonadaceae
Pseudomonadales
Xanthomonadales
Rhizobiales OTU70
Relative abundance
1-10
11-20
21-30
31-40
41-50
51-60
61-70
71-80
81-90
91-100
101-103
Smallest
Largest
Larval size rankings

**Supplementary Figure 12: Bacterial abundance across larvae of varying age.** Size used here as a proxy for age, and all but the largest of these bins (n=3 larvae) contained 10 larvae of similar size. To simplify graphics and minimize the number of statistical tests we pooled relative abundances from a number of related 97% OTUs, including those from specialized core symbionts of the Alcaligenaceae (Burkholderiales) (OTUs 010/011/012/014/016/024), Comamonadaceae (Burkholderiales) (OTUs 020/021/032),and Xanthomonadales (OTUs 002/005/015). Individual values for the specialized Pseudomonadales (OTU013) were utilized. For specialized Rhizobiales we divided data into two groups – OTU001 vs. the summed total of OTUs 004/017/023/042/056 – based on patterns observed in our phylogenetic analyses.Beyond these specialist symbionts, we computed relative abundance sums of four common Enterobacteriales (OTUs 006/033/063/084) – all non-specialists – and, ten OTUs within the Lactobacillales (OTUs 007/022/025/029/040/068/082/085/096/104), containing a mixture of specialists (OTUs 007 & 022) and non-specialists. For two more larva-abundant taxa (OTU070 - non-specialized Rhizobiales; and OTU009 - Wolbachia - not shown here), we performed statistics on individual OTUs. Finally, we summed the six OTUs from the phylum Actinobacteria (OTUs 062/067/080/088/093/095) into a single total prior to statistics. While these OTUs were distributed across the Actinomycetales and Bifidobacteriales orders, this broader approach enabled us to focus on a group commonly known for its antimicrobial capacities, and to minimize the number of statistical tests on orders or OTUs with fairly sporadic occurrence. Details on statistics can be found in **Supplementary Table 21**. Y-values reveal relative abundances. Asterisks under each size class category, along the x-axis for a given panel indicate significant differences from the overall mean, in an ANOVA analysis. Red asterisks indicate a relative abundance value greater than the overall mean; blue asterisks indicate values lower than the overall mean. *: $0.05<p<0.1$; **: $0.01<p<0.05$; ***: $p<0.01$.

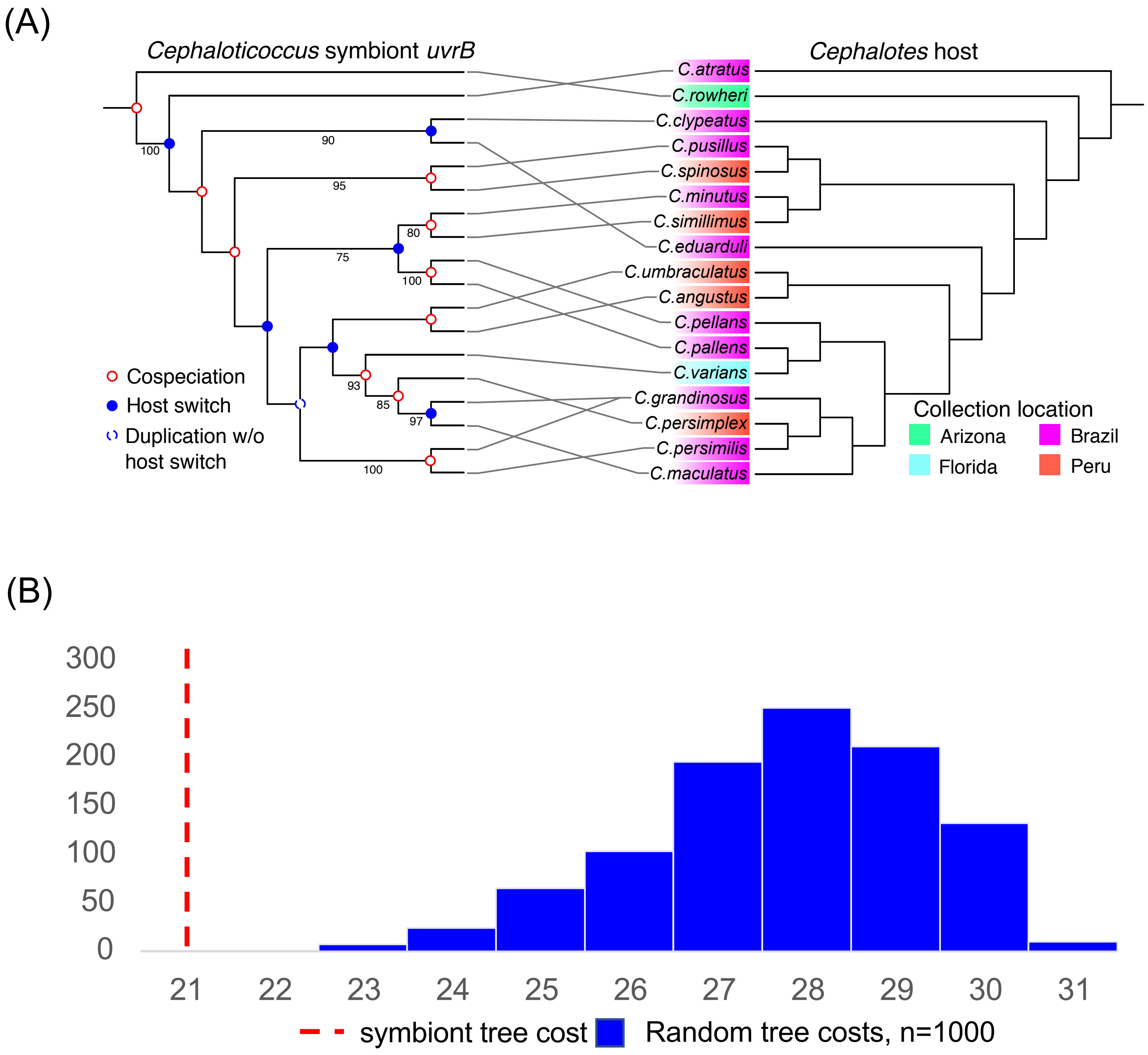


**Supplementary Figure 13: Cophylogeny analyses for *Cephaloticoccus* symbionts and *Cephalotes* ant hossts. (A**) Tanglegram of uvrB gene tree (from *Cephaloticoccus* symbionts extracted from metagenomes of Hu et al. 2018) vs. *Cephalotes* host tree, from Price et al. 2016. For the symbiont gene tree, will list all bootstrap values >70. Cospeciation and host switching events are indicated at each symbiont node and were inferred based on the top solution from Jane4. (**B**) Cost histogram from 1000 randomized tip mapping permutations in Jane4. A lower cost indicates fewer inferred host switching events, and therefore stronger phylogenetic congruence.
